# gLinDA: Global Differential Abundance Analysis of Microbiomes

**DOI:** 10.1101/2024.09.29.615668

**Authors:** Leon Fehse, Mohammad Tajabadi, Roman Martin, Hajo Holzmann, Dominik Heider

## Abstract

Microbiome composition plays a significant role in various diseases, including cancer, obesity, inflammatory bowel disease, and mental health disorders. Understanding the differences in microbial abundance between patients is key to uncovering the microbiome’s impact on these conditions. Differential abundance analysis (DAA) can detect significant changes in microbiomes at the taxa level base. However, since individuals have unique microbial fingerprints that can potentially identify them, microbiome data must be treated as sensitive. We introduce gLinDA, a global differential abundance analysis software offering a swarm learning approach to analyzing distributed datasets while maintaining result accuracy without sharing patients’ raw data.

## Introduction

Microbiomes are complex ecosystems composed of various microorganisms, such as bacteria, viruses, and fungi. These microorganisms play a crucial role in various biological processes, including human health, environmental balance, and agricultural productivity (Autenrieth, 2016). Microbiomes are as diverse as they are complex, existing in various natural reservoirs from the human gut to soil and water ecosystems (Sperlea et al., 2021), where they frequently interact with different chemical substances (Spänig et al., 2021).

Differential Abundance Analysis (DAA) is a key statistical method in microbiome research, used to compare the prevalence of microbial taxa across different groups, such as individuals with and without specific health conditions. This approach is essential for identifying taxa that show significant differences in abundance, either being more prevalent or scarce across the groups. These analyses are crucial for understanding the relationship between microbiome composition and health issues or environmental exposures. DAA employs statistical techniques to account for the variability and structure of microbiome data, ensuring accurate interpretation of observed differences. Conducting a thorough DAA on microbiome datasets presents challenges, including variability in read counts, the compositional nature of the data, and frequent zero-inflated distributions (Cappellato et al., 2022). To address these challenges, various software tools for DAA, such as LinDA (Zhou et al., 2022), ZicoSeq (Yang and Chen, 2022), MaAsLin2 (Mallick et al., 2021), and ZIBR (Chen and Li, 2016), have been developed. These tools differ in methodology, leading to variations in results, as shown in benchmark studies (Nearing et al., 2022; Yang and Chen, 2022). In their benchmark study from 2023, Yang et al. (Yang and Chen, 2023) identified LinDA as the best-performing method based on its overall performance.

DAA commonly requires the inclusion of covariates and must account for the zero-inflated nature of microbiome data. Various software tools have been developed to address these challenges using different methodologies. ZicoSeq, for instance, performs association tests by linearly regressing log-transformed taxa abundance values on covariates and employs a reference-based strategy to correct compositional effects, along with an empirical Bayes approach to handle zero inflation (Yang and Chen, 2022). LinDA also fits a linear regression model on log-transformed taxa abundances, accounting for covariates while addressing compositional and zero-inflation issues through a sample library size-based strategy (Zhou et al., 2022). Both ZicoSeq and LinDA utilize winsorization to manage outliers, with ZicoSeq applying permutation-based false discovery rate (FDR) control and LinDA using the Benjamini-Hochberg procedure. In contrast, ZIBR models longitudinal data through a two-part logistic-beta regression model, combining a logistic component and a beta distribution component (Chen and Li, 2016). MaAsLin2 similarly applies generalized linear models (GLMs), supporting linear regression, logistic regression, and zero-inflated models to handle microbiome data (Mallick et al., 2021). It conducts multivariate regression for each feature, using metadata and covariates as predictors.

To our knowledge, all existing methods for differential abundance analysis specialized for microbiome analyses require centralized data storage. While for one data source, it is reasonable, it becomes problematic for multi-centered approaches, raising concerns about data privacy and security. Generally, the law of large numbers applies, implying using more data for better statistical evidence. Sharing raw microbiome data is particularly sensitive, as individuals possess unique microbial “fingerprints” that could potentially be used for identification (Franzosa et al., 2015; Mayer et al., 2024). Consequently, microbiome data must be treated as highly sensitive. In recent years, new methodologies have emerged to address these privacy concerns. Federated learning, for example, is a well-established privacy-preserving approach that enables analysis across multiple sources without sharing raw data. Decentralized learning techniques (Tajabadi et al., 2024b), such as random forests (Hauschild et al., 2022), graph neural networks (Pfeifer et al., 2023), and genome-wide association studies (Nasirigerdeh et al., 2022), are already employed in other fields. In contrast to traditional federated learning, which relies on a centralized server, methods like swarm learning (Warnat-Herresthal et al., 2021) use peer-to-peer communication, eliminating dependency on a central server and allowing direct interaction between data owners (Tajabadi et al., 2024a).

To overcome the limitations related to privacy, robustness, and accuracy in existing methods, we propose gLinDA as a decentralized learning software for differential abundance analysis. This tool offers a privacy-preserving and flexible solution for microbiome analysis based on linear models. Utilizing a swarm learning approach, gLinDA ensures that data remains locally stored and processed, safeguarding privacy while enabling accurate and comprehensive analysis. Written in Python, gLinDA includes a graphical user interface (GUI), making it highly user-friendly for researchers. Its versatility is reflected in its availability as both a GUI and a command-line tool, as well as its open-source Python implementation, allowing it to be integrated into scripts or used as a standalone application. The statistical implementation is based on the benchmark-leading LinDA (Yang and Chen, 2023; Zhou et al., 2022), originally written in R, delivering comparable performance in a decentralized setting and identical performance when run locally on a complete dataset.

## Results

The primary function of gLinDA is to identify significant differential abundances of taxa or, more precisely, operational taxonomic units (OTUs) based on provided metadata, such as clinical outcomes or other covariates. To establish a suitable benchmark, we compared the standalone LinDA with gLinDA across various scenarios involving two, three, four, and eight peers. Two datasets were showcased in this manuscript for all scenarios: the simulated S-5000 dataset (Yang and Chen, 2023) and the real-world OB-Goodrich dataset (Goodrich et al., 2014). Additionally, benchmarks were conducted on a total of eight datasets, as it is listed in Table 2 and detailed in the supplementary information.

Furthermore, we included meta-analysis in our benchmarks to illustrate the advantages of the decentralized learning approach with gLinDA over simply averaging LinDA summaries, such as p-values, from individual peers. For the meta-analysis benchmarks, we combined p-values from each peer using a weighted averaging method. This comparison highlights why sharing model parameters, rather than just summary statistics, preserves more information and improves analytical power.

A unique challenge in decentralized analyses with gLinDA, which does not arise in locally run LinDA, is handling taxa or OTUs that are not consistently present across all peers. To address this, gLinDA offers two modes for combining results from individual peers: union and intersection. The intersection mode excludes OTUs that are not represented in every peer, ensuring consistency across datasets, whereas the union mode includes as many OTUs as possible, even if they appear in only some of the peers.

### Identification of OTUs

To assess the effectiveness of gLinDA in identifying significantly abundant OTUs, we compared the total number of OTUs identified by LinDA, gLinDA, meta-analysis, and individual peers. Figure 1 presents the results for the union approach across four scenarios: two, three, four, and eight peers, with data evenly distributed among them. As expected, individual peers were less effective at identifying OTUs, highlighting the importance of data aggregation in decentralized learning to enhance detection capability. Notably, in the two-peer scenario, gLinDA performed almost as well as centralized LinDA, with only a slight decrease in identified OTUs as the number of peers increased. Most importantly, while gLinDA’s performance declines slightly with more peers, it consistently surpasses each individual peer in all scenarios.

**Figure 1.**
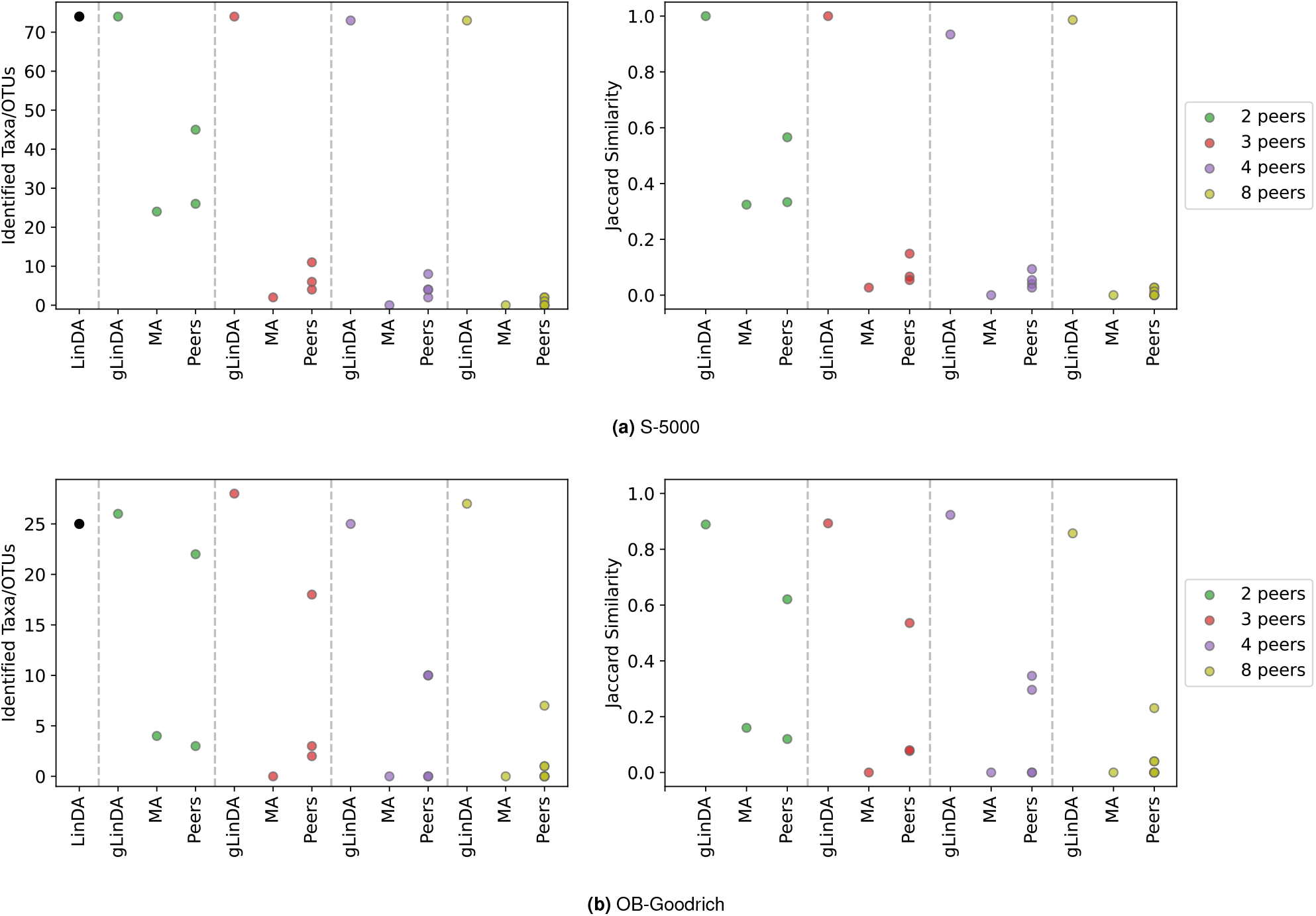
Number of identified taxa/OTUs using LinDA, gLinDA with two, three, four, and eight peers, and meta-analysis, along with the corresponding Jaccard similarity based on LinDA’s predictions for the datasets **(a)** S-5000 and **(b)** OB-Goodrich.

In addition to the total number of identified OTUs, it is crucial to assess whether the identified OTUs match those predicted by standalone LinDA. To evaluate this, the Jaccard similarity was calculated and compared across different peer setups, including meta-analysis. The results, shown in Figure 1 confirm that gLinDA consistently identified the same OTUs as LinDA in almost all scenarios. The figures for the intersection approach, as well as for other datasets, are provided in the supplementary material.

Table 1 summarizes the results for all peer configurations using both union and intersection approaches for the two datasets, S-5000 and OB-Goodrich. The table presents key metrics, including the number of true positives, true negatives, false positives, false negatives, Matthews Correlation Coefficient, F1 score, and Jaccard similarity index in comparison with centralized LinDA. These metrics clearly demonstrate that gLinDA outperforms individual nodes as well as meta-analysis. Additional results for other datasets are available in the supplementary material.

**Table 1.**
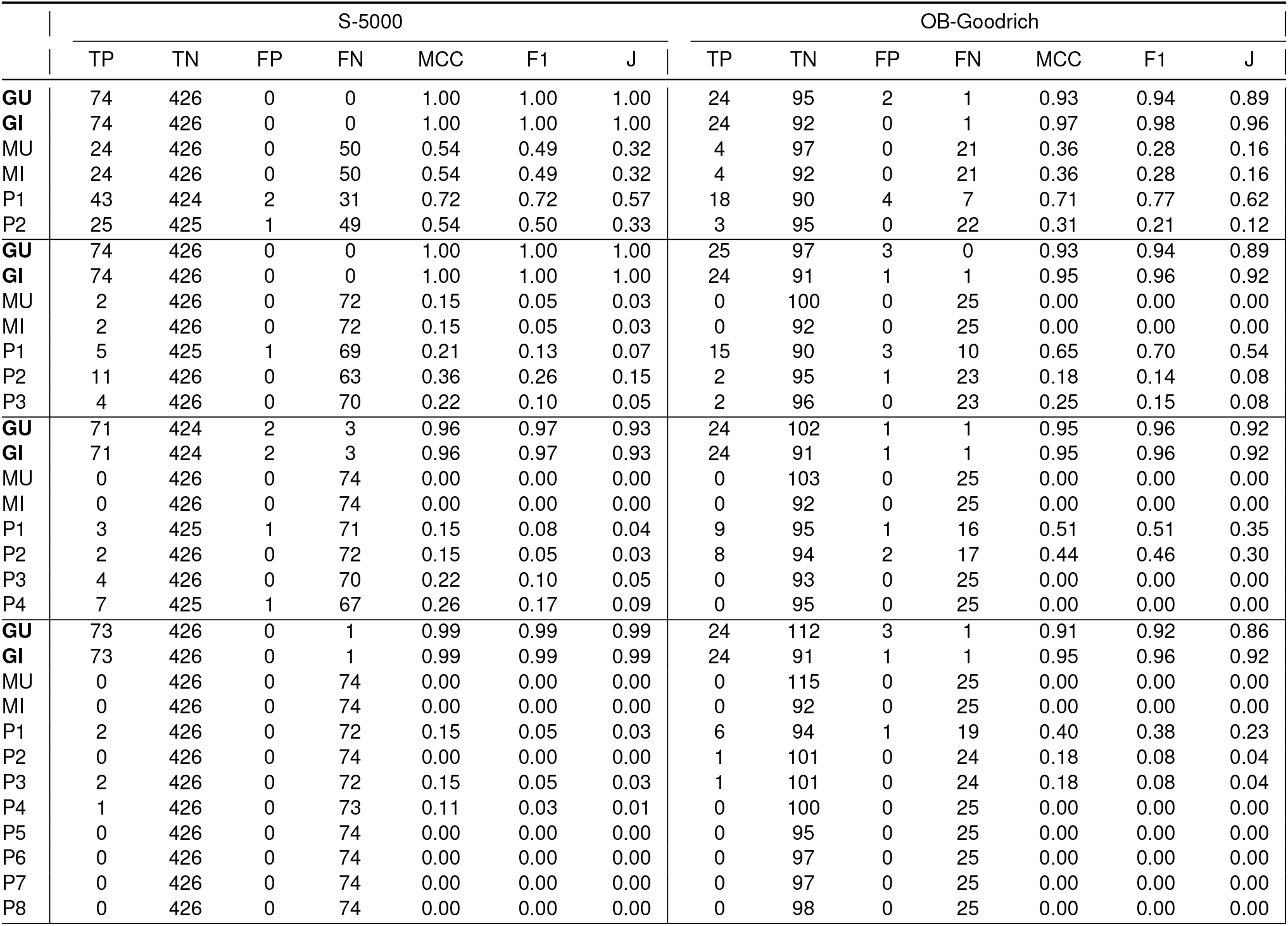
Benchmarks on the dataset S-5000 and OB-Goodrich, depicts the predictive performance including the Jaccard similarity (**J**) of gLinDA with the unified approach (**GU**), gLinDA with the intersectional approach (**GI**), meta-analysis with the unified approach (**MU**), the meta-analysis with the intersectional approach (**MI**), and the single peers (**P1-P8**).

**Table 2.** Names and dimensions of the datasets used for benchmarking as well as the target variables used for DAA; results of non-bold datasets are shown in the supplement.

| Dataset | Reference | Type | Samples | Taxa/OTUs | Target |
| --- | --- | --- | --- | --- | --- |
| **S-5000** | **Yang and Chen (2023)** | **Sim. OTU** | **5000** | **500** | **sim. group** |
| **OB-Goodrich** | **Goodrich et al. (2014)** | **Genus** | **613** | **200** | **obesity** |
| AGP-Diabetes | McDonald et al. (2018) | Genus | 8135 | 537 | diabetes |
| AGP-Ibs | McDonald et al. (2018) | Genus | 1594 | 537 | IBS |
| CDI-Shubert | Schubert et al. (2014) | Genus | 237 | 160 | CDI |
| Arctic-Freshwater | Crump et al. (2012) | OTU | 2466 | 3689 | location |
| GWMC-Hot-Cold | Wu et al. (2019) | OTU | 1021 | 4402 | temperature |
| Smoke | Charlson et al. (2010) | OTU | 145 | 2156 | smoking |

### Exact p-values

In the previous section, we evaluated gLinDA’s ability to identify OTUs within a 95 % confidence interval. This section examines how closely the p-values from gLinDA align with those generated by LinDA. Figure 2 displays the p-values for each OTU, comparing LinDA with gLinDA, meta-analysis, and individual peers under the union approach, where data is equally distributed across three peers.

**Figure 2.**
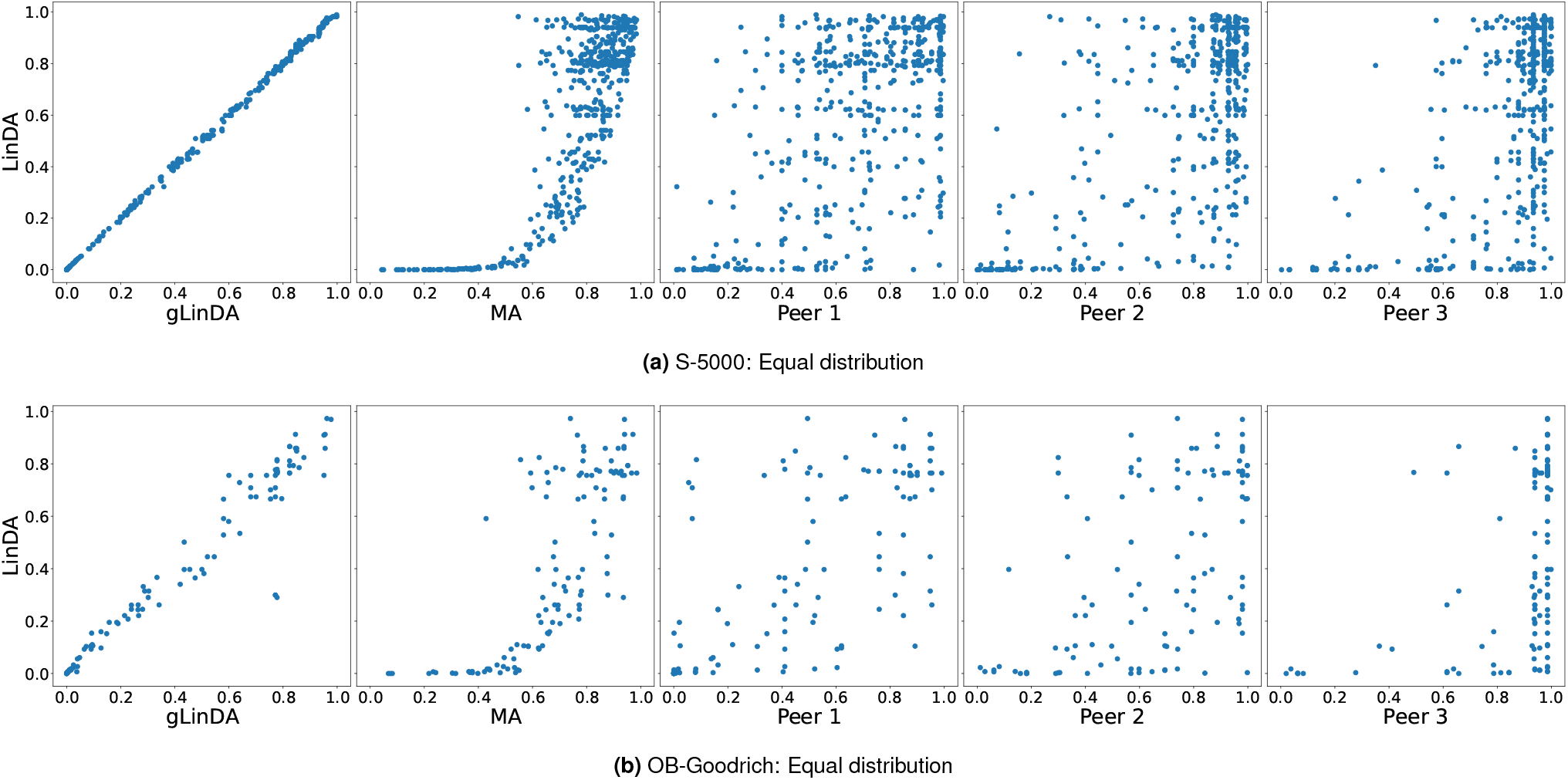
Scatter plots presenting the exact p-values for each OTU, comparing LinDA with gLinDA, meta-analysis (MA), and individual peers under the union approach, where data is equally distributed across three peers for the **(a)** S-5000 and **(b)** OB-Goodrich datasets.

The findings reveal that gLinDA not only identifies the same OTUs as LinDA but also produces p-values that align closely with those of LinDA, outperforming individual peers and meta-analyses. In the scatter plot for the S-5000 dataset shown in Figure 2a, gLinDA forms an almost perfect diagonal line when compared to LinDA. However, for the OB-Goodrich dataset, as presented in Figure 2b, the p-values are more dispersed, likely due to the dataset’s eight-fold smaller sample size. Overall, gLinDA consistently outperforms both meta-analysis and individual peers.

For completeness, all datasets were tested under various scenarios, ranging from two to eight peers, using both union and intersection modes, illustrated in the supplementary information. The results consistently demonstrate that gLinDA’s performance is robust and scalable across different settings.

To further ensure that these results are robust, the p-value benchmarks were repeated with unequally distributed datasets among three peers with ratios of 0.1, 0.3, and 0.6, as shown in Figure 3. The performance trends remained unchanged, indicating that gLinDA’s effectiveness is independent of the dataset distribution across peers. This evaluation represents a more realistic scenario, addressing concerns of fairness in decentralized learning approaches (Tajabadi et al., 2023; Tajabadi and Heider, 2024) and assessing the impact of single large cross-silo peers on the overall performance of the decentralized learning model.

**Figure 3.**
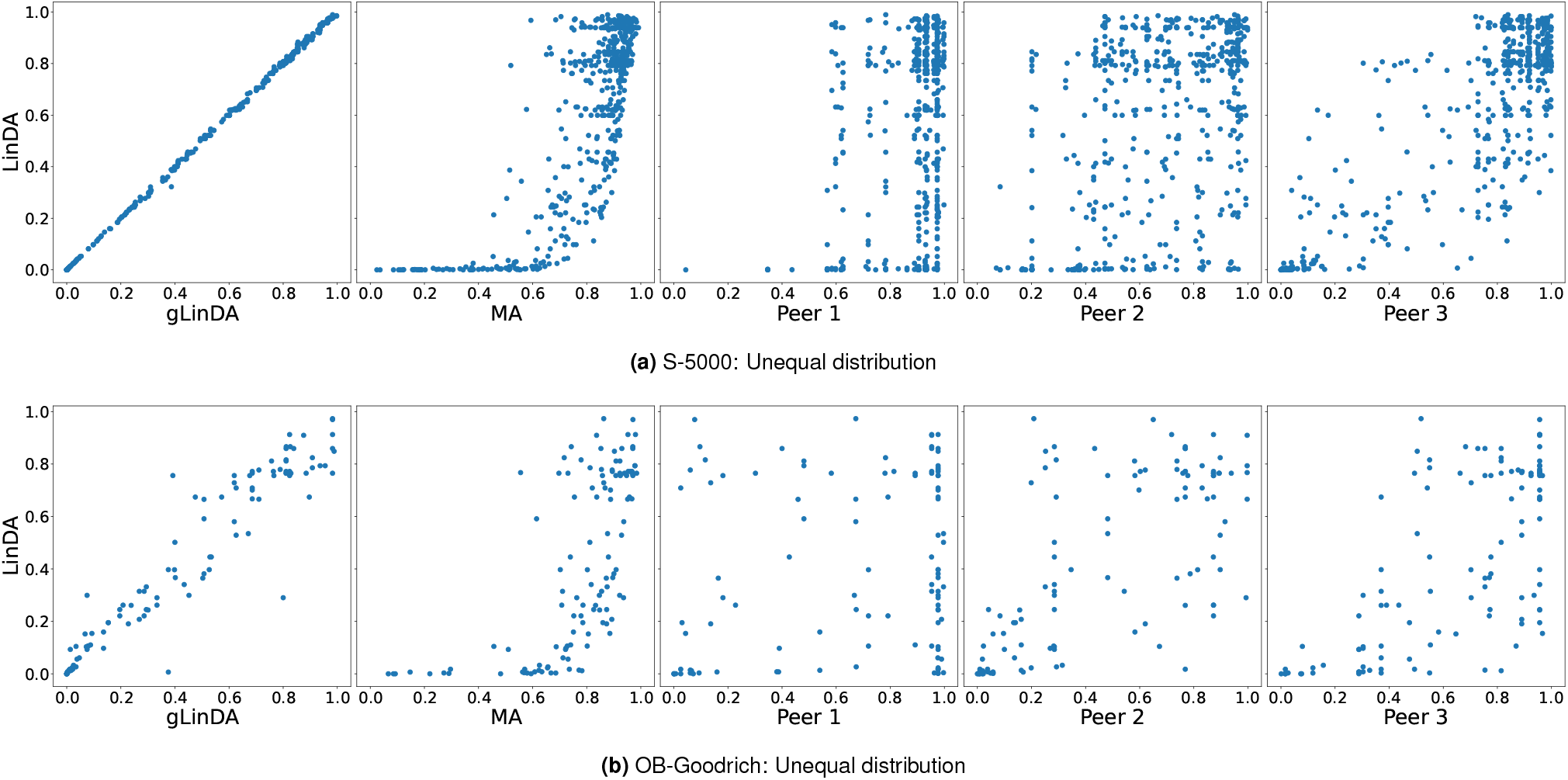
Scatter plots presenting the exact p-values for each OTU, comparing LinDA, gLinDA, meta-analysis (MA), and individual peers under the union approach, where data is **unequally** distributed across three peers for the **(a)** S-5000 and **(b)** OB-Goodrich datasets. The three peers contain 10 %, 30 %, and 60 % of the entire dataset, respectively.

## Discussion

Considering the performance of OTU identification in the decentralized gLinDA approach, the Jaccard similarity ranges from 0.86 to 1.00, while the MCC ranges from 0.93 to 1.00, as detailed in Table 1. MCC is widely regarded as a state-of-the-art metric for classification tasks and ranges from -1 to 1(Chicco and Jurman, 2020), and these results suggest that gLinDA offers robust performance compared to LinDA. Although there is a slight trend toward lower performance with an increasing number of peers, this largely depends on dataset size and how the data is distributed across peers. Importantly, the secondary goal of the gLinDA benchmark, which compares direct p-values between gLinDA and LinDA, demonstrates that it is possible to achieve similar or closely matching p-values, though this is highly dependent on dataset size and split ratios.

It is important to recognize that we evaluated LinDA’s performance on the entire dataset because of its established accuracy as a widely used DAA algorithm. However, this approach does not eliminate the possibility that some OTUs identified as significant may be false positives, or that some non-identified OTUs may, in fact, be true positives. This introduces a degree of variance in our results, as we assume LinDA outcomes to be the ground truth.

Moreover, we worked with artificially equal and unequal split datasets to simulate cross-silo performance in individual peers, thereby excluding potential issues such as batch effects. These effects can be minimized by applying strict standardization and quality control during data collection and measurement (Leek et al., 2010).

Although gLinDA successfully aggregates models from different peers, a minimum peer size is necessary to achieve robust results. If a peer’s dataset is too small, the outcome lacks statistical power and tends to show an increased false discovery rate (Lin and Peddada, 2020; Zhou et al., 2022). In one unequal benchmark, we found that no significant results were detected for a peer with fewer than 15 samples.

## Conclusion

gLinDA implements a swarm learning-based peer-to-peer version of LinDA, a state-of-the-art tool for Differential Abundance Analysis (DAA). Throughout this manuscript, we demonstrated that gLinDA’s decentralized approach can effectively match the predictive performance of the traditional R-based LinDA implementation. Moreover, when examining the performance of individual peers on partitions of a dataset, the results clearly emphasize how aggregating more data significantly improves performance. This suggests that hospitals or institutions conducting DAA would benefit greatly from using gLinDA, especially when working with relatively small datasets. Additionally, we showed that the decentralized approach frequently outperforms individual peers, even when those peers possess a substantial portion of the dataset. This is particularly evident when comparing the distinct OTUs’ p-values between LinDA and individual peers.

## Methods

The original LinDA implementation, written in R, uses linear regression models on log-transformed microbiome data. However, R packages can be more challenging to access, particularly for users with limited computational experience, such as many life scientists. To facilitate broader adoption and serve as the foundation for the P2P network, the entire LinDA codebase was rewritten in Python 3. Each function and dependency was rigorously tested to ensure that gLinDA mirrors the behavior of the original R version. Multiple datasets with various settings were used to confirm that the Python translation of LinDA produces identical results to the original R implementation.

The graphical user interface GUI uses the Qt6 library and was written with PyQt6, ensuring a robustness base with high compatibility for Linux, MacOS, and Windows. Moreover, the interface allows less computer-savvy users to apply LinDA computations alone or within a P2P network. A screenshot of gLinDA is shown in Figure S1. The user interface is optional since all gLinDA computations can be performed via the command-line interface. gLinDA operates cross-interface, so each peer can use whatever interface is preferred.

### gLinDA implementation: Decentralizing LinDA

Since we did not modify any of the core functionality of LinDA, all relevant details regarding the basic linear model algorithm are comprehensively discussed in the original publication by Zhou et al. (2022). Therefore, we will only outline the necessary modifications that pertain to the decentralized learning approach of gLinDA.

Each peer initially builds linear regressions on their respective local datasets, using the covariates of interest and the target variable. Specifically, for each taxon *i*, peers calculate:

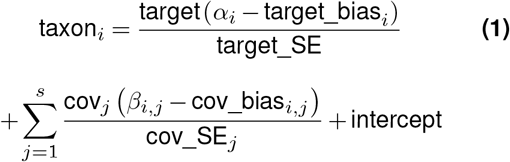

Afterward, each peer transmits its model parameters to all other peers within the network. These parameters consist of the regression coefficients and the standard errors associated with each covariate in the model. Upon receiving the parameters from the other peers, each peer aggregates the information by computing a weighted average of the coefficients and standard errors, as described in Formulas 2 and 3, thereby constructing a global model that integrates data from all peers across the network.

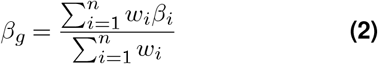

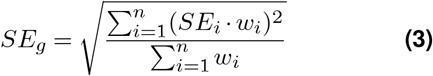

Then, the peers calculate the t-statistic by dividing the global coefficients by their respective global standard errors, which they use to calculate the p-values, as shown in Formula 4.

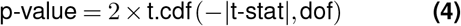

### Network implementation

For peer-to-peer communication in swarm learning, gLinDA utilizes basic socket communication. To ensure secure network transmission, gLinDA employs 256-bit AES (Advanced Encryption Standard) encryption with temporarily generated keys. For the initial handshake, a common AES key, derived from peer addresses and a shared password, is used across all peers. During this handshake, the client sends a nonce (number used once), and both the client and server generate a new AES key based on confirmation number derived from the nonce. This key is valid only for the current gLinDA network session, preventing replay attacks. Additionally, all AES keys undergo 100,000 iterations of SHA-512 hashing to resist brute-force attacks. The entire handshake and encryption procedure is illustrated in Figure S2. To further safeguard communication, data are divided into small encrypted packets, mitigating issues related to network transmission limits, such as the maximum transmission unit (MTU).

### Benchmarks

Since the R LinDA code was entirely re-implemented and verified to produce identical output, any reference to LinDA results actually pertains to gLinDA operating in “single mode,” where the analysis is performed without the network communication aspect on the entire dataset.

We gathered multiple publicly available datasets from the literature to conduct gLinDA benchmarks as listed in Table 2. With the exception of one, all datasets are real-world examples. To mitigate potential data impurities or artifacts, we also included the S-5000 dataset, based on simulated data, as the sole artificial dataset, to demonstrate gLinDA’s algorithmic performance under controlled conditions.

All benchmarks were conducted with one, two, three, four, and eight peers in both intersection and union modes across all datasets, with the exception of the unequal split benchmark. For this benchmark, we opted to perform only one highly imbalanced test with three peers and the ratios of 0.1, 0.3, and 0.6 across all datasets. The complete results are provided in the supplementary information.

We compared the performance of gLinDA and the individual peer models to a conventional meta-analysis approach. In situations where pooling data is either impractical or undesirable, meta-analysis is typically performed by aggregating the end-results from multiple local analyses into a global summary. For DAA, this often involves combining p-values from local models at each site, using methods such as weighted averaging, where the weights are based on the sample size of each site.

To facilitate this comparison between the meta-analysis approach and gLinDA, we first calculated local p-values at each peer during our benchmarking process. These p-values were then shared across all peers, allowing for the calculation of a global p-value via a weighted average. We then compared the global p-values from the meta-analysis, the gLinDA-derived p-values, and the individual peer-level p-values. Their classification as true or false positives, or true or false negatives, was determined by comparing their significance levels to those obtained by LinDA when applied to the full dataset.

For the comparison benchmarks, we calculated the Matthews correlation coefficient (MCC) as shown in Formula 5, the F1-score in Formula 6, and the Jaccard similarity represented in Formula 7, assuming that the outcome of LinDA on the entire dataset represents the ground truth:

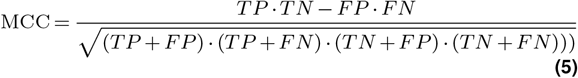

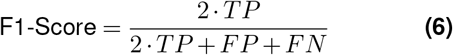

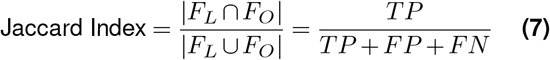

For the Jaccard Index *F*_*L*_ represents the set of OTUs found by LinDA and *F*_*O*_ is the set of OTUs found by any other method that is compared with LinDA.

## Declarations

- **Funding:** This research was funded by the German Federal Ministry of Education and Research (BMBF) under the project PerMed-COPD (grant number 01EK2203F).
- **Conflict of interest**: The authors declare no conflict of interest.
- **Code availability:** The entire MIT-licensed source code of gLinDA is publicly available at https://imigitlab.uni-muenster.de/published/glinda or at Zenodo (DOI: 10.5281/zenodo.13843588).
- **Author contributions**: **Leon Fehse**: Conceptualization (equal), Data Curation (equal), Methodology (equal), Software (equal), Writing – Original Draft Preparation (equal), Writing – Review & Editing (equal). **Mohammad Tajabadi**: Conceptualization (equal), Data Curation (equal), Methodology (equal), Software (equal), Writing – Original Draft Preparation (equal), Writing – Review & Editing (equal). **Roman Martin**: Conceptualization (equal), Data Curation (equal), Methodology (equal), Software (equal), Methodology (equal), Writing – Original Draft Preparation (equal), Writing – Review & Editing (equal). **Hajo Holzmann**: Methodology (supporting). **Dominik Heider**: Conceptualization (equal), Supvervision, Writing – Original Draft Preparation (supporting), Writing – Review & Editing (equal)

