## Supplementary Information for "gLinDA: Global Differential Abundance Analysis of Microbiomes"

### This PDF file includes:

Tables S1 to S13
Figs. S1 to S65

**Fig. S1.** Screenshot of the first complete gLinDA graphical user interface.

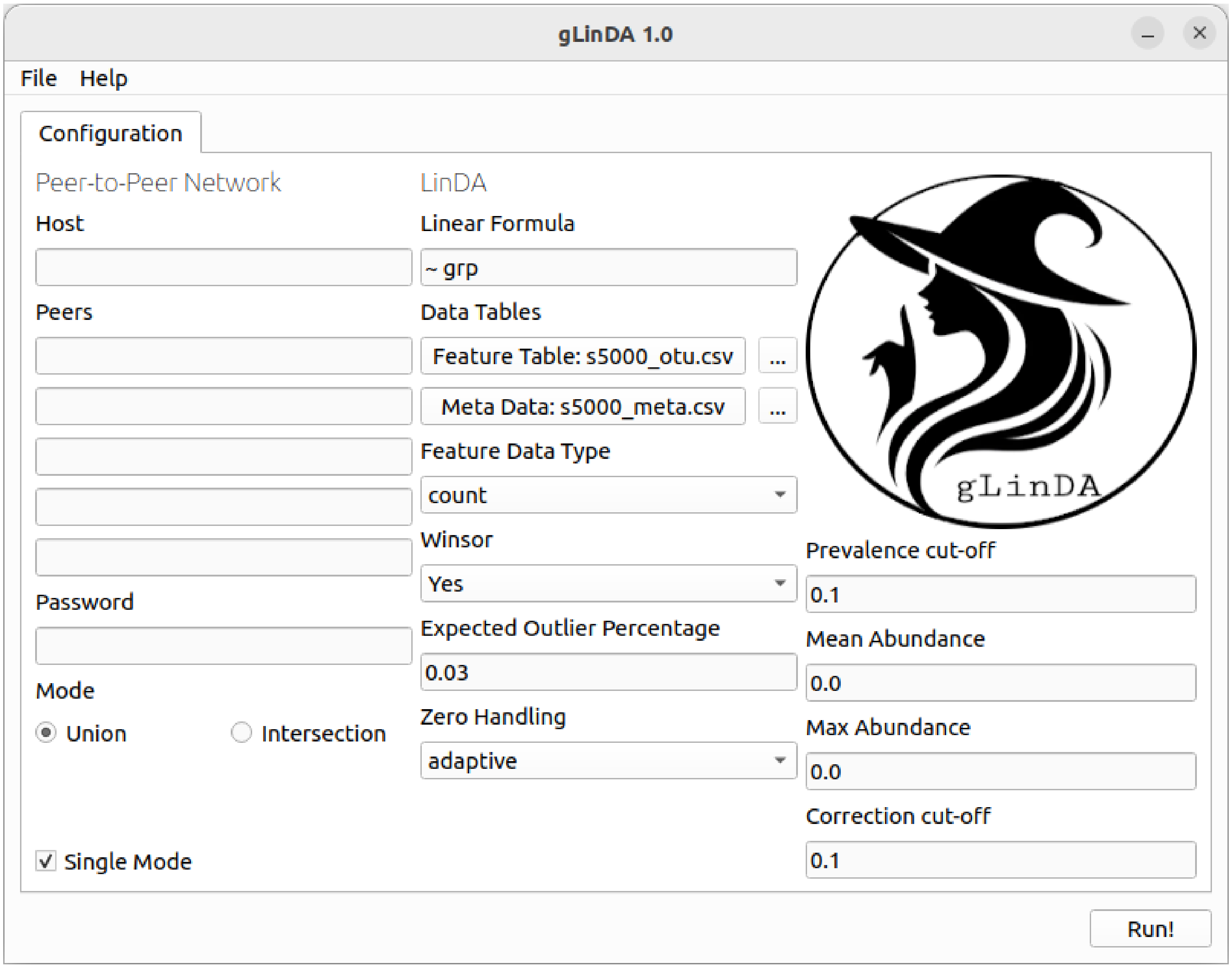


 Leon Fehse, Mohammad Tajabadi, Roman Martin, Hajo Holzmann, Dominik Heider

**Fig. S2.** Illustration of the peer-to-peer swarm learning network communication between a client and server module from two gLinDA peers. **(1)** The server and client modules generate a 256-bit AES key, derived from a shared password and salt, and hashed 100,000 times using SHA-512. This **initial AES key** is used for the first interaction. **(2)** The client module sends a nonce (a random number used once) to the server module of the other peer. **(3)** The server module confirms successful decryption by sending a confirmation number (conce) based on the nonce back to the client. **(4)** Both the client and server modules then generate a new **session AES key** for continued communication, based on the password, nonce, and conce. Any subsequent submission from the client **(5)** will be securely responded to **(6)** using the session AES key, maintaining encryption integrity until a new gLinDA execution is initiated.

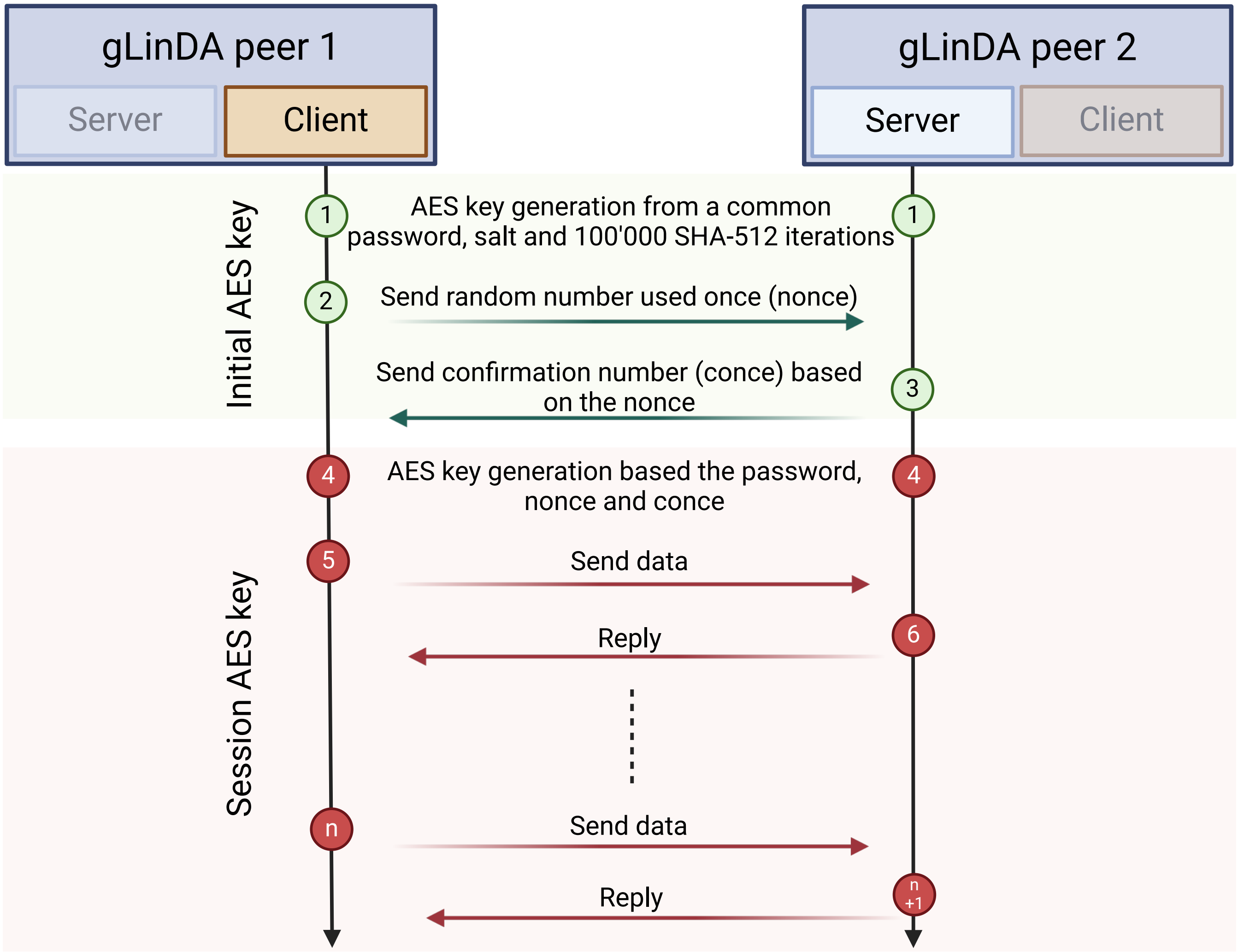

**Table S1. Benchmarks on the AGP-Ibs dataset, depicting the predictive performance of gLinDA with the unified approach, gLinDA with the intersectional approach, the meta-analysis, and the single peers.**

| | TP | TN | FP | FN | MCC | F1 | Jaccard |
|---|---|---|---|---|---|---|---|
| gLinDA (Union) | 6 | 57 | 0 | 6 | 0.67 | 0.67 | 0.50 |
| gLinDA (Intersection) | 7 | 54 | 0 | 5 | 0.73 | 0.74 | 0.58 |
| Meta Analysis (Union) | 1 | 57 | 0 | 11 | 0.26 | 0.15 | 0.08 |
| Meta Analysis (Intersection) | 1 | 54 | 0 | 11 | 0.26 | 0.15 | 0.08 |
| Peer 1 | 3 | 55 | 0 | 9 | 0.46 | 0.40 | 0.25 |
| Peer 2 | 4 | 56 | 0 | 8 | 0.54 | 0.50 | 0.33 |
| gLinDA (Union) | 6 | 57 | 1 | 6 | 0.61 | 0.63 | 0.46 |
| gLinDA (Intersection) | 6 | 54 | 0 | 6 | 0.67 | 0.67 | 0.50 |
| Meta Analysis (Union) | 0 | 58 | 0 | 12 | 0.00 | 0.00 | 0.00 |
| Meta Analysis (Intersection) | 0 | 54 | 0 | 12 | 0.00 | 0.00 | 0.00 |
| Peer 1 | 0 | 56 | 0 | 12 | 0.00 | 0.00 | 0.00 |
| Peer 2 | 0 | 55 | 0 | 12 | 0.00 | 0.00 | 0.00 |
| Peer 3 | 1 | 56 | 0 | 11 | 0.26 | 0.15 | 0.08 |
| gLinDA (Union) | 4 | 53 | 5 | 8 | 0.28 | 0.38 | 0.24 |
| gLinDA (Intersection) | 6 | 49 | 5 | 6 | 0.42 | 0.52 | 0.35 |
| Meta Analysis (Union) | 0 | 58 | 0 | 12 | 0.00 | 0.00 | 0.00 |
| Meta Analysis (Intersection) | 0 | 54 | 0 | 12 | 0.00 | 0.00 | 0.00 |
| Peer 1 | 0 | 56 | 0 | 12 | 0.00 | 0.00 | 0.00 |
| Peer 2 | 2 | 54 | 1 | 10 | 0.28 | 0.27 | 0.15 |
| Peer 3 | 0 | 56 | 0 | 12 | 0.00 | 0.00 | 0.00 |
| Peer 4 | 2 | 56 | 0 | 10 | 0.38 | 0.29 | 0.17 |
| gLinDA (Union) | 12 | 63 | 0 | 0 | 1.00 | 1.00 | 1.00 |
| gLinDA (Intersection) | 12 | 53 | 1 | 0 | 0.95 | 0.96 | 0.92 |
| Meta Analysis (Union) | 0 | 63 | 0 | 12 | 0.00 | 0.00 | 0.00 |
| Meta Analysis (Intersection) | 0 | 54 | 0 | 12 | 0.00 | 0.00 | 0.00 |
| Peer 1 | 0 | 56 | 0 | 12 | 0.00 | 0.00 | 0.00 |
| Peer 2 | 0 | 55 | 0 | 12 | 0.00 | 0.00 | 0.00 |
| Peer 3 | 0 | 55 | 0 | 12 | 0.00 | 0.00 | 0.00 |
| Peer 4 | 0 | 56 | 0 | 12 | 0.00 | 0.00 | 0.00 |
| Peer 5 | 0 | 57 | 0 | 12 | 0.00 | 0.00 | 0.00 |
| Peer 6 | 0 | 59 | 0 | 12 | 0.00 | 0.00 | 0.00 |
| Peer 7 | 1 | 56 | 0 | 11 | 0.26 | 0.15 | 0.08 |
| Peer 8 | 0 | 55 | 0 | 12 | 0.00 | 0.00 | 0.00 |

**Table S2. Benchmarks on the AGP-Diabetes dataset, depicting the predictive performance of gLinDA with the unified approach, gLinDA with the intersectional approach, the meta-analysis, and the single peers.**

| | TP | TN | FP | FN | MCC | F1 | Jaccard |
|---|---|---|---|---|---|---|---|
| gLinDA (Union) | 7 | 59 | 0 | 0 | 1.00 | 1.00 | 1.00 |
| gLinDA (Intersection) | 7 | 59 | 0 | 0 | 1.00 | 1.00 | 1.00 |
| Meta Analysis (Union) | 2 | 59 | 0 | 5 | 0.51 | 0.44 | 0.29 |
| Meta Analysis (Intersection) | 2 | 59 | 0 | 5 | 0.51 | 0.44 | 0.29 |
| Peer 1 | 3 | 58 | 1 | 4 | 0.53 | 0.55 | 0.38 |
| Peer 2 | 2 | 58 | 1 | 5 | 0.40 | 0.40 | 0.25 |
| gLinDA (Union) | 7 | 58 | 1 | 0 | 0.93 | 0.93 | 0.88 |
| gLinDA (Intersection) | 7 | 58 | 1 | 0 | 0.93 | 0.93 | 0.88 |
| Meta Analysis (Union) | 0 | 59 | 0 | 7 | 0.00 | 0.00 | 0.00 |
| Meta Analysis (Intersection) | 0 | 59 | 0 | 7 | 0.00 | 0.00 | 0.00 |
| Peer 1 | 1 | 58 | 1 | 6 | 0.23 | 0.22 | 0.12 |
| Peer 2 | 0 | 59 | 0 | 7 | 0.00 | 0.00 | 0.00 |
| Peer 3 | 0 | 59 | 0 | 7 | 0.00 | 0.00 | 0.00 |
| gLinDA (Union) | 7 | 61 | 0 | 0 | 1.00 | 1.00 | 1.00 |
| gLinDA (Intersection) | 7 | 58 | 1 | 0 | 0.93 | 0.93 | 0.88 |
| Meta Analysis (Union) | 0 | 61 | 0 | 7 | 0.00 | 0.00 | 0.00 |
| Meta Analysis (Intersection) | 0 | 59 | 0 | 7 | 0.00 | 0.00 | 0.00 |
| Peer 1 | 0 | 59 | 0 | 7 | 0.00 | 0.00 | 0.00 |
| Peer 2 | 0 | 59 | 0 | 7 | 0.00 | 0.00 | 0.00 |
| Peer 3 | 0 | 60 | 0 | 7 | 0.00 | 0.00 | 0.00 |
| Peer 4 | 1 | 60 | 0 | 6 | 0.36 | 0.25 | 0.14 |
| gLinDA (Union) | 7 | 62 | 0 | 0 | 1.00 | 1.00 | 1.00 |
| gLinDA (Intersection) | 7 | 57 | 2 | 0 | 0.87 | 0.88 | 0.78 |
| Meta Analysis (Union) | 0 | 62 | 0 | 7 | 0.00 | 0.00 | 0.00 |
| Meta Analysis (Intersection) | 0 | 59 | 0 | 7 | 0.00 | 0.00 | 0.00 |
| Peer 1 | 1 | 58 | 1 | 6 | 0.23 | 0.22 | 0.12 |
| Peer 2 | 0 | 59 | 0 | 7 | 0.00 | 0.00 | 0.00 |
| Peer 3 | 0 | 60 | 0 | 7 | 0.00 | 0.00 | 0.00 |
| Peer 4 | 0 | 59 | 0 | 7 | 0.00 | 0.00 | 0.00 |
| Peer 5 | 0 | 58 | 2 | 7 | -0.06 | 0.00 | 0.00 |
| Peer 6 | 0 | 61 | 0 | 7 | 0.00 | 0.00 | 0.00 |
| Peer 7 | 0 | 60 | 0 | 7 | 0.00 | 0.00 | 0.00 |
| Peer 8 | 2 | 58 | 1 | 5 | 0.40 | 0.40 | 0.25 |

**Table S3. Benchmarks on the Arctic-Freshwater dataset, depicting the predictive performance of gLinDA with the unified approach, gLinDA with the intersectional approach, the meta-analysis, and the single peers.**

| | TP | TN | FP | FN | MCC | F1 | Jaccard |
|---|---|---|---|---|---|---|---|
| gLinDA (Union) | 232 | 44 | 5 | 1 | 0.92 | 0.99 | 0.97 |
| gLinDA (Intersection) | 224 | 40 | 2 | 9 | 0.86 | 0.98 | 0.95 |
| Meta Analysis (Union) | 209 | 49 | 0 | 24 | 0.78 | 0.95 | 0.90 |
| Meta Analysis (Intersection) | 203 | 42 | 0 | 30 | 0.71 | 0.93 | 0.87 |
| Peer 1 | 213 | 40 | 4 | 20 | 0.73 | 0.95 | 0.90 |
| Peer 2 | 215 | 44 | 3 | 18 | 0.77 | 0.95 | 0.91 |
| gLinDA (Union) | 229 | 46 | 16 | 4 | 0.79 | 0.96 | 0.92 |
| gLinDA (Intersection) | 216 | 40 | 2 | 17 | 0.78 | 0.96 | 0.92 |
| Meta Analysis (Union) | 182 | 61 | 1 | 51 | 0.64 | 0.88 | 0.78 |
| Meta Analysis (Intersection) | 176 | 42 | 0 | 57 | 0.57 | 0.86 | 0.76 |
| Peer 1 | 197 | 42 | 2 | 36 | 0.65 | 0.91 | 0.84 |
| Peer 2 | 196 | 55 | 5 | 37 | 0.66 | 0.90 | 0.82 |
| Peer 3 | 198 | 39 | 6 | 35 | 0.60 | 0.91 | 0.83 |
| gLinDA (Union) | 230 | 56 | 17 | 3 | 0.81 | 0.96 | 0.92 |
| gLinDA (Intersection) | 211 | 31 | 11 | 22 | 0.59 | 0.93 | 0.86 |
| Meta Analysis (Union) | 156 | 66 | 7 | 77 | 0.49 | 0.79 | 0.65 |
| Meta Analysis (Intersection) | 150 | 42 | 0 | 83 | 0.47 | 0.78 | 0.64 |
| Peer 1 | 168 | 18 | 26 | 65 | 0.10 | 0.79 | 0.65 |
| Peer 2 | 191 | 47 | 9 | 42 | 0.56 | 0.88 | 0.79 |
| Peer 3 | 184 | 55 | 2 | 49 | 0.63 | 0.88 | 0.78 |
| Peer 4 | 190 | 40 | 8 | 43 | 0.54 | 0.88 | 0.79 |
| gLinDA (Union) | 229 | 61 | 33 | 4 | 0.72 | 0.93 | 0.86 |
| gLinDA (Intersection) | 172 | 18 | 24 | 61 | 0.13 | 0.80 | 0.67 |
| Meta Analysis (Union) | 123 | 79 | 15 | 110 | 0.34 | 0.66 | 0.50 |
| Meta Analysis (Intersection) | 104 | 42 | 0 | 129 | 0.33 | 0.62 | 0.45 |
| Peer 1 | 142 | 40 | 13 | 91 | 0.28 | 0.73 | 0.58 |
| Peer 2 | 163 | 42 | 3 | 70 | 0.48 | 0.82 | 0.69 |
| Peer 3 | 158 | 57 | 6 | 75 | 0.48 | 0.80 | 0.66 |
| Peer 4 | 150 | 60 | 2 | 83 | 0.50 | 0.78 | 0.64 |
| Peer 5 | 145 | 66 | 6 | 88 | 0.46 | 0.76 | 0.61 |
| Peer 6 | 142 | 51 | 0 | 91 | 0.47 | 0.76 | 0.61 |
| Peer 7 | 173 | 43 | 4 | 60 | 0.51 | 0.84 | 0.73 |
| Peer 8 | 160 | 50 | 7 | 73 | 0.45 | 0.80 | 0.67 |

**Table S4. Benchmarks on the CDI-Schubert dataset, depicting the predictive performance of gLinDA with the unified approach, gLinDA with the intersectional approach, the meta-analysis, and the single peers.**

| | TP | TN | FP | FN | MCC | F1 | Jaccard |
|---|---|---|---|---|---|---|---|
| gLinDA (Union) | 52 | 20 | 3 | 6 | 0.74 | 0.92 | 0.85 |
| gLinDA (Intersection) | 54 | 18 | 0 | 4 | 0.87 | 0.96 | 0.93 |
| Meta Analysis (Union) | 40 | 20 | 3 | 18 | 0.51 | 0.79 | 0.66 |
| Meta Analysis (Intersection) | 40 | 18 | 0 | 18 | 0.59 | 0.82 | 0.69 |
| Peer 1 | 42 | 18 | 2 | 16 | 0.55 | 0.82 | 0.70 |
| Peer 2 | 52 | 18 | 3 | 6 | 0.72 | 0.92 | 0.85 |
| gLinDA (Union) | 50 | 24 | 10 | 8 | 0.58 | 0.85 | 0.74 |
| gLinDA (Intersection) | 54 | 17 | 1 | 4 | 0.83 | 0.96 | 0.92 |
| Meta Analysis (Union) | 36 | 31 | 3 | 22 | 0.52 | 0.74 | 0.59 |
| Meta Analysis (Intersection) | 33 | 18 | 0 | 25 | 0.49 | 0.73 | 0.57 |
| Peer 1 | 38 | 23 | 0 | 20 | 0.59 | 0.79 | 0.66 |
| Peer 2 | 45 | 22 | 2 | 13 | 0.64 | 0.86 | 0.75 |
| Peer 3 | 39 | 22 | 2 | 19 | 0.54 | 0.79 | 0.65 |
| gLinDA (Union) | 49 | 26 | 15 | 9 | 0.49 | 0.80 | 0.67 |
| gLinDA (Intersection) | 54 | 17 | 1 | 4 | 0.83 | 0.96 | 0.92 |
| Meta Analysis (Union) | 33 | 31 | 10 | 25 | 0.32 | 0.65 | 0.49 |
| Meta Analysis (Intersection) | 31 | 18 | 0 | 27 | 0.46 | 0.70 | 0.53 |
| Peer 1 | 34 | 22 | 0 | 24 | 0.53 | 0.74 | 0.59 |
| Peer 2 | 31 | 22 | 2 | 27 | 0.42 | 0.68 | 0.52 |
| Peer 3 | 44 | 21 | 6 | 14 | 0.51 | 0.81 | 0.69 |
| Peer 4 | 39 | 21 | 7 | 19 | 0.40 | 0.75 | 0.60 |
| gLinDA (Union) | 49 | 51 | 17 | 9 | 0.59 | 0.79 | 0.65 |
| gLinDA (Intersection) | 39 | 11 | 7 | 19 | 0.25 | 0.75 | 0.60 |
| Meta Analysis (Union) | 11 | 66 | 2 | 47 | 0.26 | 0.31 | 0.18 |
| Meta Analysis (Intersection) | 11 | 18 | 0 | 47 | 0.23 | 0.32 | 0.19 |
| Peer 1 | 24 | 26 | 1 | 34 | 0.38 | 0.58 | 0.41 |
| Peer 2 | 13 | 36 | 0 | 45 | 0.32 | 0.37 | 0.22 |
| Peer 3 | 25 | 35 | 1 | 33 | 0.44 | 0.60 | 0.42 |
| Peer 4 | 11 | 30 | 0 | 47 | 0.27 | 0.32 | 0.19 |
| Peer 5 | 30 | 26 | 4 | 28 | 0.37 | 0.65 | 0.48 |
| Peer 6 | 28 | 36 | 1 | 30 | 0.48 | 0.64 | 0.47 |
| Peer 7 | 29 | 24 | 4 | 29 | 0.34 | 0.64 | 0.47 |
| Peer 8 | 15 | 29 | 2 | 43 | 0.24 | 0.40 | 0.25 |

Table S5. **Benchmarks on the GWMC-Hot-Cold dataset, depicting the predictive performance of gLinDA with the unified approach, gLinDA with the intersectional approach, the meta-analysis, and the single peers.**

| | TP | TN | FP | FN | MCC | F1 | Jaccard |
|---|---|---|---|---|---|---|---|
| gLinDA (Union) | 888 | 626 | 29 | 18 | 0.94 | 0.97 | 0.95 |
| gLinDA (Intersection) | 877 | 595 | 21 | 29 | 0.93 | 0.97 | 0.95 |
| Meta Analysis (Union) | 557 | 641 | 14 | 349 | 0.61 | 0.75 | 0.61 |
| Meta Analysis (Intersection) | 545 | 615 | 1 | 361 | 0.61 | 0.75 | 0.60 |
| Peer 1 | 618 | 569 | 71 | 288 | 0.57 | 0.77 | 0.63 |
| Peer 2 | 693 | 577 | 54 | 213 | 0.67 | 0.84 | 0.72 |
| gLinDA (Union) | 883 | 649 | 47 | 23 | 0.91 | 0.96 | 0.93 |
| gLinDA (Intersection) | 844 | 581 | 35 | 62 | 0.87 | 0.95 | 0.90 |
| Meta Analysis (Union) | 367 | 680 | 16 | 539 | 0.44 | 0.57 | 0.40 |
| Meta Analysis (Intersection) | 353 | 616 | 0 | 553 | 0.45 | 0.56 | 0.39 |
| Peer 1 | 443 | 622 | 31 | 463 | 0.47 | 0.64 | 0.47 |
| Peer 2 | 579 | 603 | 40 | 327 | 0.58 | 0.76 | 0.61 |
| Peer 3 | 543 | 625 | 25 | 363 | 0.57 | 0.74 | 0.58 |
| gLinDA (Union) | 884 | 665 | 51 | 22 | 0.91 | 0.96 | 0.92 |
| gLinDA (Intersection) | 843 | 578 | 38 | 63 | 0.86 | 0.94 | 0.89 |
| Meta Analysis (Union) | 235 | 702 | 14 | 671 | 0.33 | 0.41 | 0.26 |
| Meta Analysis (Intersection) | 229 | 616 | 0 | 677 | 0.35 | 0.40 | 0.25 |
| Peer 1 | 392 | 621 | 37 | 514 | 0.42 | 0.59 | 0.42 |
| Peer 2 | 427 | 626 | 16 | 479 | 0.49 | 0.63 | 0.46 |
| Peer 3 | 529 | 596 | 47 | 377 | 0.52 | 0.71 | 0.56 |
| Peer 4 | 351 | 641 | 15 | 555 | 0.42 | 0.55 | 0.38 |
| gLinDA (Union) | 876 | 769 | 75 | 30 | 0.88 | 0.94 | 0.89 |
| gLinDA (Intersection) | 781 | 575 | 41 | 125 | 0.78 | 0.90 | 0.82 |
| Meta Analysis (Union) | 39 | 838 | 6 | 867 | 0.11 | 0.08 | 0.04 |
| Meta Analysis (Intersection) | 35 | 616 | 0 | 871 | 0.13 | 0.07 | 0.04 |
| Peer 1 | 110 | 680 | 5 | 796 | 0.22 | 0.22 | 0.12 |
| Peer 2 | 198 | 668 | 8 | 708 | 0.30 | 0.36 | 0.22 |
| Peer 3 | 72 | 674 | 3 | 834 | 0.17 | 0.15 | 0.08 |
| Peer 4 | 276 | 649 | 8 | 630 | 0.37 | 0.46 | 0.30 |
| Peer 5 | 258 | 658 | 20 | 648 | 0.33 | 0.44 | 0.28 |
| Peer 6 | 287 | 661 | 8 | 619 | 0.39 | 0.48 | 0.31 |
| Peer 7 | 88 | 686 | 2 | 818 | 0.20 | 0.18 | 0.10 |
| Peer 8 | 157 | 660 | 8 | 749 | 0.26 | 0.29 | 0.17 |

**Table S6. Benchmarks on the Smoke dataset, depicting the predictive performance of gLinDA with the unified approach, gLinDA with the intersectional approach, the meta-analysis, and the single peers.**

| | TP | TN | FP | FN | MCC | F1 | Jaccard |
|---|---|---|---|---|---|---|---|
| gLinDA (Union) | 17 | 199 | 1 | 5 | 0.84 | 0.85 | 0.74 |
| gLinDA (Intersection) | 18 | 178 | 4 | 4 | 0.80 | 0.82 | 0.69 |
| Meta Analysis (Union) | 3 | 200 | 0 | 19 | 0.35 | 0.24 | 0.14 |
| Meta Analysis (Intersection) | 2 | 182 | 0 | 20 | 0.29 | 0.17 | 0.09 |
| Peer 1 | 9 | 192 | 2 | 13 | 0.55 | 0.55 | 0.38 |
| Peer 2 | 7 | 187 | 1 | 15 | 0.50 | 0.47 | 0.30 |
| gLinDA (Union) | 19 | 247 | 2 | 3 | 0.87 | 0.88 | 0.79 |
| gLinDA (Intersection) | 17 | 179 | 3 | 5 | 0.79 | 0.81 | 0.68 |
| Meta Analysis (Union) | 0 | 249 | 0 | 22 | 0.00 | 0.00 | 0.00 |
| Meta Analysis (Intersection) | 0 | 182 | 0 | 22 | 0.00 | 0.00 | 0.00 |
| Peer 1 | 9 | 197 | 3 | 13 | 0.52 | 0.53 | 0.36 |
| Peer 2 | 1 | 208 | 0 | 21 | 0.20 | 0.09 | 0.05 |
| Peer 3 | 1 | 216 | 0 | 21 | 0.20 | 0.09 | 0.05 |
| gLinDA (Union) | 10 | 259 | 1 | 12 | 0.62 | 0.61 | 0.43 |
| gLinDA (Intersection) | 10 | 180 | 2 | 12 | 0.58 | 0.59 | 0.42 |
| Meta Analysis (Union) | 0 | 260 | 0 | 22 | 0.00 | 0.00 | 0.00 |
| Meta Analysis (Intersection) | 0 | 182 | 0 | 22 | 0.00 | 0.00 | 0.00 |
| Peer 1 | 4 | 197 | 0 | 18 | 0.41 | 0.31 | 0.18 |
| Peer 2 | 0 | 216 | 0 | 22 | 0.00 | 0.00 | 0.00 |
| Peer 3 | 0 | 205 | 1 | 22 | -0.02 | 0.00 | 0.00 |
| Peer 4 | 0 | 212 | 0 | 22 | 0.00 | 0.00 | 0.00 |
| gLinDA (Union) | 13 | 430 | 1 | 9 | 0.73 | 0.72 | 0.57 |
| gLinDA (Intersection) | 17 | 181 | 1 | 5 | 0.84 | 0.85 | 0.74 |
| Meta Analysis (Union) | 0 | 431 | 0 | 22 | 0.00 | 0.00 | 0.00 |
| Meta Analysis (Intersection) | 0 | 182 | 0 | 22 | 0.00 | 0.00 | 0.00 |
| Peer 1 | 0 | 226 | 0 | 22 | 0.00 | 0.00 | 0.00 |
| Peer 2 | 0 | 217 | 0 | 22 | 0.00 | 0.00 | 0.00 |
| Peer 3 | 1 | 241 | 0 | 21 | 0.20 | 0.09 | 0.05 |
| Peer 4 | 0 | 248 | 0 | 22 | 0.00 | 0.00 | 0.00 |
| Peer 5 | 0 | 248 | 0 | 22 | 0.00 | 0.00 | 0.00 |
| Peer 6 | 0 | 241 | 0 | 22 | 0.00 | 0.00 | 0.00 |
| Peer 7 | 0 | 232 | 0 | 22 | 0.00 | 0.00 | 0.00 |
| Peer 8 | 0 | 273 | 0 | 22 | 0.00 | 0.00 | 0.00 |

**Table S7. Benchmarks on the AGP-Ibs dataset, depicting the predictive performance of gLinDA with the unified approach, gLinDA with the intersectional approach, the meta-analysis, and the single peers with unequal data distribution with ratios 0.1, 0.3, 0.6.**

| | TP | TN | FP | FN | MCC | F1 | Jaccard |
|---|---|---|---|---|---|---|---|
| gLinDA (Union) | 4 | 55 | 4 | 8 | 0.31 | 0.40 | 0.25 |
| gLinDA (Intersection) | 4 | 50 | 4 | 8 | 0.31 | 0.40 | 0.25 |
| Meta Analysis (Union) | 0 | 58 | 0 | 12 | 0.00 | 0.00 | 0.00 |
| Meta Analysis (Intersection) | 0 | 54 | 0 | 12 | 0.00 | 0.00 | 0.00 |
| Peer 1 | 0 | 58 | 0 | 12 | 0.00 | 0.00 | 0.00 |
| Peer 2 | 1 | 55 | 0 | 11 | 0.26 | 0.15 | 0.08 |
| Peer 3 | 8 | 53 | 2 | 4 | 0.68 | 0.73 | 0.57 |

**Table S8. Benchmarks on the AGP-Diabetes dataset, depicting the predictive performance of gLinDA with the unified approach, gLinDA with the intersectional approach, the meta-analysis, and the single peers with unequal data distribution with ratios 0.1, 0.3, 0.6.**

| | TP | TN | FP | FN | MCC | F1 | Jaccard |
|---|---|---|---|---|---|---|---|
| gLinDA (Union) | 7 | 59 | 1 | 0 | 0.93 | 0.93 | 0.88 |
| gLinDA (Intersection) | 7 | 58 | 1 | 0 | 0.93 | 0.93 | 0.88 |
| Meta Analysis (Union) | 0 | 59 | 0 | 7 | 0.00 | 0.00 | 0.00 |
| Meta Analysis (Intersection) | 0 | 59 | 0 | 7 | 0.00 | 0.00 | 0.00 |
| Peer 1 | 1 | 55 | 4 | 6 | 0.09 | 0.17 | 0.09 |
| Peer 2 | 0 | 60 | 0 | 7 | 0.00 | 0.00 | 0.00 |
| Peer 3 | 4 | 56 | 3 | 3 | 0.52 | 0.57 | 0.40 |

**Table S9. Benchmarks on the Arctic-Freshwater dataset, depicting the predictive performance of gLinDA with the unified approach, gLinDA with the intersectional approach, the meta-analysis, and the single peers with unequal data distribution with ratios 0.1, 0.3, 0.6.**

| | TP | TN | FP | FN | MCC | F1 | Jaccard |
|---|---|---|---|---|---|---|---|
| gLinDA (Union) | 223 | 49 | 12 | 10 | 0.77 | 0.95 | 0.91 |
| gLinDA (Intersection) | 205 | 33 | 9 | 28 | 0.58 | 0.92 | 0.85 |
| Meta Analysis (Union) | 185 | 61 | 1 | 48 | 0.66 | 0.88 | 0.79 |
| Meta Analysis (Intersection) | 166 | 42 | 0 | 67 | 0.52 | 0.83 | 0.71 |
| Peer 1 | 135 | 47 | 4 | 98 | 0.38 | 0.73 | 0.57 |
| Peer 2 | 202 | 44 | 2 | 31 | 0.69 | 0.92 | 0.86 |
| Peer 3 | 219 | 46 | 3 | 14 | 0.81 | 0.96 | 0.93 |

**Table S10. Benchmarks on the CDI-Shubert dataset, depicting the predictive performance of gLinDA with the unified approach, gLinDA with the intersectional approach, the meta-analysis, and the single peers with unequal data distribution with ratios 0.1, 0.3, 0.6.**

| | TP | TN | FP | FN | MCC | F1 | Jaccard |
|---|---|---|---|---|---|---|---|
| gLinDA (Union) | 54 | 28 | 4 | 4 | 0.81 | 0.93 | 0.87 |
| gLinDA (Intersection) | 46 | 14 | 4 | 12 | 0.51 | 0.85 | 0.74 |
| Meta Analysis (Union) | 41 | 31 | 3 | 17 | 0.60 | 0.80 | 0.67 |
| Meta Analysis (Intersection) | 27 | 18 | 0 | 31 | 0.41 | 0.64 | 0.47 |
| Peer 1 | 17 | 26 | 0 | 41 | 0.34 | 0.45 | 0.29 |
| Peer 2 | 37 | 19 | 3 | 21 | 0.45 | 0.76 | 0.61 |
| Peer 3 | 51 | 18 | 2 | 7 | 0.73 | 0.92 | 0.85 |

**Table S11. Benchmarks on the GWMC-Hot-Cold dataset, depicting the predictive performance of gLinDA with the unified approach, gLinDA with the intersectional approach, the meta-analysis, and the single peers with unequal data distribution with ratios 0.1, 0.3, 0.6.**

| | TP | TN | FP | FN | MCC | F1 | Jaccard |
|---|---|---|---|---|---|---|---|
| gLinDA (Union) | 885 | 675 | 25 | 21 | 0.94 | 0.97 | 0.95 |
| gLinDA (Intersection) | 858 | 592 | 24 | 48 | 0.90 | 0.96 | 0.92 |
| Meta Analysis (Union) | 411 | 679 | 17 | 495 | 0.48 | 0.62 | 0.45 |
| Meta Analysis (Intersection) | 222 | 615 | 1 | 684 | 0.34 | 0.39 | 0.24 |
| Peer 1 | 37 | 662 | 0 | 869 | 0.13 | 0.08 | 0.04 |
| Peer 2 | 417 | 627 | 26 | 489 | 0.46 | 0.62 | 0.45 |
| Peer 3 | 766 | 581 | 50 | 140 | 0.76 | 0.89 | 0.80 |

**Table S12. Benchmarks on the OB-Goodrich dataset, depicting the predictive performance of gLinDA with the unified approach, gLinDA with the intersectional approach, the meta-analysis, and the single peers with unequal data distribution with ratios 0.1, 0.3, 0.6.**

| | TP | TN | FP | FN | MCC | F1 | Jaccard |
|---|---|---|---|---|---|---|---|
| gLinDA (Union) | 24 | 94 | 5 | 1 | 0.86 | 0.89 | 0.80 |
| gLinDA (Intersection) | 23 | 90 | 2 | 2 | 0.90 | 0.92 | 0.85 |
| Meta Analysis (Union) | 0 | 100 | 0 | 25 | 0.00 | 0.00 | 0.00 |
| Meta Analysis (Intersection) | 3 | 92 | 0 | 22 | 0.31 | 0.21 | 0.12 |
| Peer 1 | 6 | 92 | 3 | 19 | 0.32 | 0.35 | 0.21 |
| Peer 2 | 16 | 92 | 3 | 9 | 0.68 | 0.73 | 0.57 |
| Peer 3 | 6 | 93 | 1 | 19 | 0.40 | 0.38 | 0.23 |

**Table S13. Benchmarks on the S-5000 dataset, depicting the predictive performance of gLinDA with the unified approach, gLinDA with the intersectional approach, the meta-analysis, and the single peers with unequal data distribution with ratios 0.1, 0.3, 0.6.**

| | TP | TN | FP | FN | MCC | F1 | Jaccard |
|---|---|---|---|---|---|---|---|
| gLinDA (Union) | 74 | 425 | 1 | 0 | 0.99 | 0.99 | 0.99 |
| gLinDA (Intersection) | 74 | 425 | 1 | 0 | 0.99 | 0.99 | 0.99 |
| Meta Analysis (Union) | 2 | 426 | 0 | 72 | 0.15 | 0.05 | 0.03 |
| Meta Analysis (Intersection) | 0 | 426 | 0 | 74 | 0.00 | 0.00 | 0.00 |
| Peer 1 | 1 | 426 | 0 | 73 | 0.11 | 0.03 | 0.01 |
| Peer 2 | 0 | 426 | 0 | 74 | 0.00 | 0.00 | 0.00 |
| Peer 3 | 49 | 425 | 1 | 25 | 0.78 | 0.79 | 0.65 |

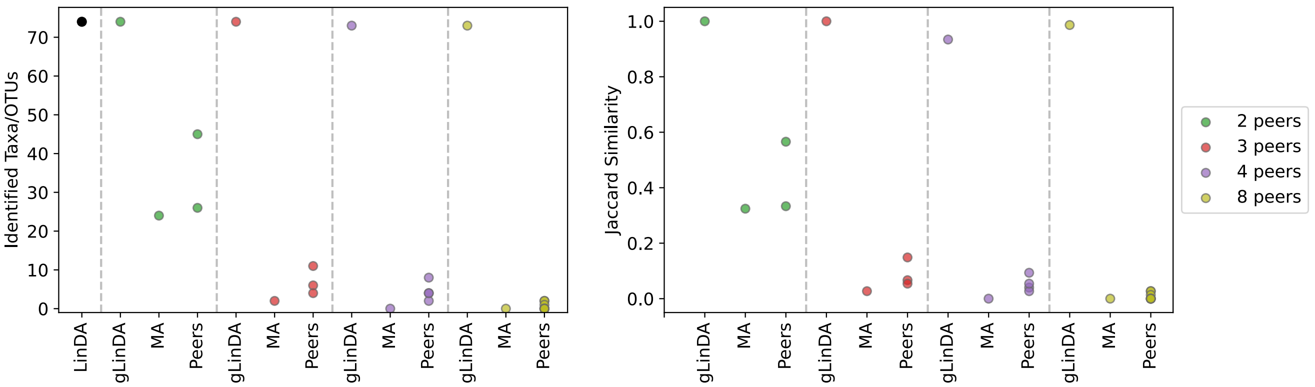


**(a)** Union approach

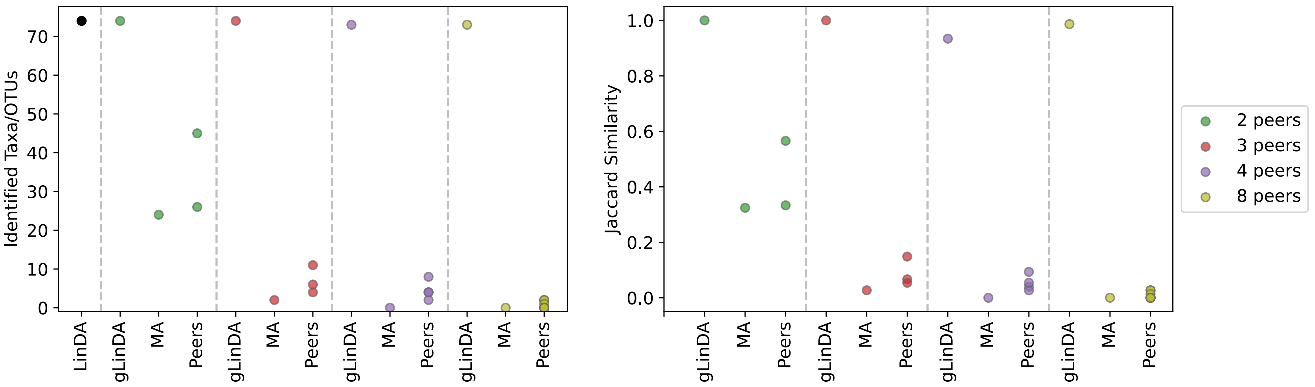


**(b)** Intersection approach

**Fig. S3.** Number of identified taxa/OTUs using LinDA, gLinDA with two, three, four, and eight peers, and meta-analysis, along with the corresponding Jaccard similarity based on LinDA's predictions for the S-5000 dataset.

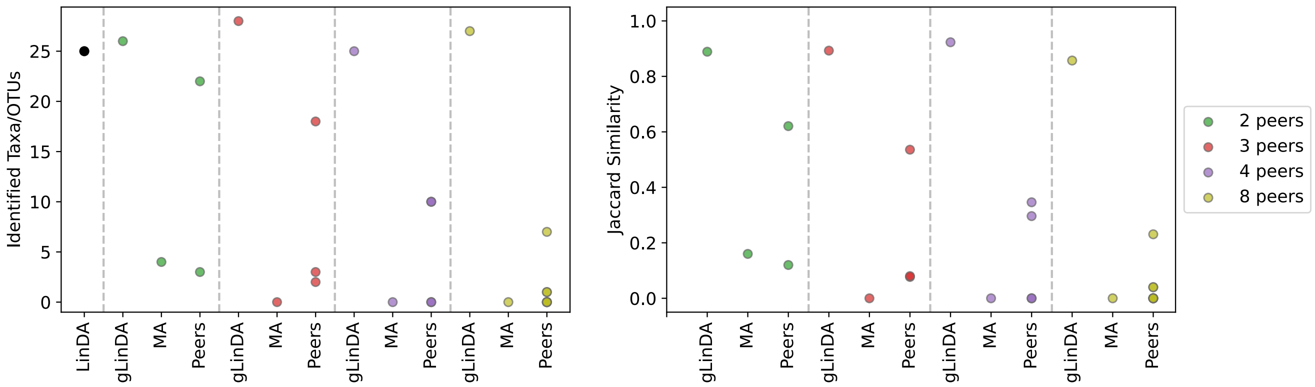


**(a)** Union approach

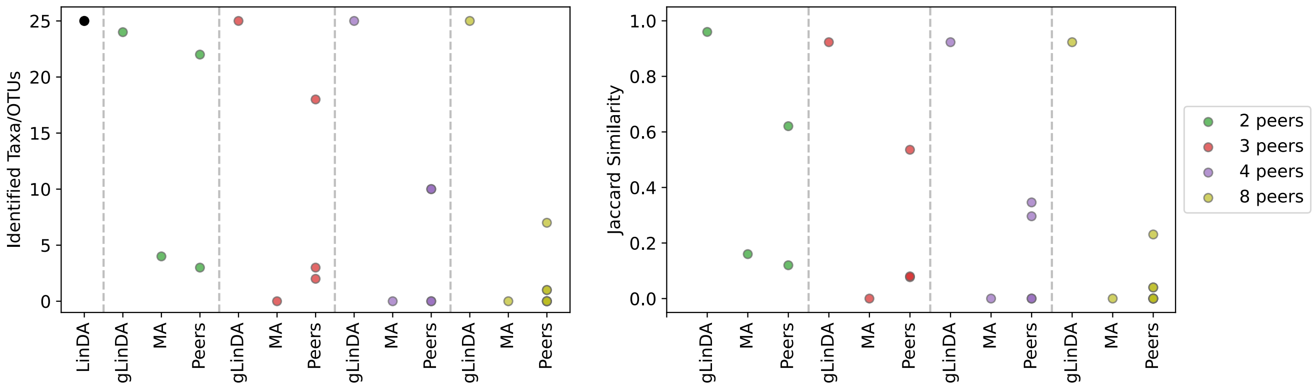


**(b)** Intersection approach

**Fig. S4.** Number of identified taxa/OTUs using LinDA, gLinDA with two, three, four, and eight peers, and meta-analysis, along with the corresponding Jaccard similarity based on LinDA's predictions for the OB-Goodrich dataset.

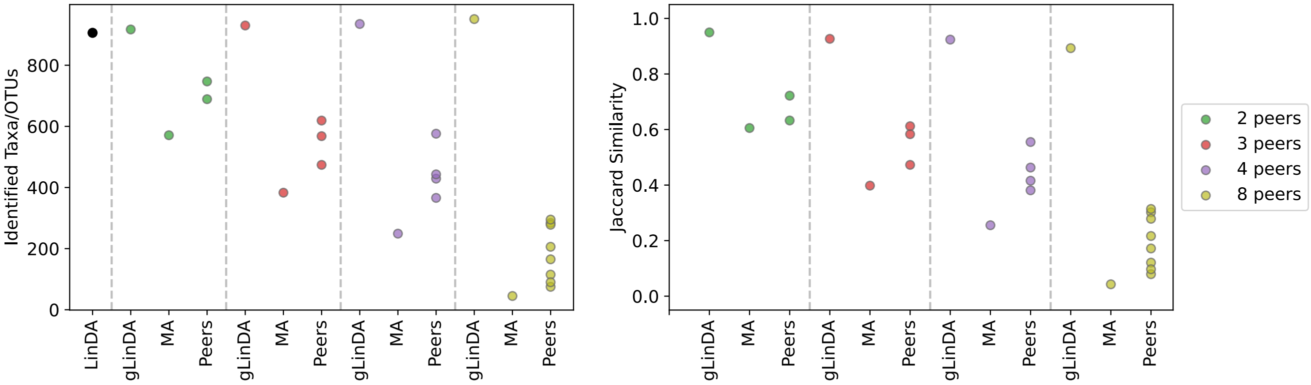


**(a)** Union approach

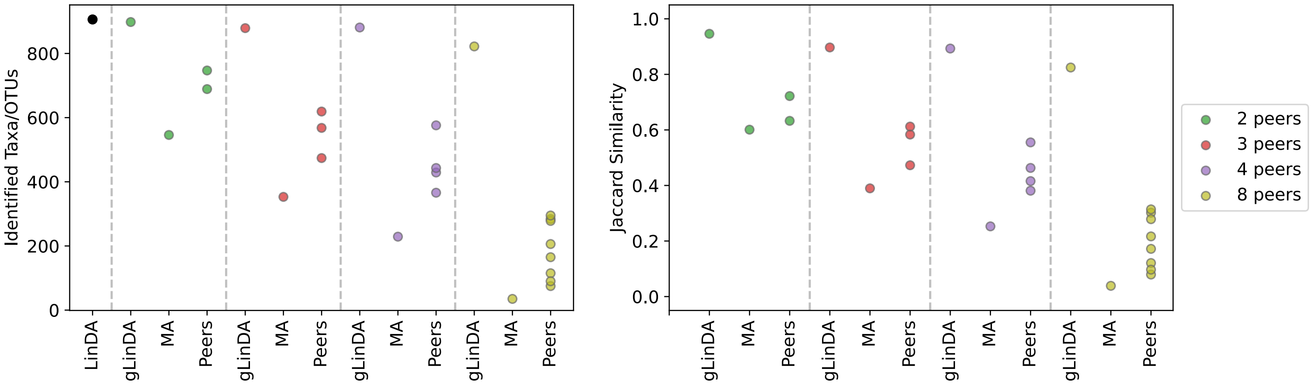


**(b)** Intersection approach

**Fig. S5.** Number of identified taxa/OTUs using LinDA, gLinDA with two, three, four, and eight peers, and meta-analysis, along with the corresponding Jaccard similarity based on LinDA's predictions for the GWMC-Hot-Cold dataset.

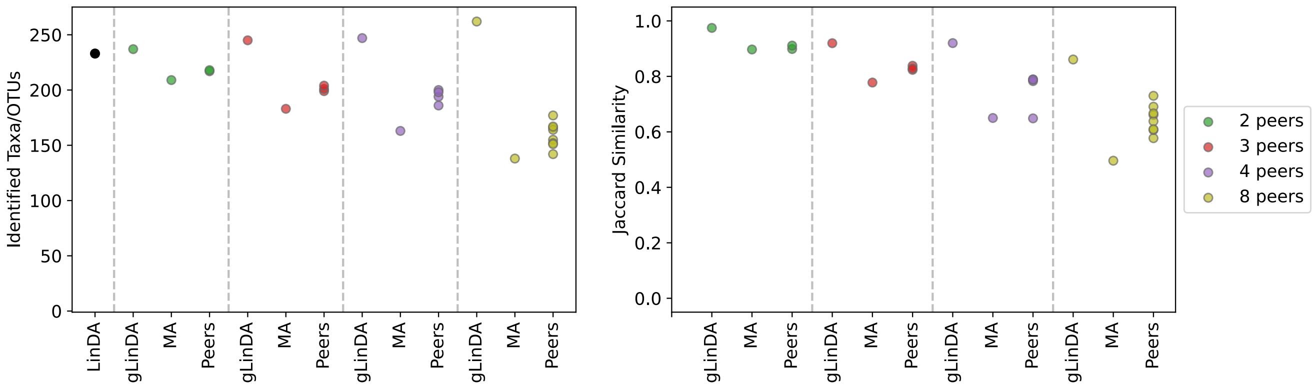


**(a)** Union approach

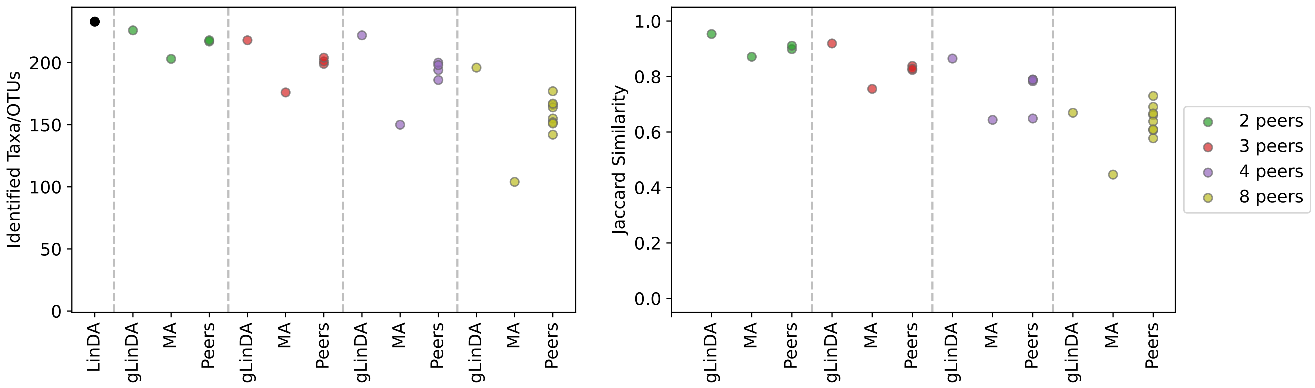


**(b)** Intersection approach

**Fig. S6.** Number of identified taxa/OTUs using LinDA, gLinDA with two, three, four, and eight peers, and meta-analysis, along with the corresponding Jaccard similarity based on LinDA's predictions for the Arctic-Freshwater dataset.

 

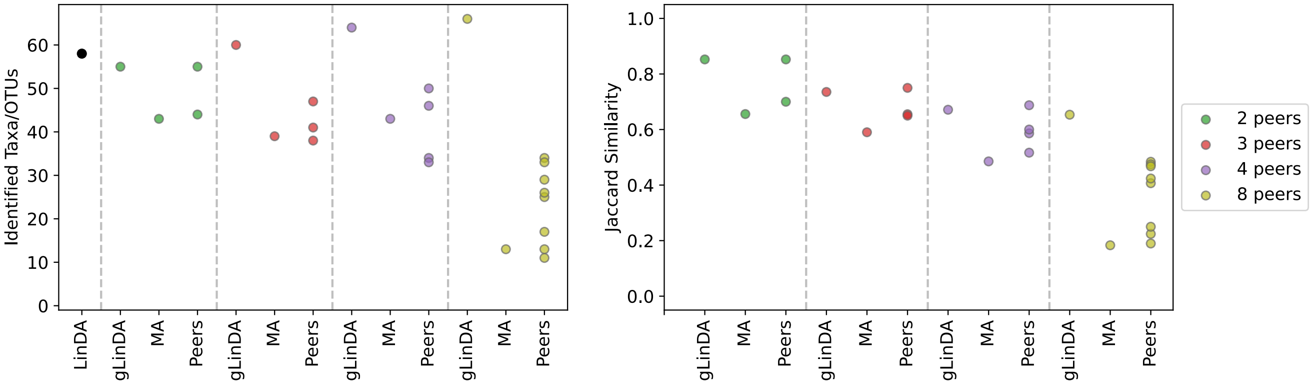


**(a)** Union approach

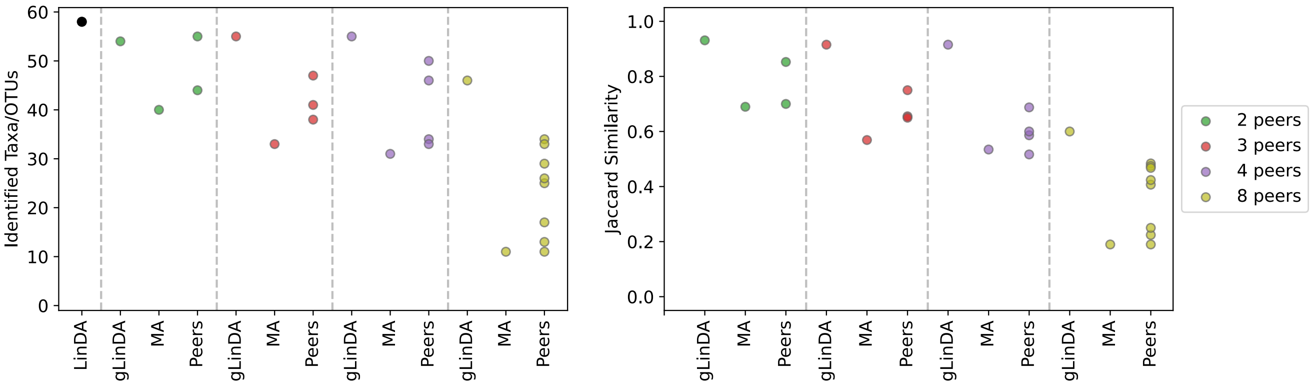


**(b)** Intersection approach

**Fig. S7.** Number of identified taxa/OTUs using LinDA, gLinDA with two, three, four, and eight peers, and meta-analysis, along with the corresponding Jaccard similarity based on LinDA's predictions for the CDI-Shubert dataset.

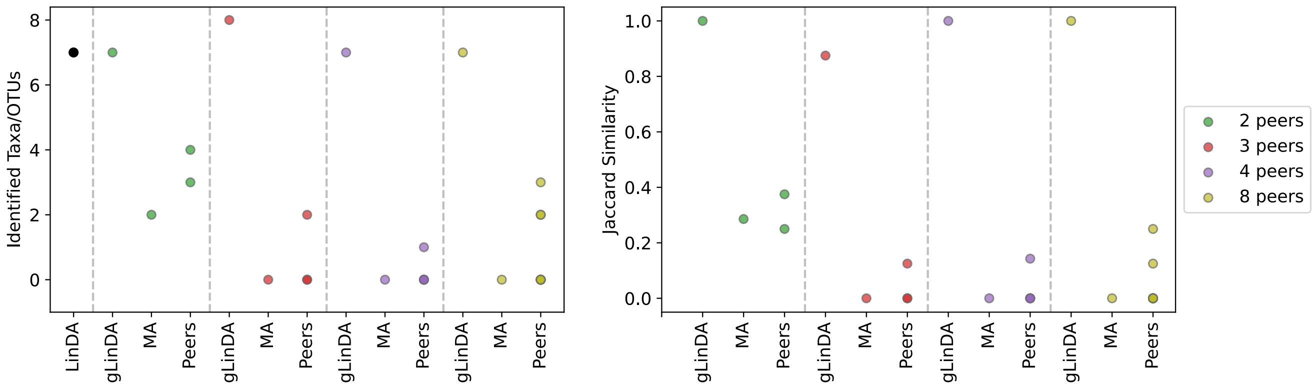


**(a)** Union approach

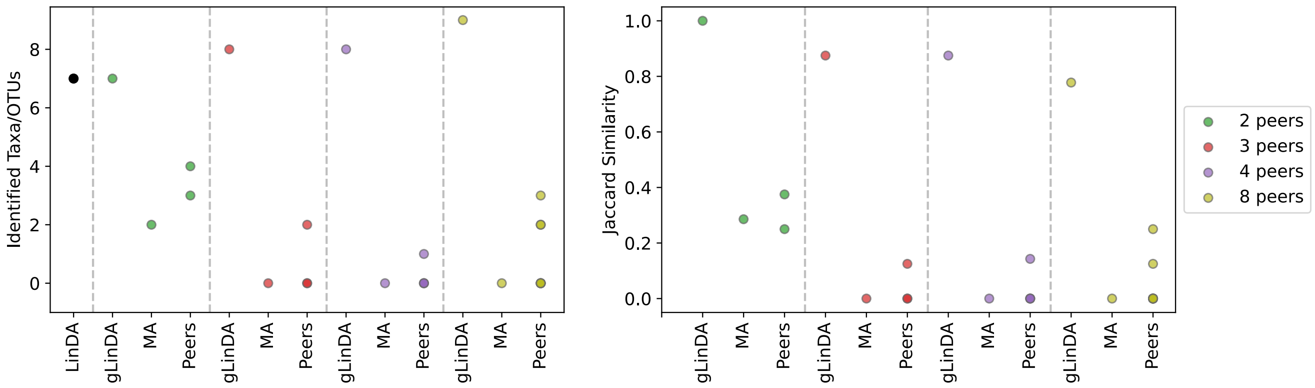


**(b)** Intersection approach

**Fig. S8.** Number of identified taxa/OTUs using LinDA, gLinDA with two, three, four, and eight peers, and meta-analysis, along with the corresponding Jaccard similarity based on LinDA's predictions for the AGP-Diabetes dataset.

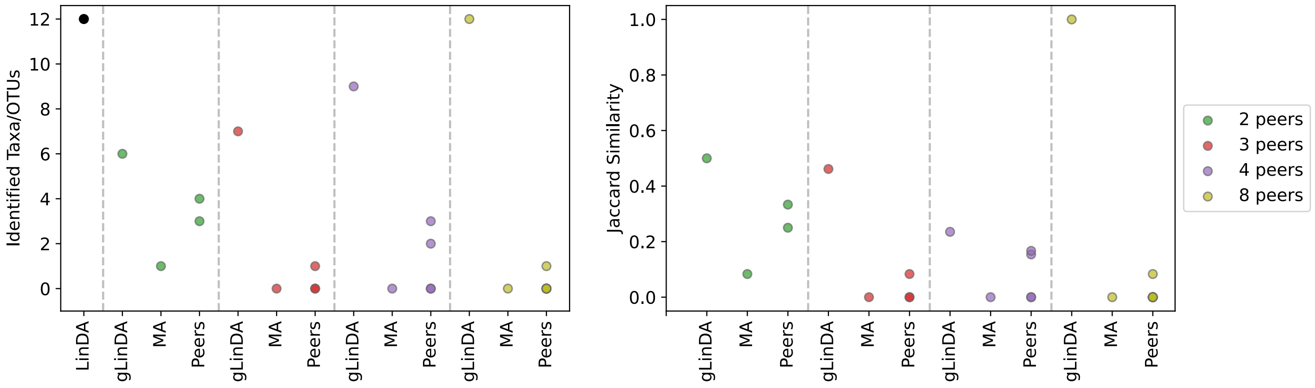


**(a)** Union approach

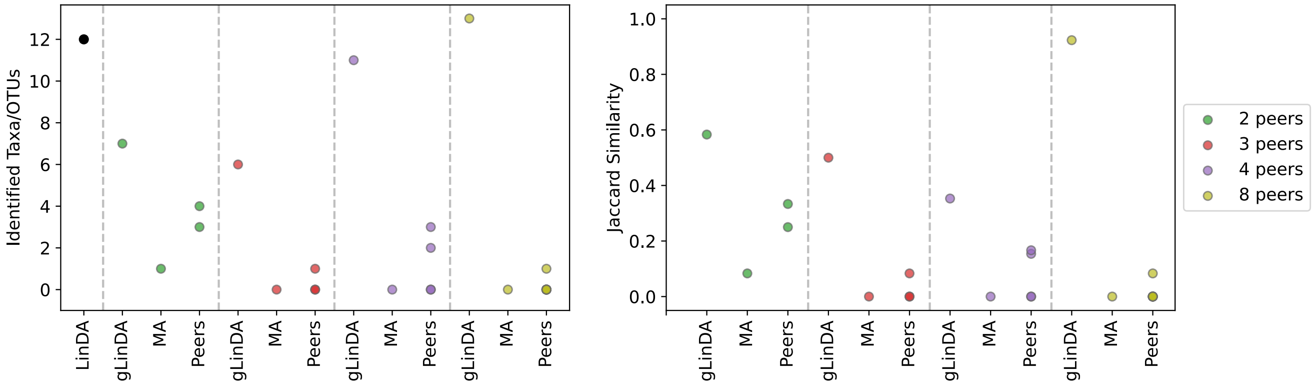


**(b)** Intersection approach

**Fig. S9.** Number of identified taxa/OTUs using LinDA, gLinDA with two, three, four, and eight peers, and meta-analysis, along with the corresponding Jaccard similarity based on LinDA's predictions for the AGP-Ibs dataset.

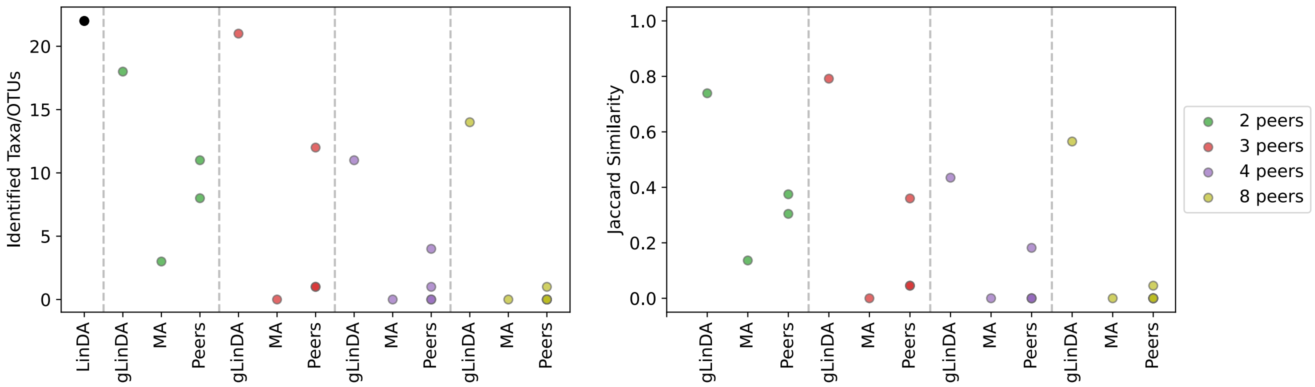


**(a)** Union approach

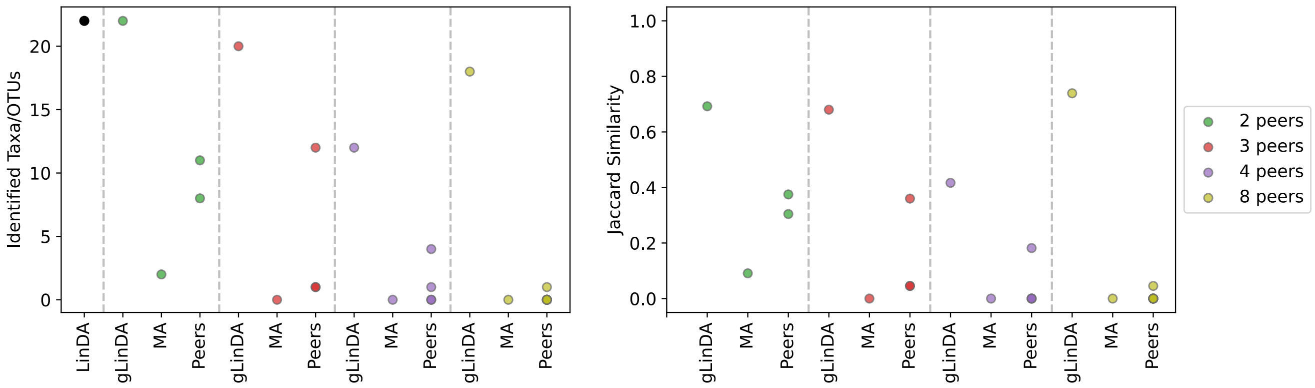


**(b)** Intersection approach

**Fig. S10.** Number of identified taxa/OTUs using LinDA, gLinDA with two, three, four, and eight peers, and meta-analysis, along with the corresponding Jaccard similarity based on LinDA's predictions for the Smoke dataset.

 Leon Fehse, Mohammad Tajabadi, Roman Martin, Hajo Holzmann, Dominik Heider

**Fig. S11.** P-values for the S-5000 dataset for two peers with equal data distribution.

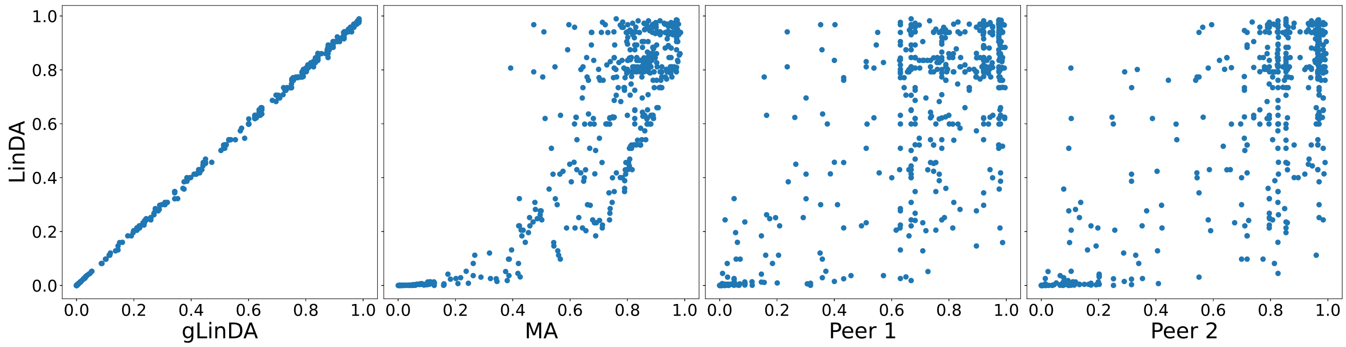


**(a)** Union approach

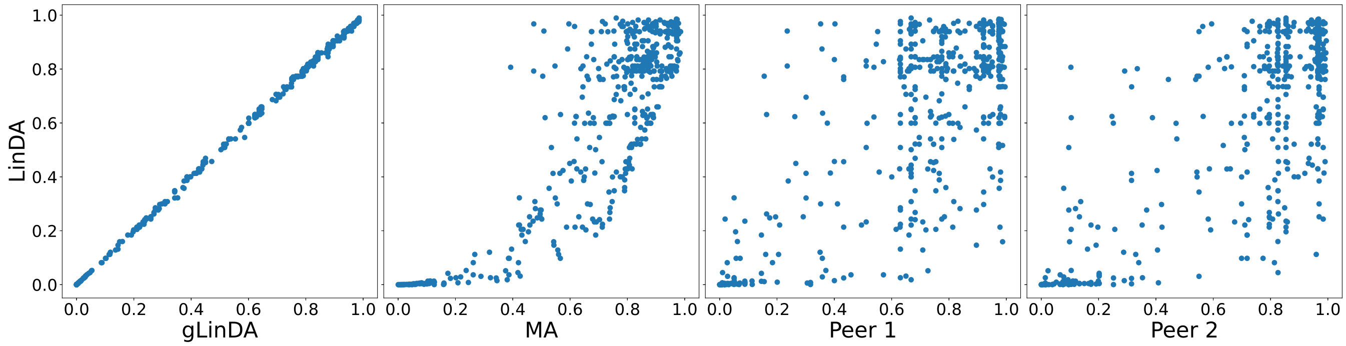


**(b)** Intersection approach

**Fig. S12.** P-values for the OB-Goodrich dataset for two peers with equal data distribution.

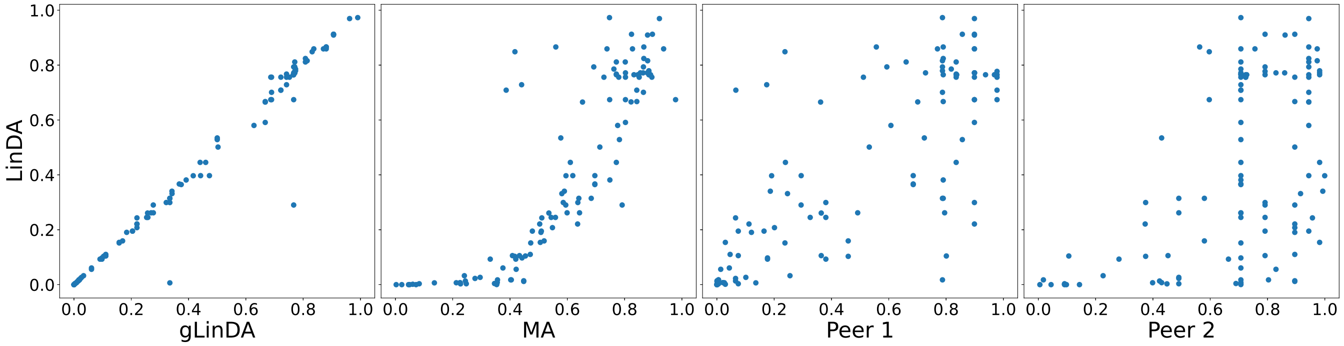


**(a)** Union approach

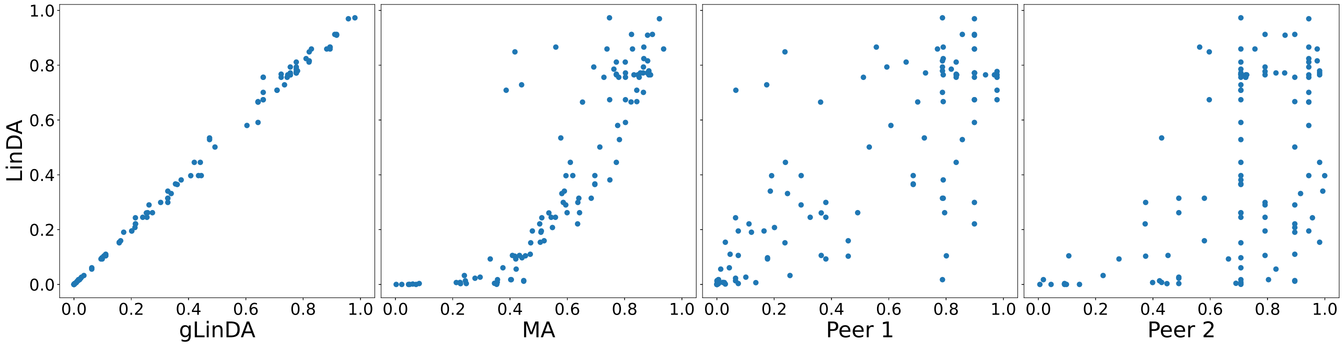


**(b)** Intersection approach

**Fig. S13.** P-values for the GWMC-Hot-Cold dataset for two peers with equal data distribution.

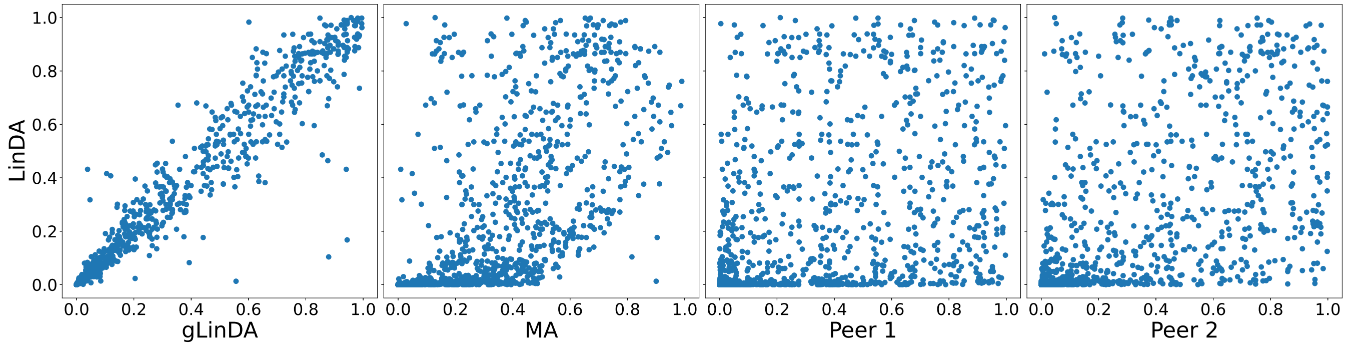


**(a)** Union approach

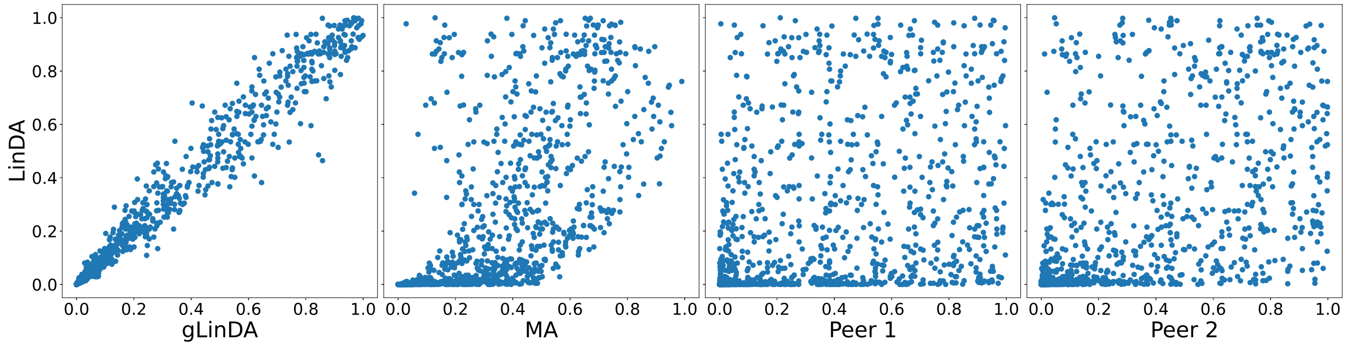


**(b)** Intersection approach

**Fig. S14.** P-values for the Arctic-Freshwater dataset for two peers with equal data distribution.

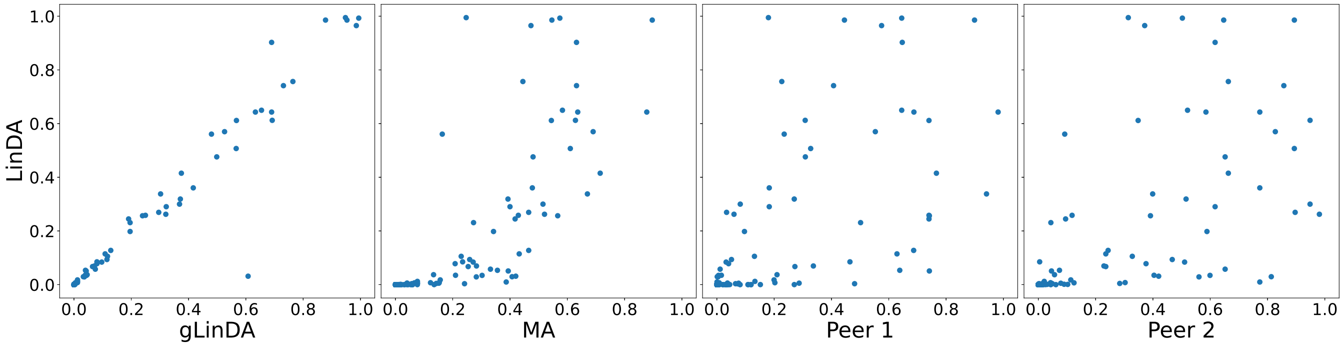


**(a)** Union approach

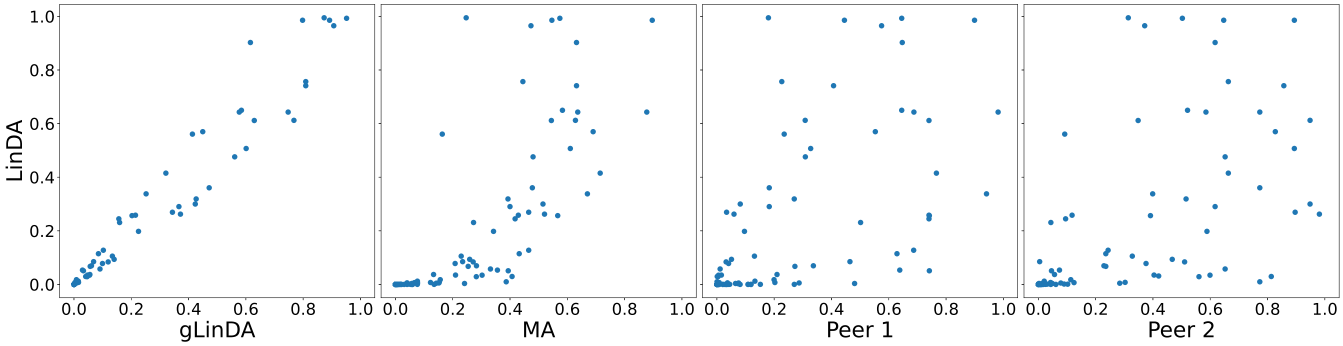


**(b)** Intersection approach

**Fig. S15.** P-values for the CDI-Shubert dataset for two peers with equal data distribution.

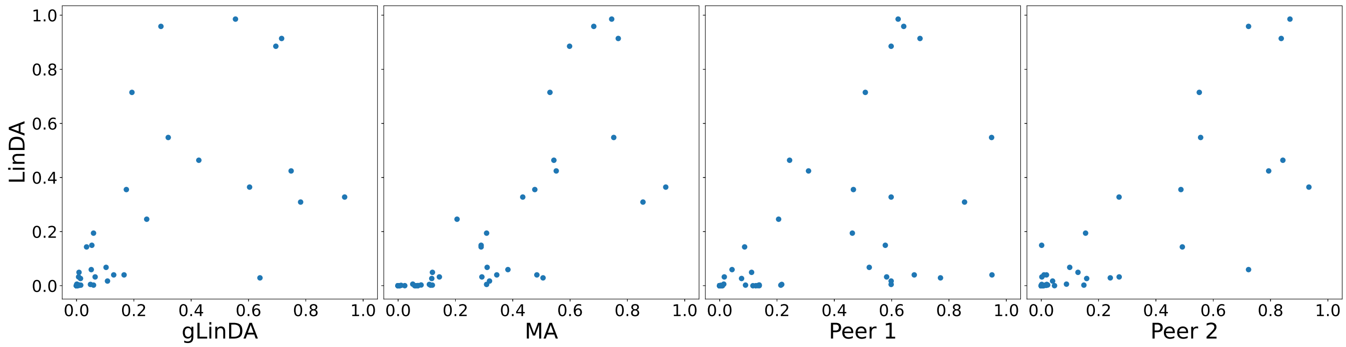


**(a)** Union approach

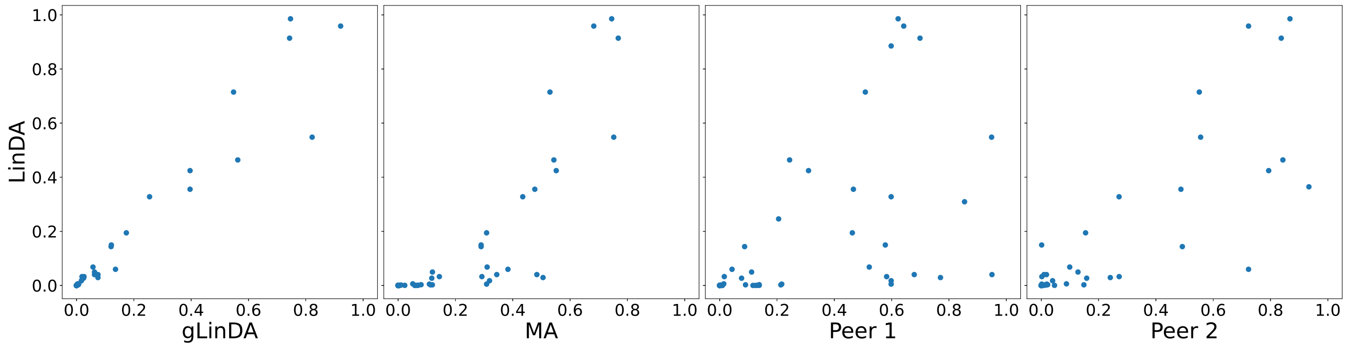


**(b)** Intersection approach

**Fig. S16.** P-values for the AGP-Diabetes dataset for two peers with equal data distribution.

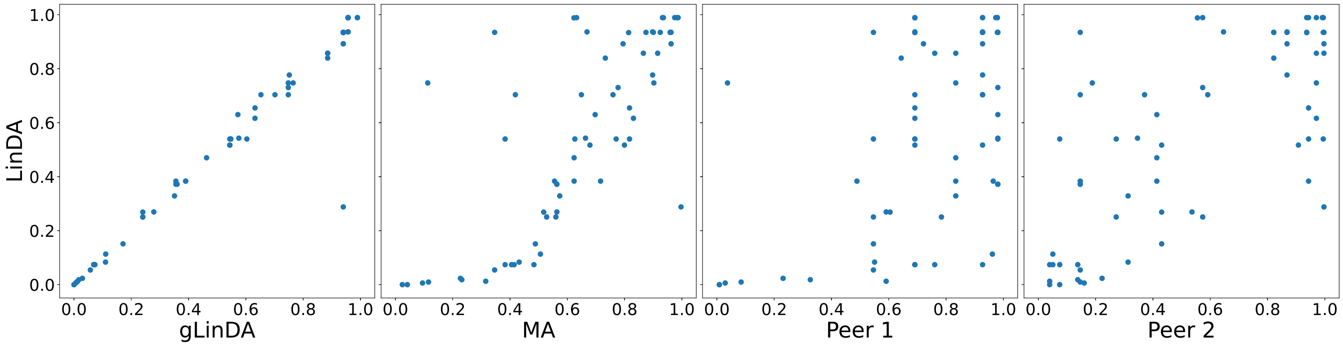


**(a)** Union approach

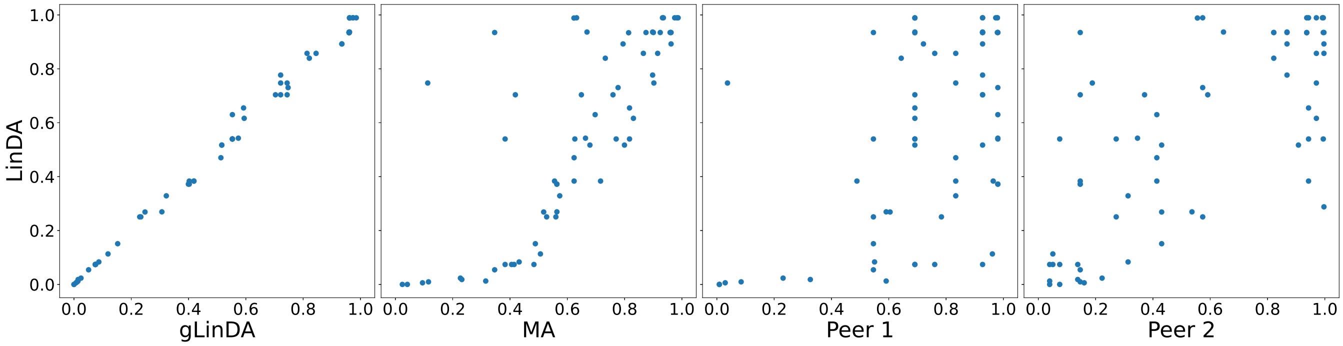


**(b)** Intersection approach

**Fig. S17.** P-values for the AGP-lbs dataset for two peers with equal data distribution.

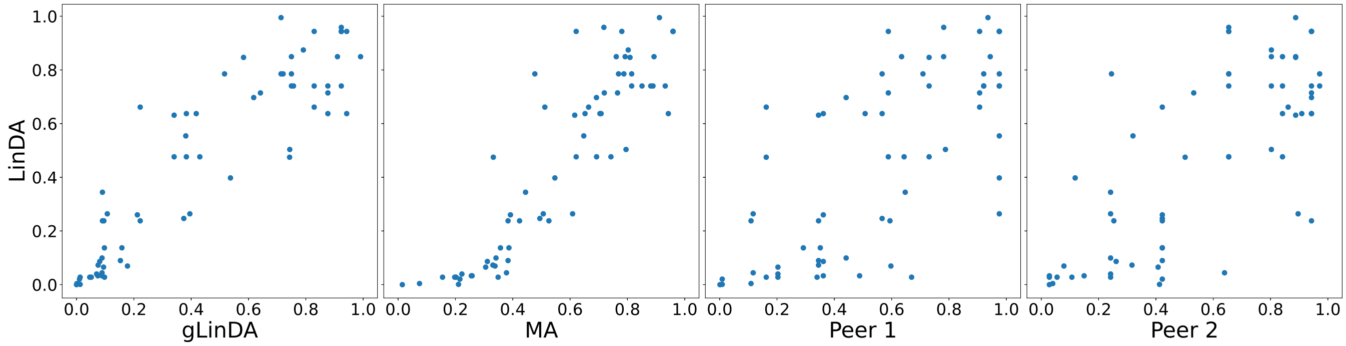


**(a)** Union approach

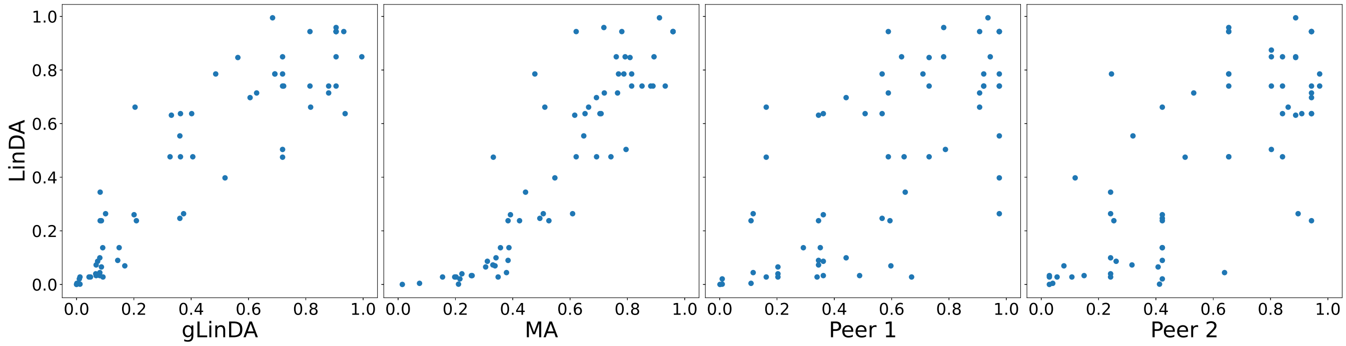


**(b)** Intersection approach

**Fig. S18.** P-values for the Smoke dataset for two peers with equal data distribution.

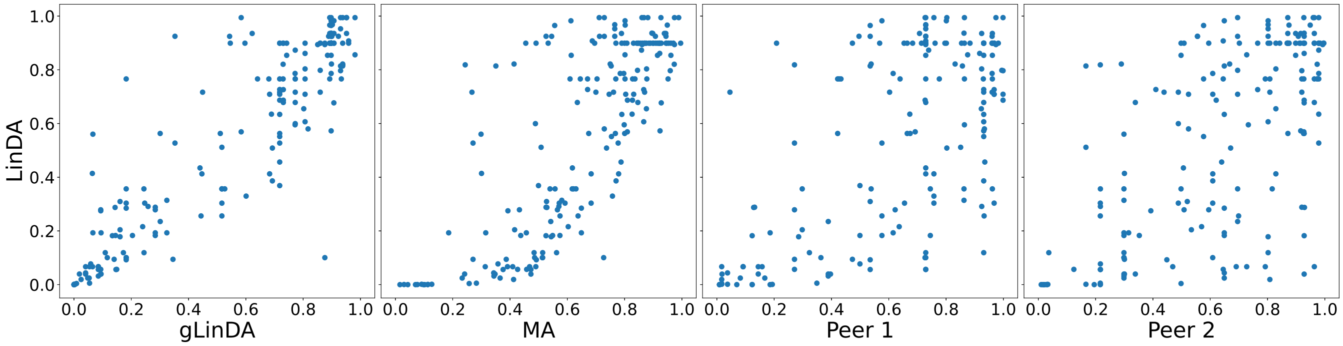


**(a)** Union approach

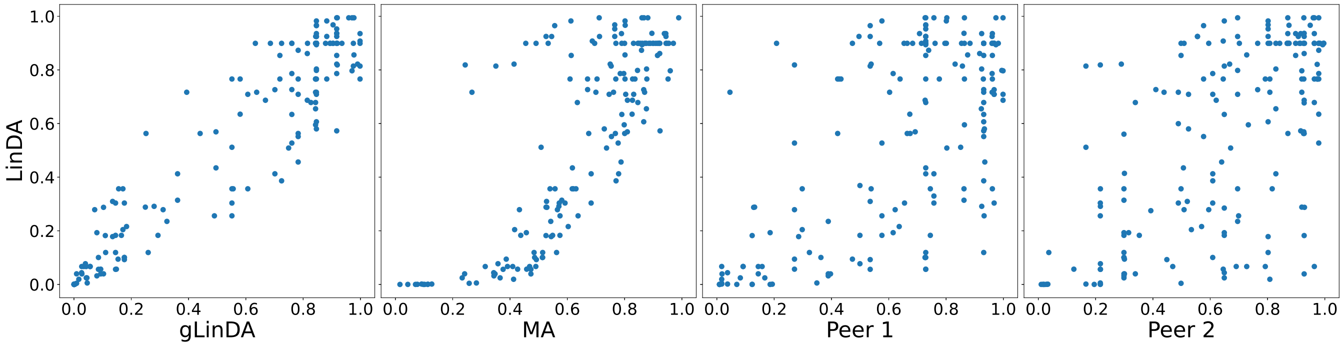


**(b)** Intersection approach

**Fig. S19.** P-values for the S-5000 dataset for three peers with equal data distribution.

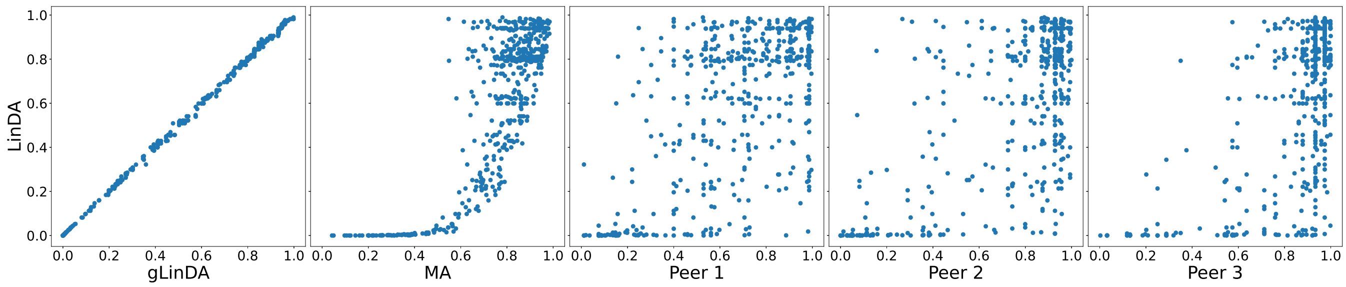


**(a)** Union approach

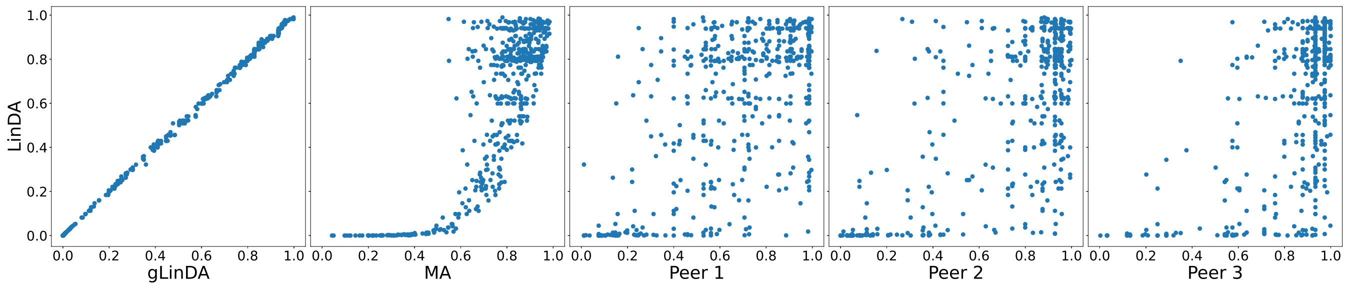


**(b)** Intersection approach

**Fig. S20.** P-values for the OB-Goodrich dataset for three peers with equal data distribution.

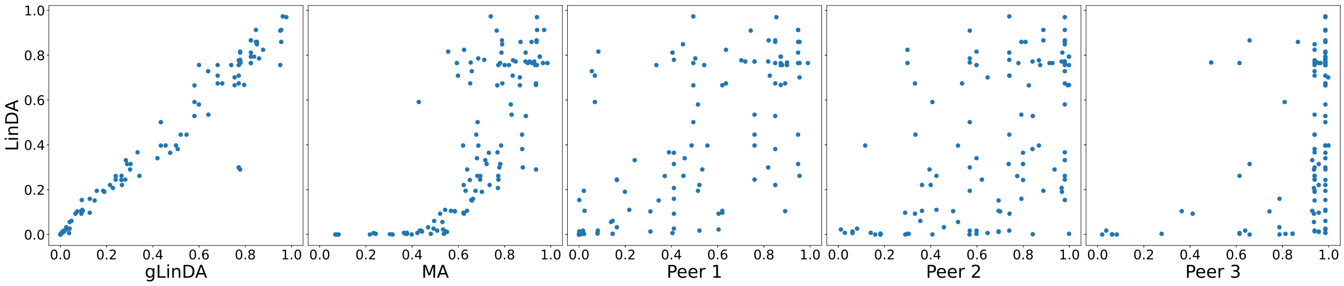


**(a)** Union approach

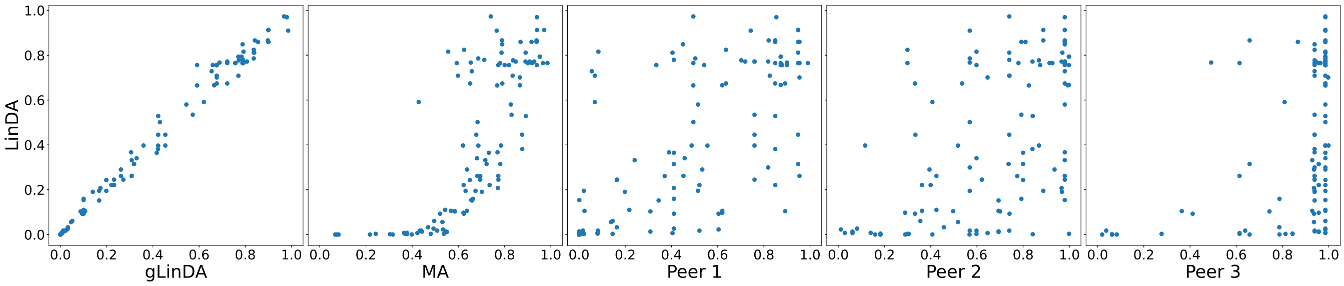


**(b)** Intersection approach

 Leon Fehse, Mohammad Tajabadi, Roman Martin, Hajo Holzmann, Dominik Heider

**Fig. S21.** P-values for the GWMC-Hot-Cold dataset for three peers with equal data distribution.

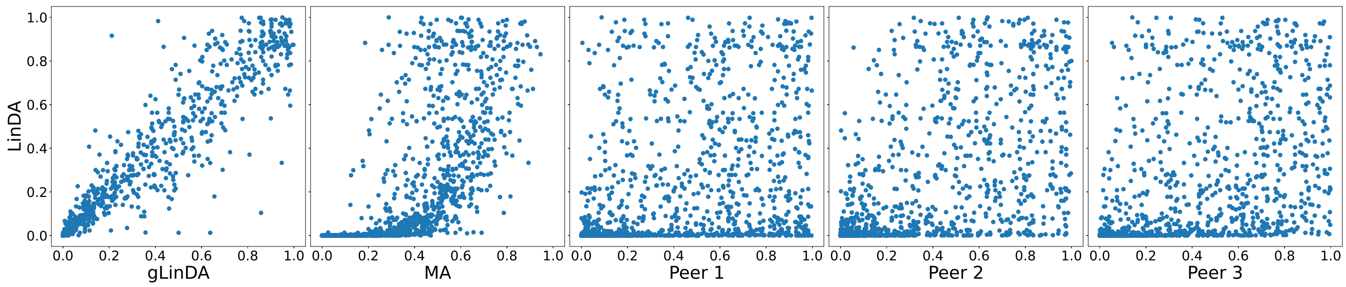


**(a)** Union approach

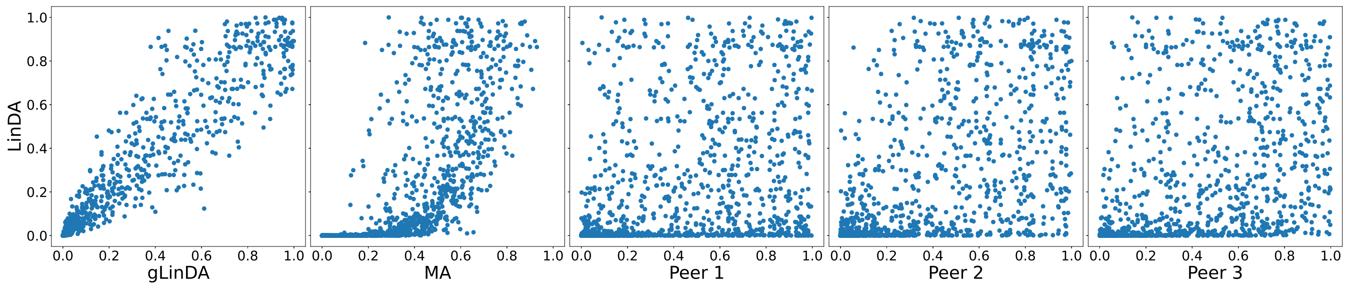


**(b)** Intersection approach

**Fig. S22.** P-values for the Arctic-Freshwater dataset for three peers with equal data distribution.

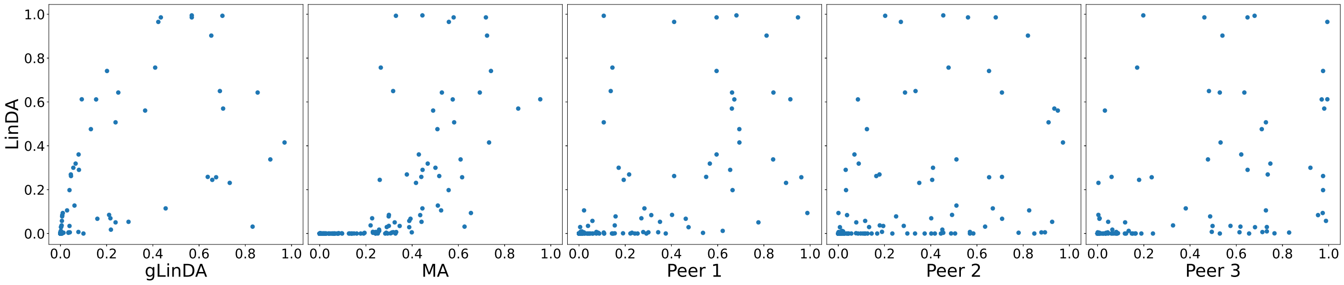


**(a)** Union approach

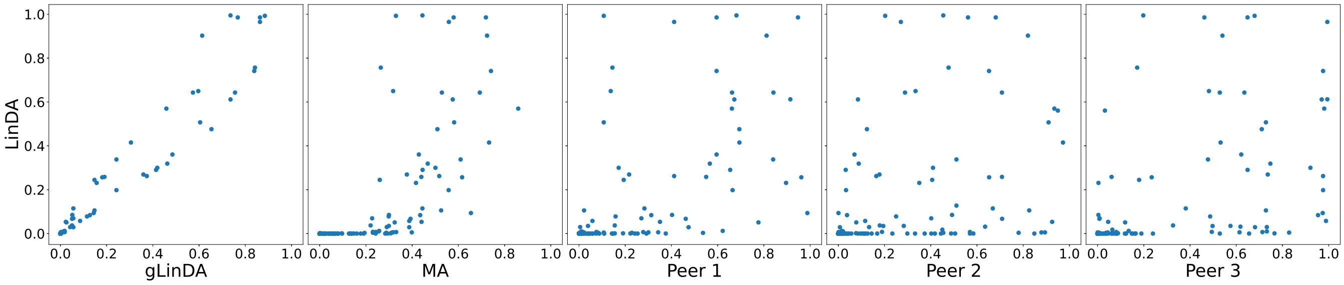


**(b)** Intersection approach

**Fig. S23.** P-values for the CDI-Shubert dataset for three peers with equal data distribution.

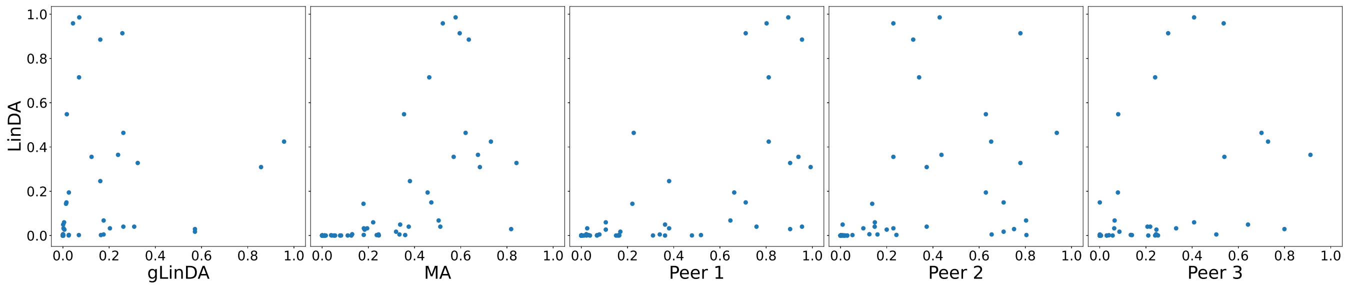


**(a)** Union approach

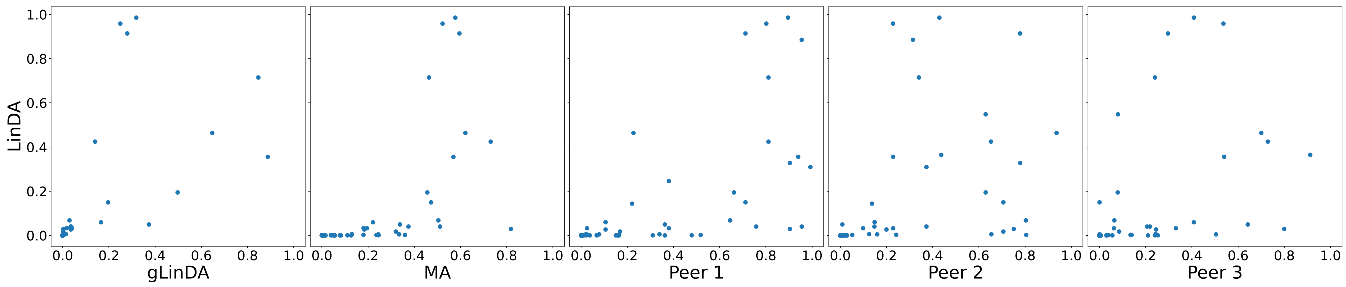


**(b)** Intersection approach

**Fig. S24.** P-values for the AGP-Diabetes dataset for three peers with equal data distribution.

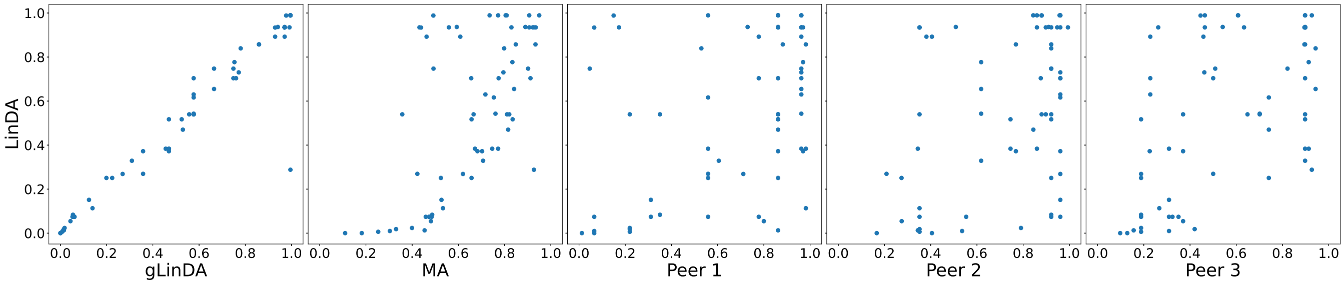


**(a)** Union approach

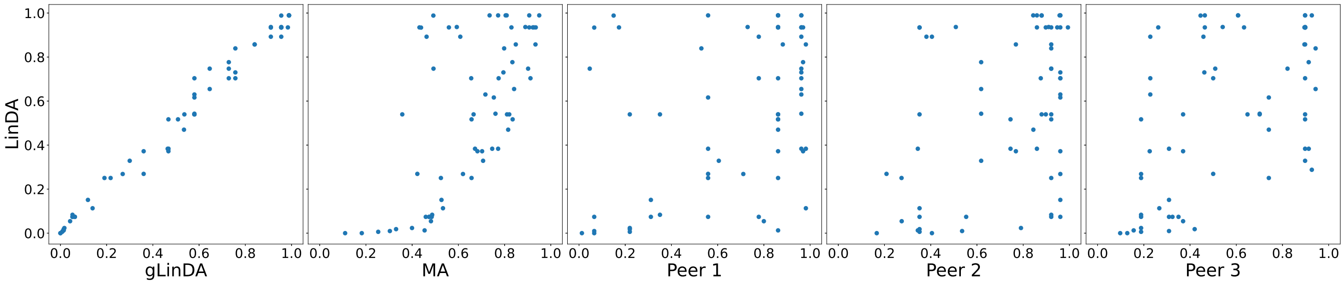


**(b)** Intersection approach

**Fig. S25.** P-values for the AGP-lbs dataset for three peers with equal data distribution.

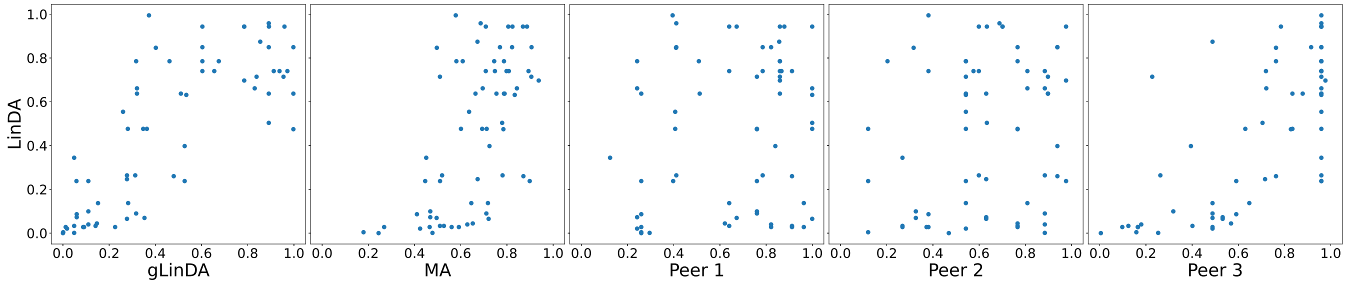


**(a)** Union approach

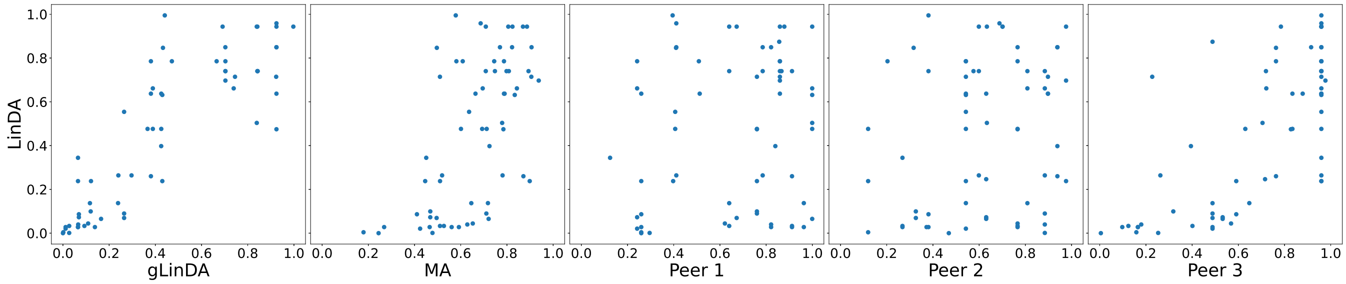


**(b)** Intersection approach

**Fig. S26.** P-values for the Smoke dataset for three peers with equal data distribution.

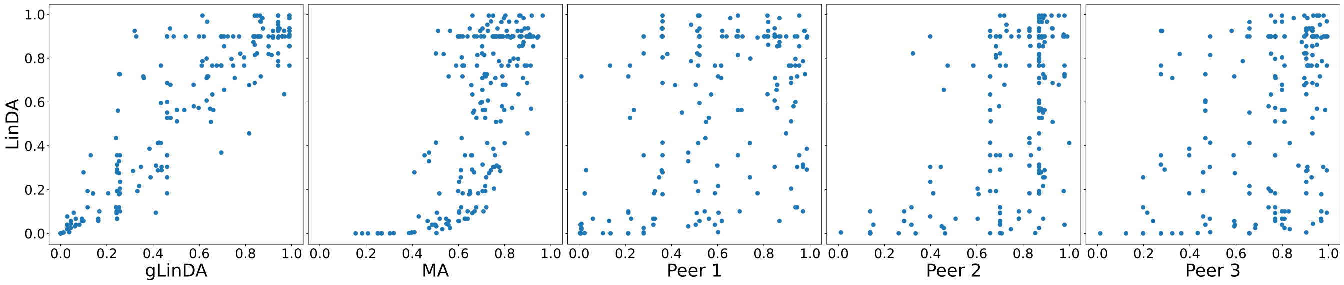


**(a)** Union approach

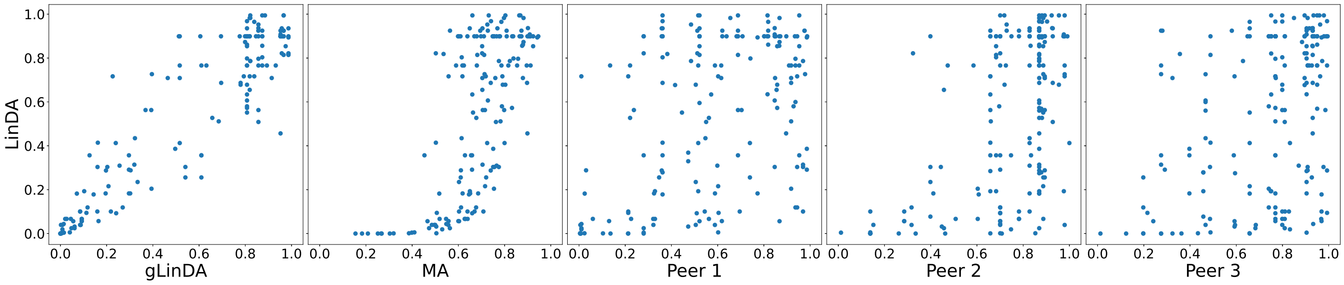


**(b)** Intersection approach

**Fig. S27.** P-values for the S-5000 dataset for three peers with unequal data distribution with ratios 0.1, 0.3, and 0.6.

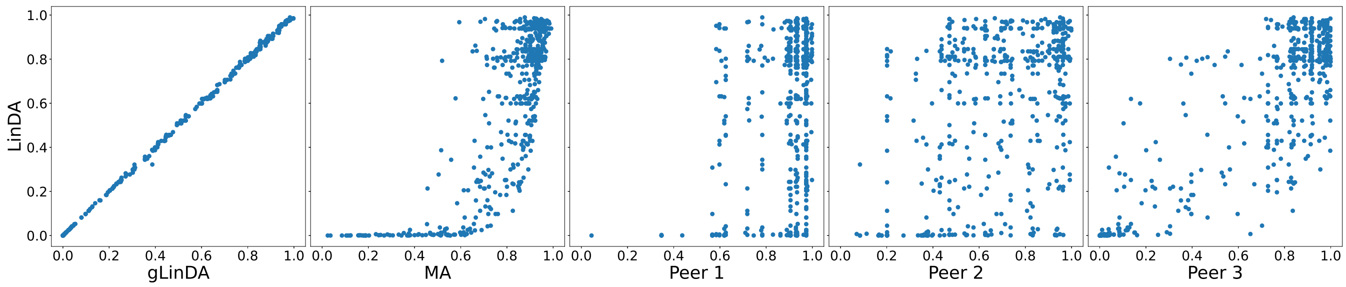


**(a)** Union approach

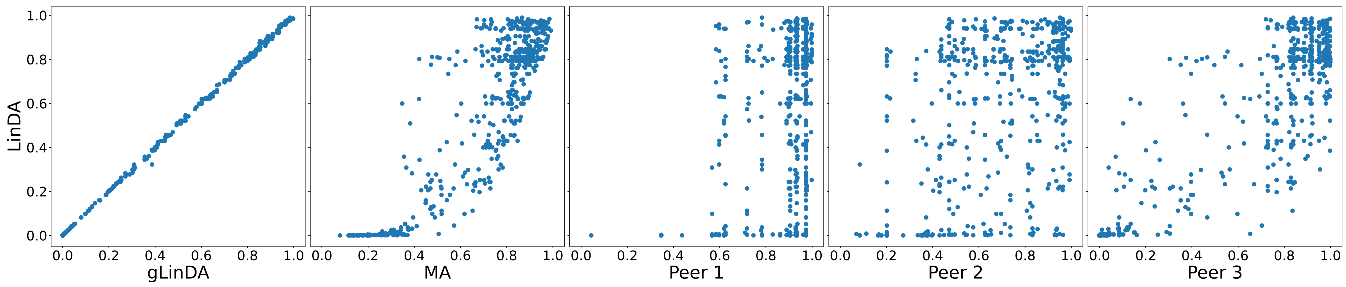


**(b)** Intersection approach

**Fig. S28.** P-values for the OB-Goodrich dataset for three peers with unequal data distribution with ratios 0.1, 0.3, and 0.6.

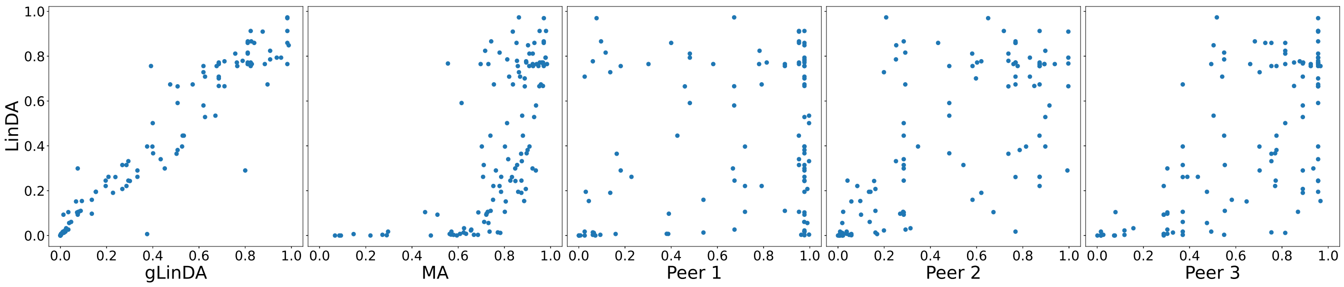


**(a)** Union approach

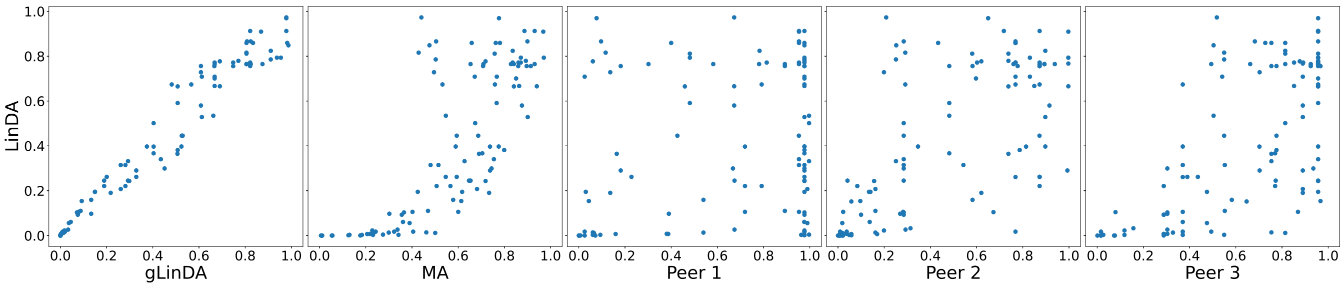


**(b)** Intersection approach

**Fig. S29.** P-values for the GWMC-Hot-Cold dataset for three peers with unequal data distribution with ratios 0.1, 0.3, and 0.6.

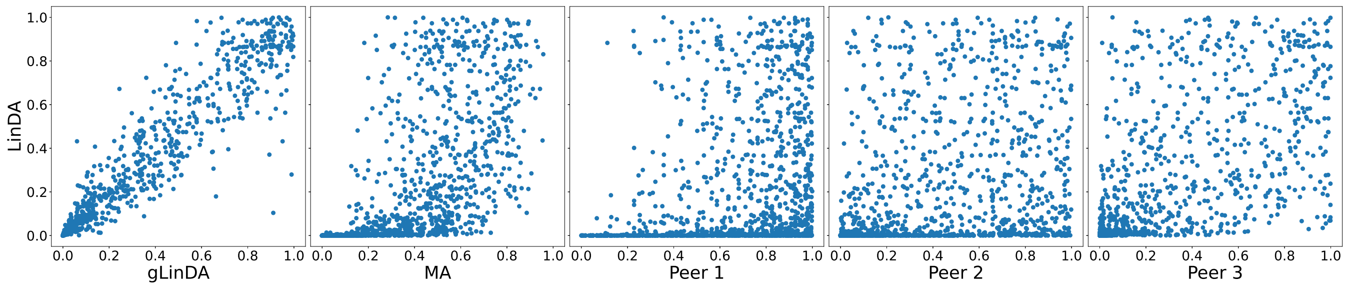


**(a)** Union approach

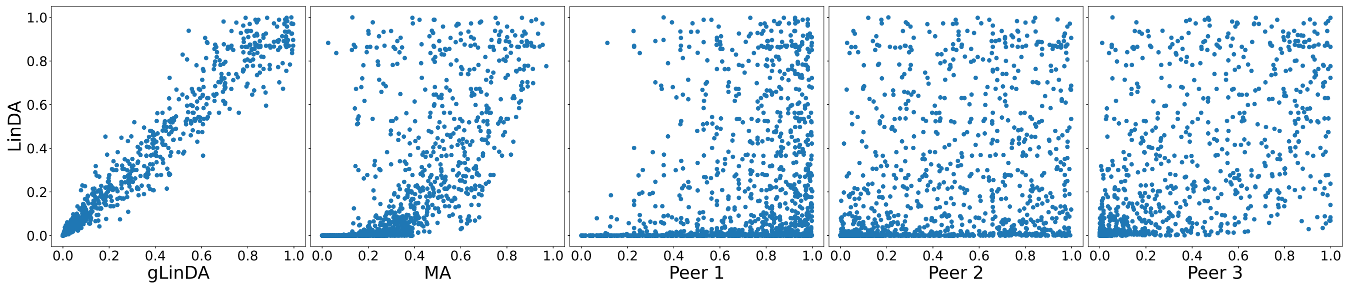


**(b)** Intersection approach

**Fig. S30.** P-values for the Arctic-Freshwater dataset for three peers with unequal data distribution with ratios 0.1, 0.3, and 0.6.

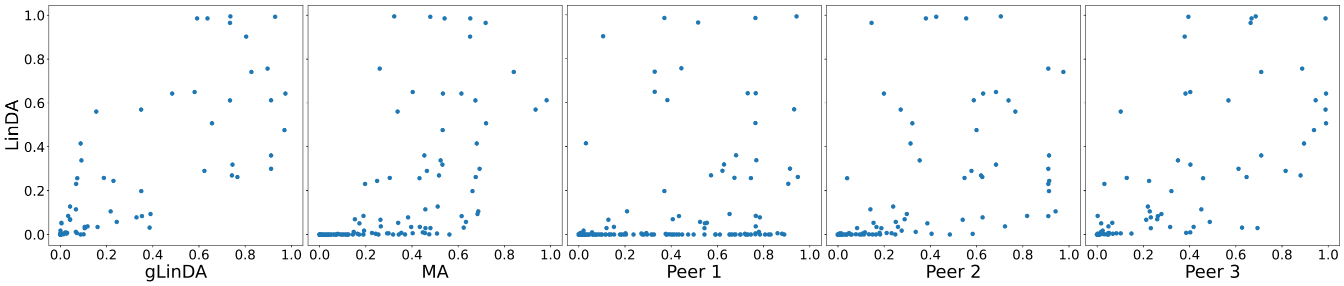


**(a)** Union approach

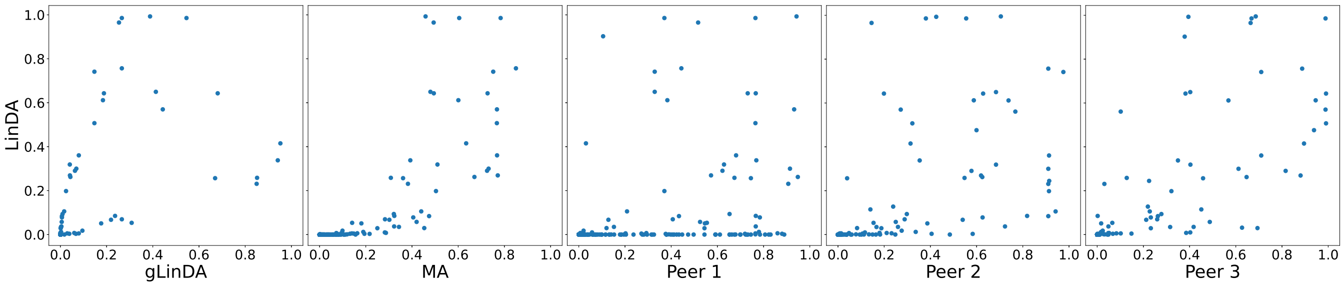


**(b)** Intersection approach

**Fig. S31.** P-values for the CDI-Shubert dataset for three peers with unequal data distribution with ratios 0.1, 0.3, and 0.6.

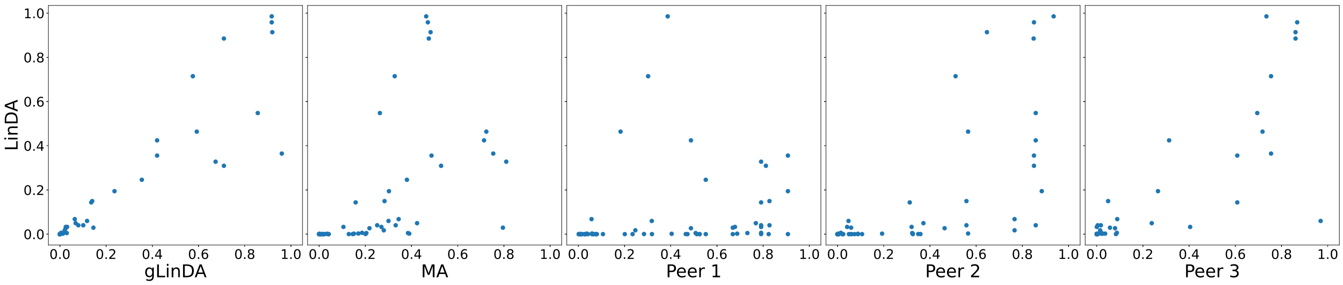


**(a)** Union approach

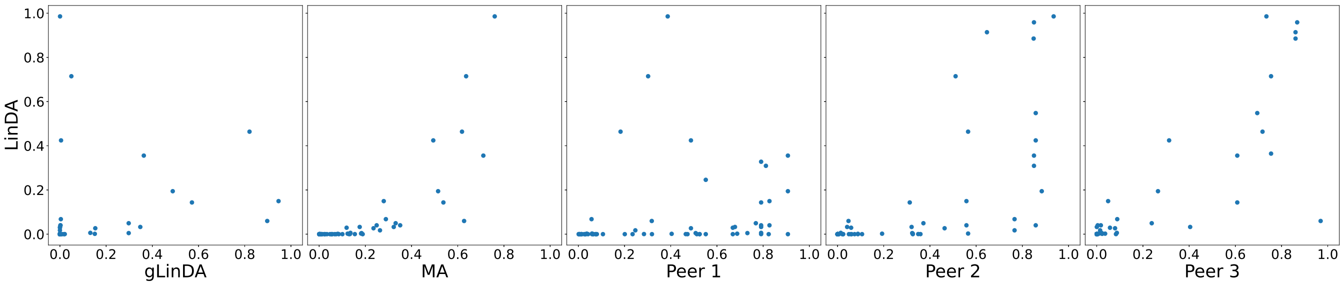


**(b)** Intersection approach

**Fig. S32.** P-values for the AGP-Diabetes dataset for three peers with unequal data distribution with ratios 0.1, 0.3, and 0.6.

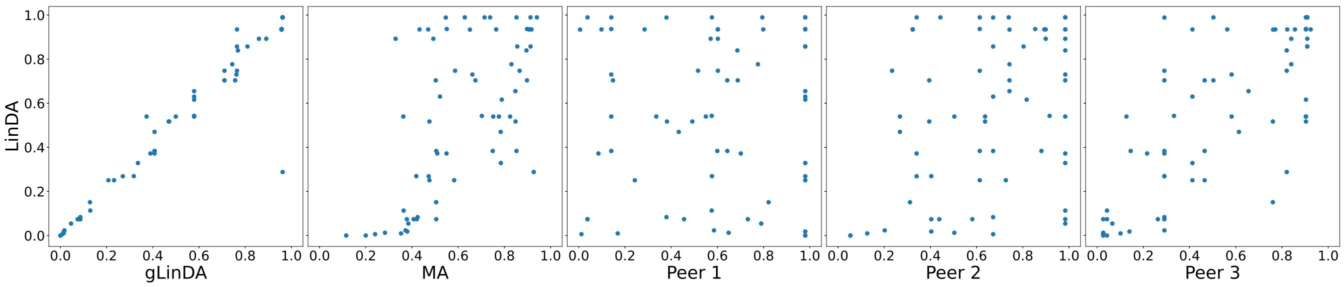


**(a)** Union approach

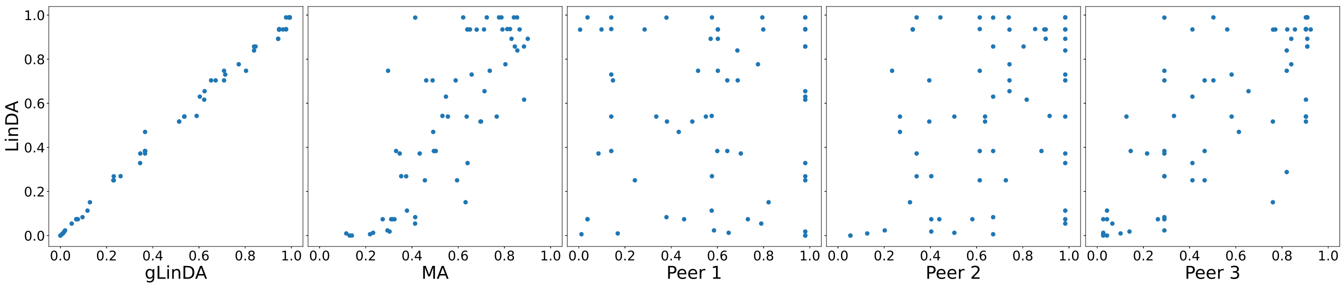


**(b)** Intersection approach

**Fig. S33.** P-values for the AGP-lbs dataset for three peers with unequal data distribution with ratios 0.1, 0.3, and 0.6.

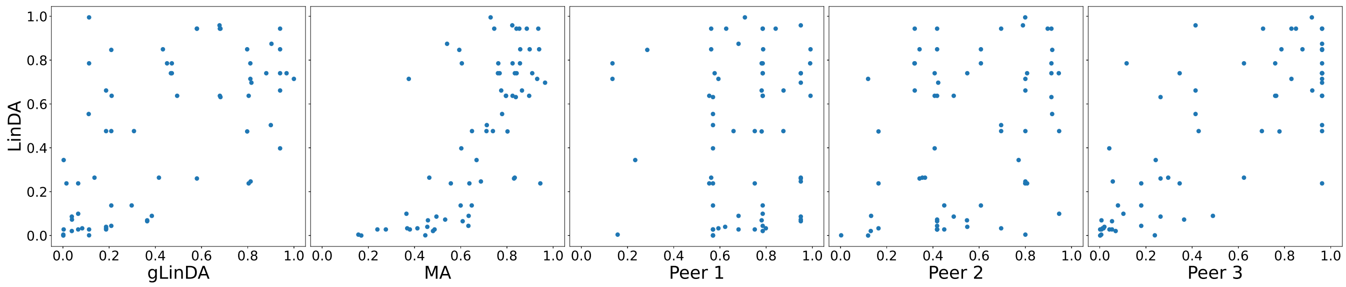


**(a)** Union approach

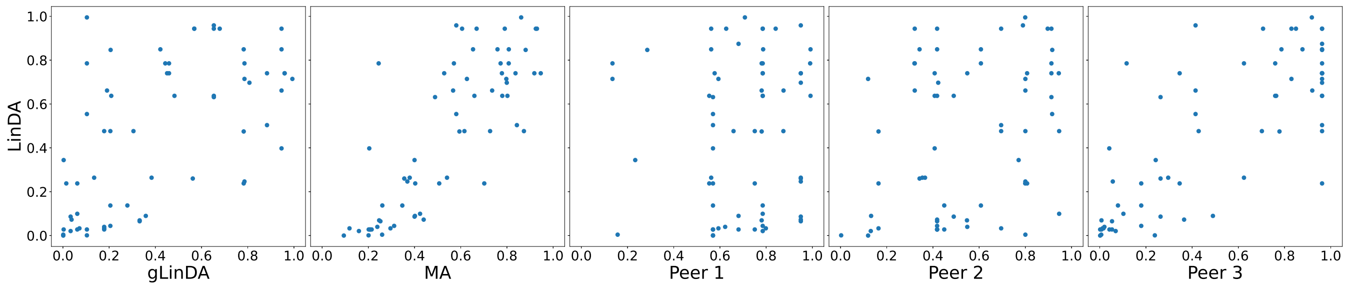


**(b)** Intersection approach

**Fig. S34.** P-values for the S-5000 dataset for four peers with equal data distribution with the union approach.

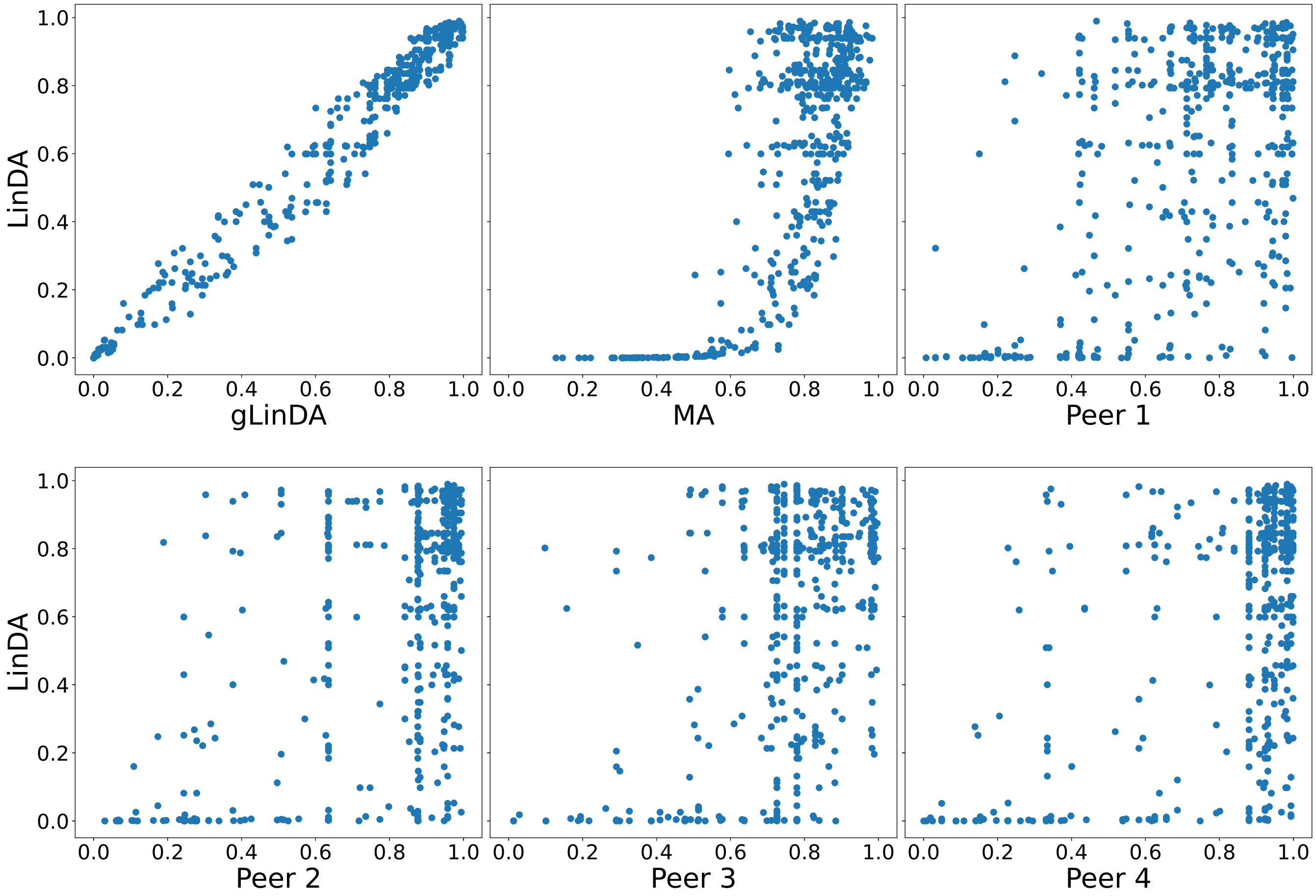


 Leon Fehse, Mohammad Tajabadi, Roman Martin, Hajo Holzmann, Dominik Heider

**Fig. S35.** P-values for the S-5000 dataset for four peers with equal data distribution with the intersection approach.

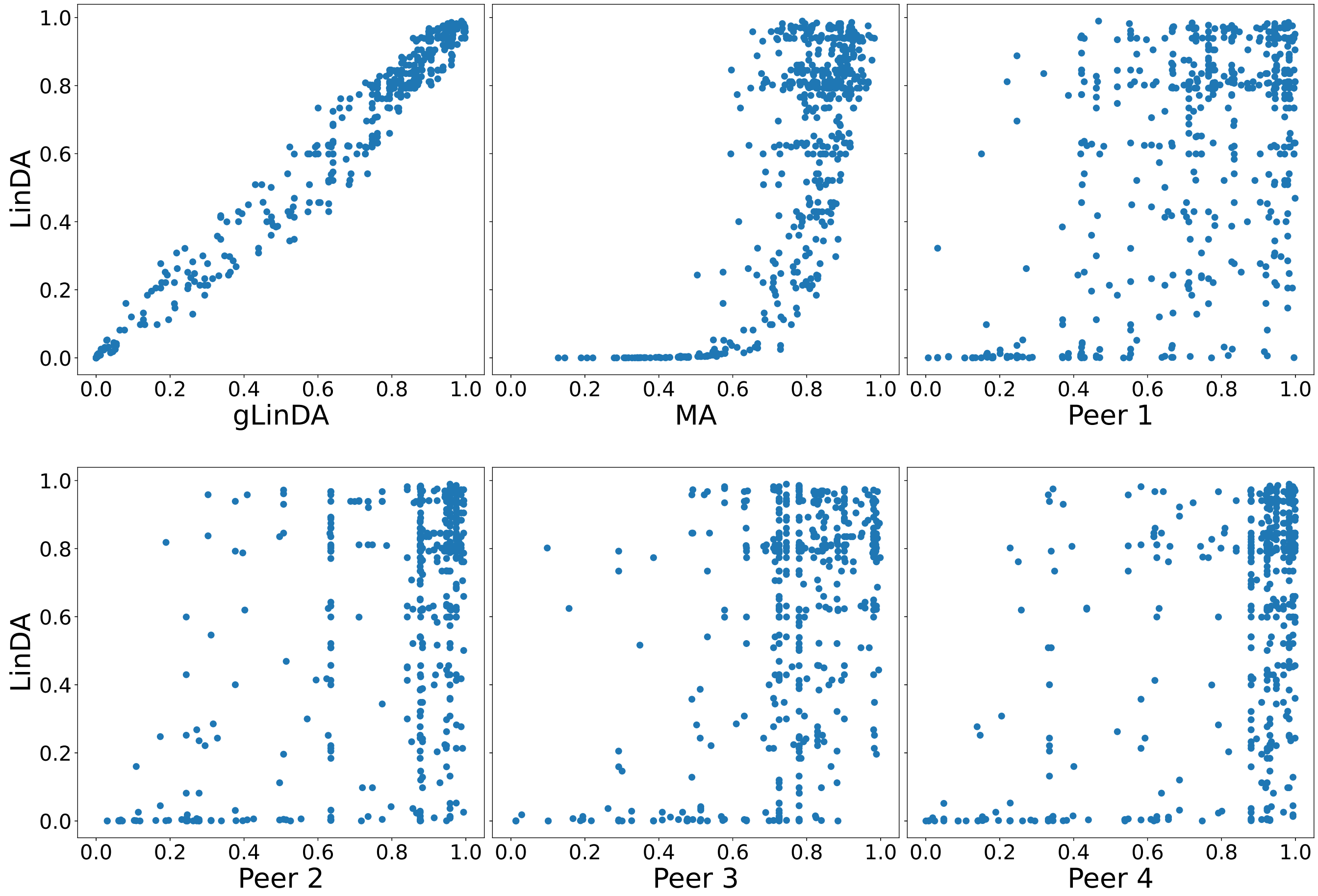

**Fig. S36.** P-values for the OB-Goodrich dataset for four peers with equal data distribution with the union approach.

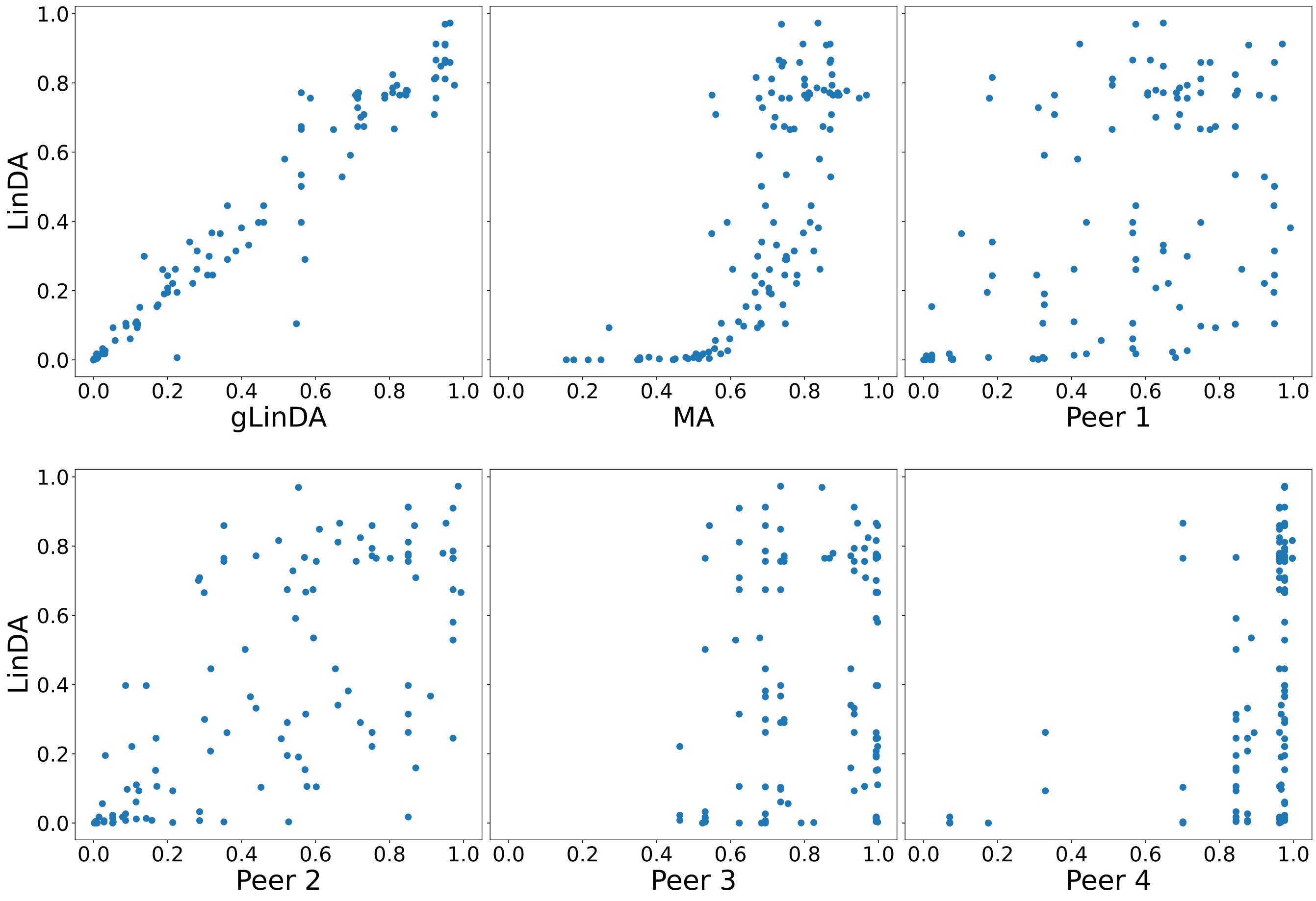

**Fig. S37.** P-values for the OB-Goodrich dataset for four peers with equal data distribution with the intersection approach.

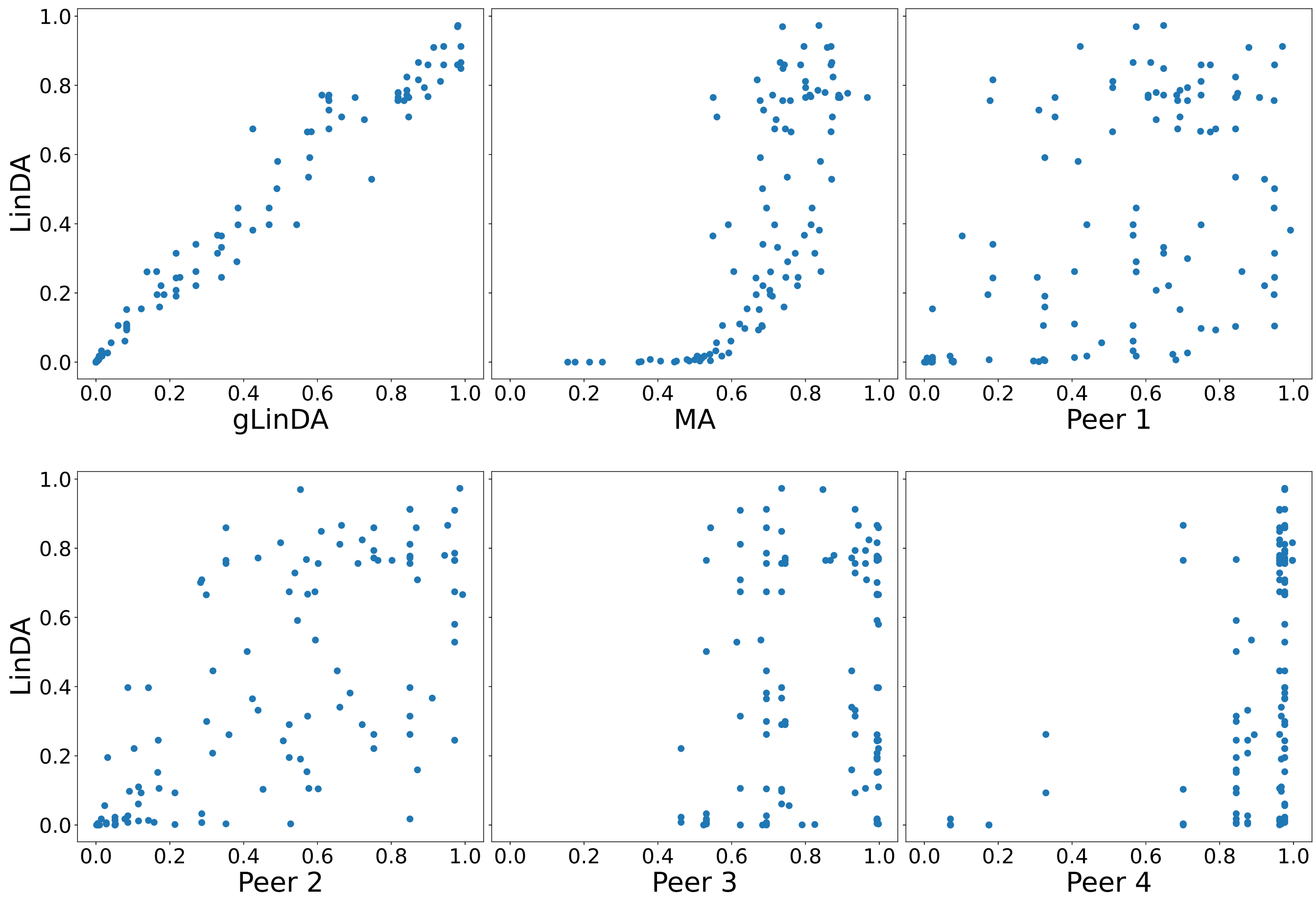

**Fig. S38.** P-values for the GWMC-Hot-Cold dataset for four peers with equal data distribution with the union approach.

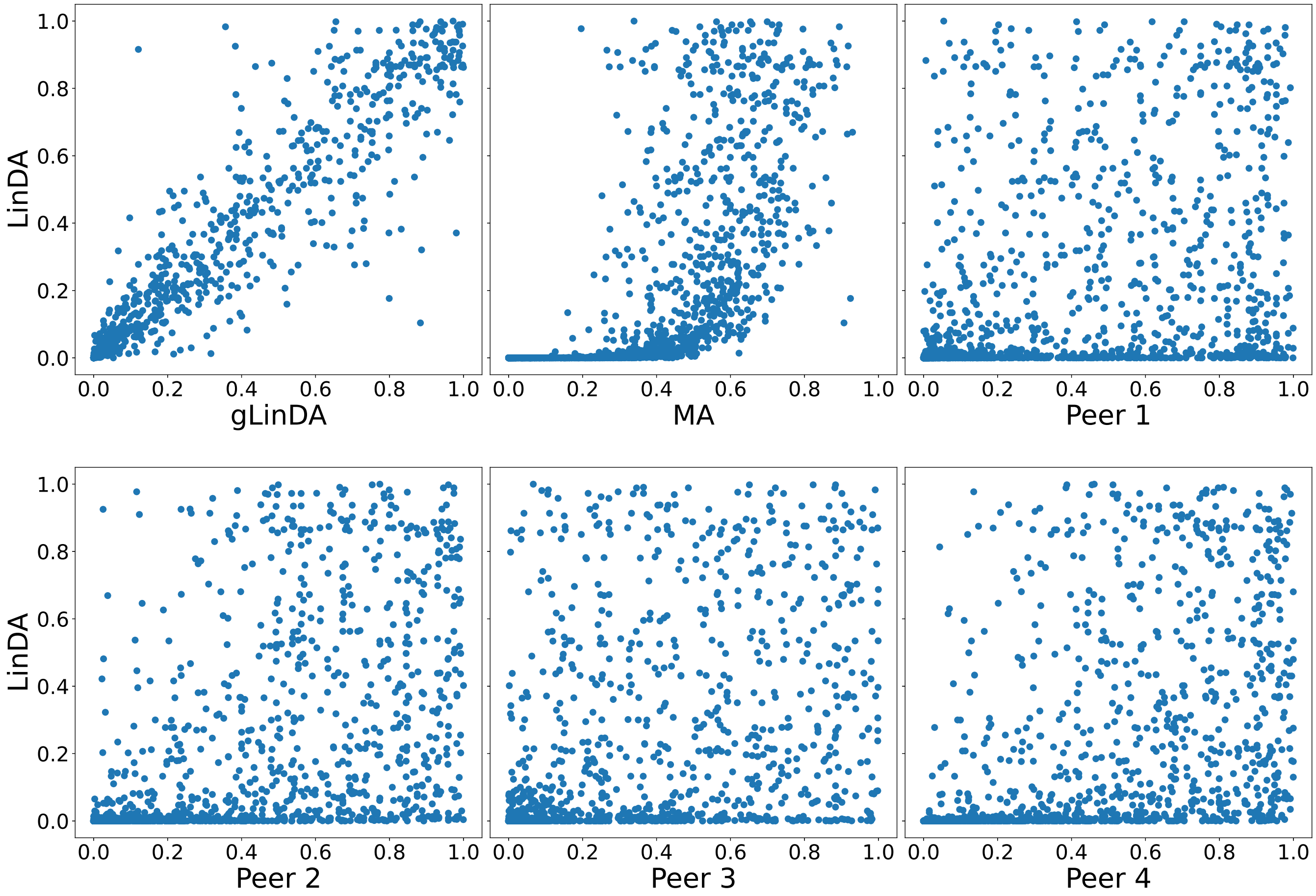


 Leon Fehse, Mohammad Tajabadi, Roman Martin, Hajo Holzmann, Dominik Heider

**Fig. S39.** P-values for the GWMC-Hot-Cold dataset for four peers with equal data distribution with the intersection approach.

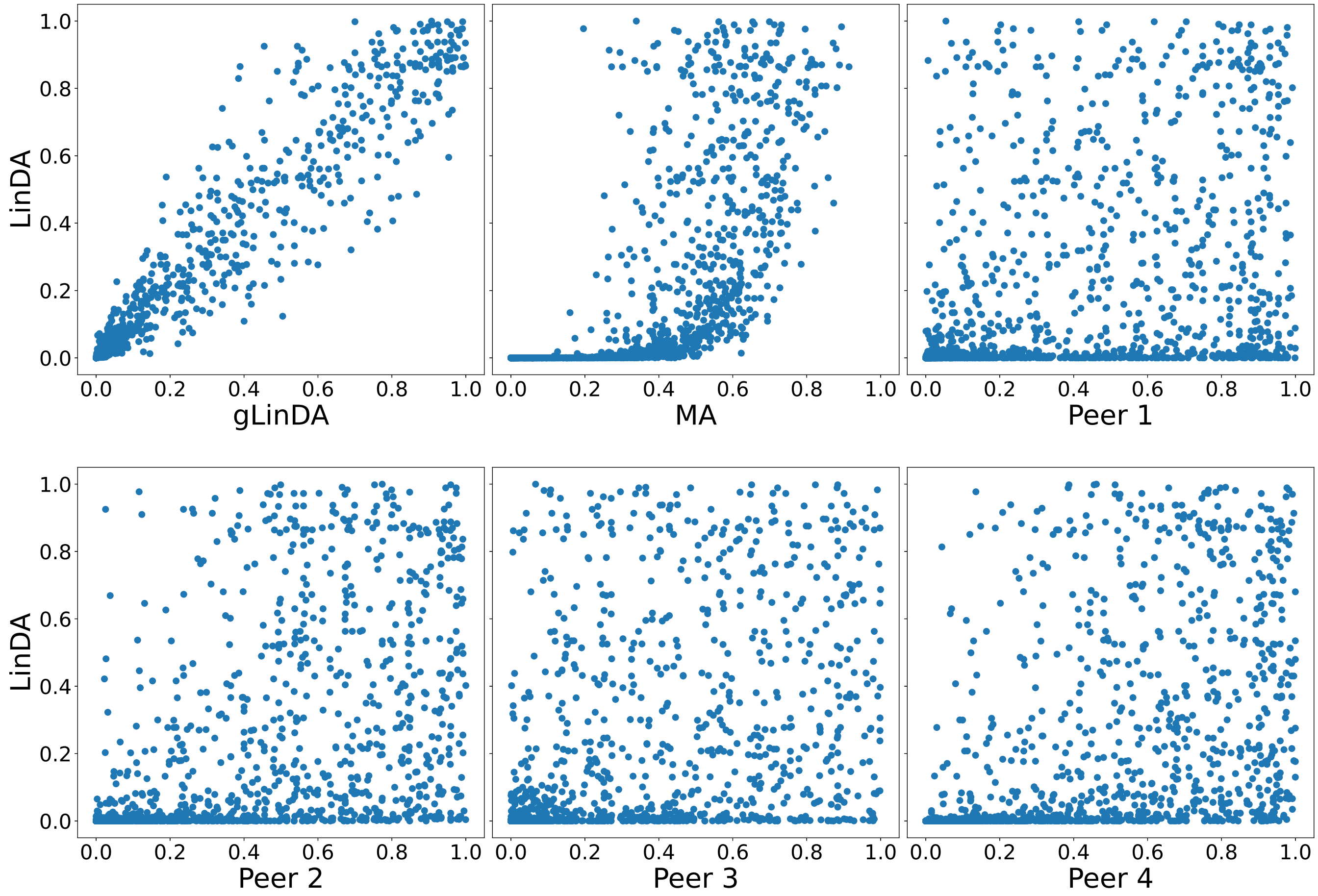

**Fig. S40.** P-values for the Arctic-Freshwater dataset for four peers with equal data distribution with the union approach.

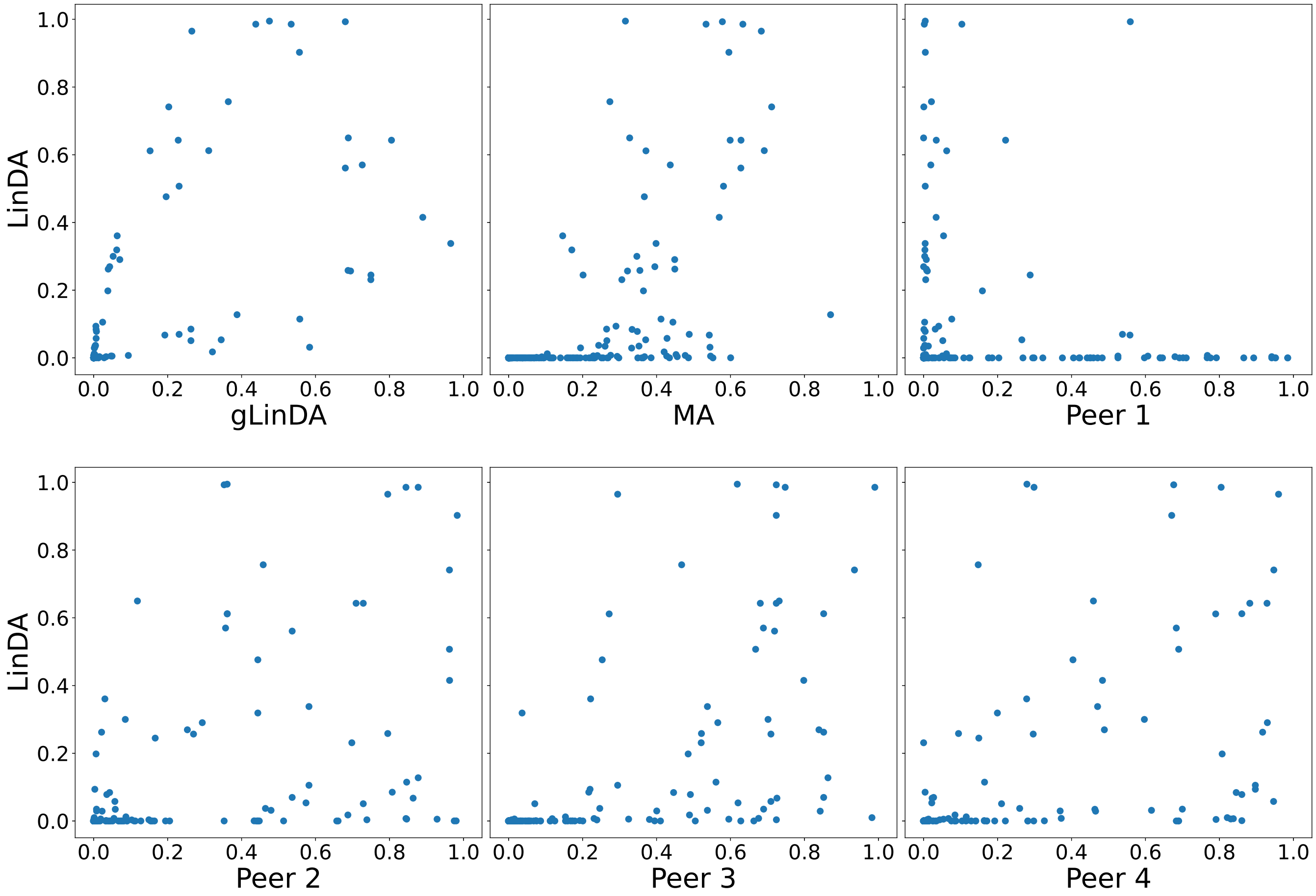

**Fig. S41.** P-values for the Arctic-Freshwater dataset for four peers with equal data distribution with the intersection approach.

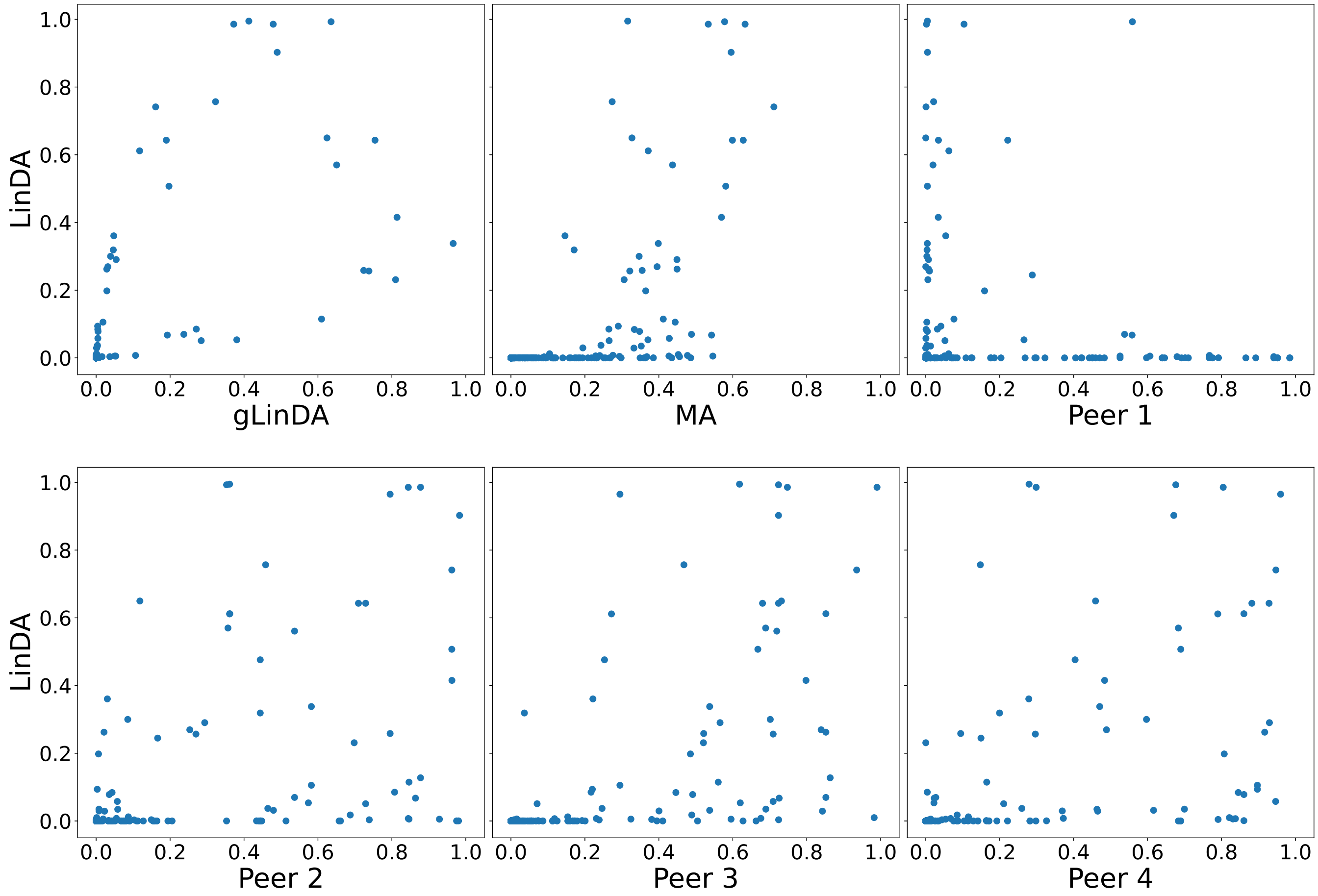

**Fig. S42.** P-values for the CDI-Shubert dataset for four peers with equal data distribution with the union approach.

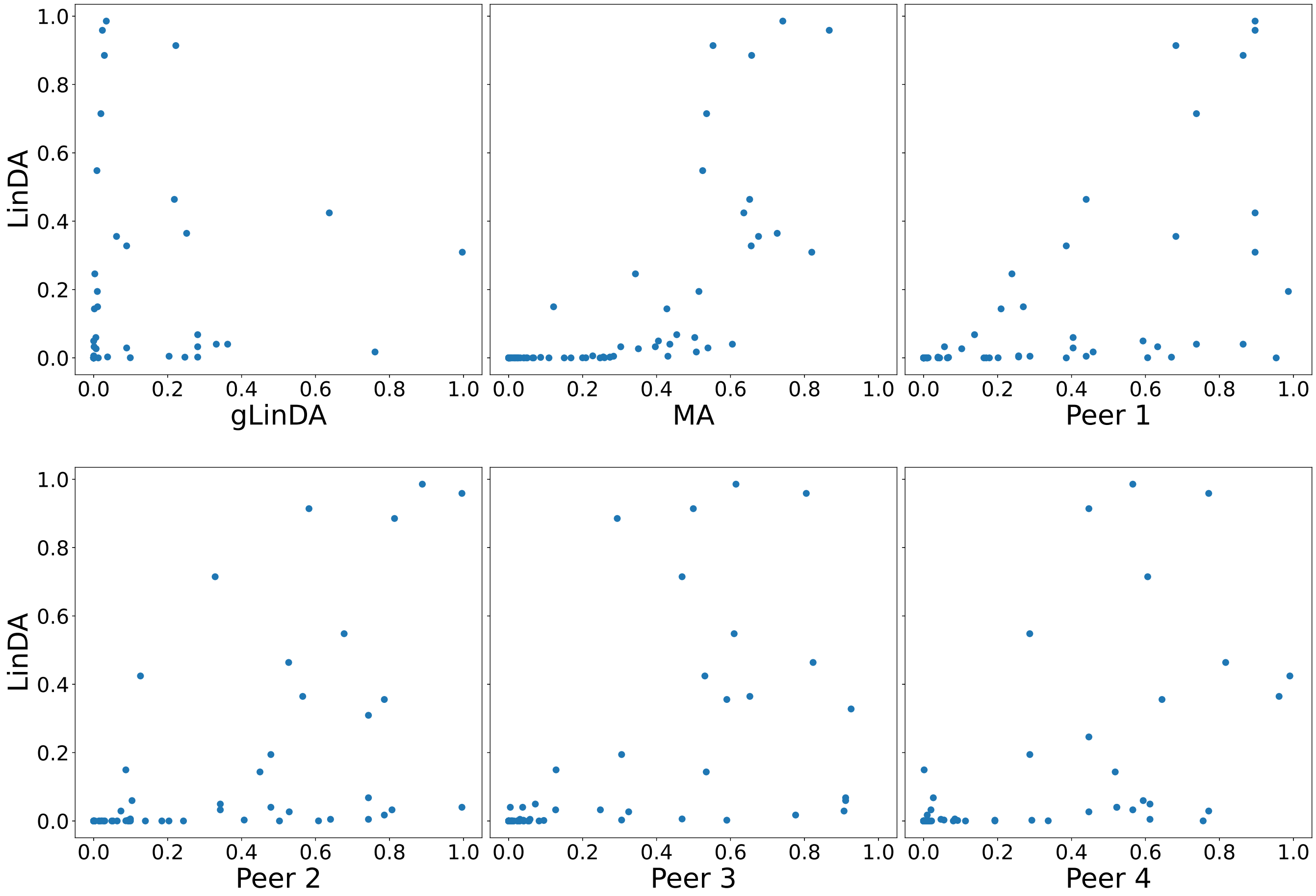

**Fig. S43.** P-values for the CDI-Shubert dataset for four peers with equal data distribution with the intersection approach.

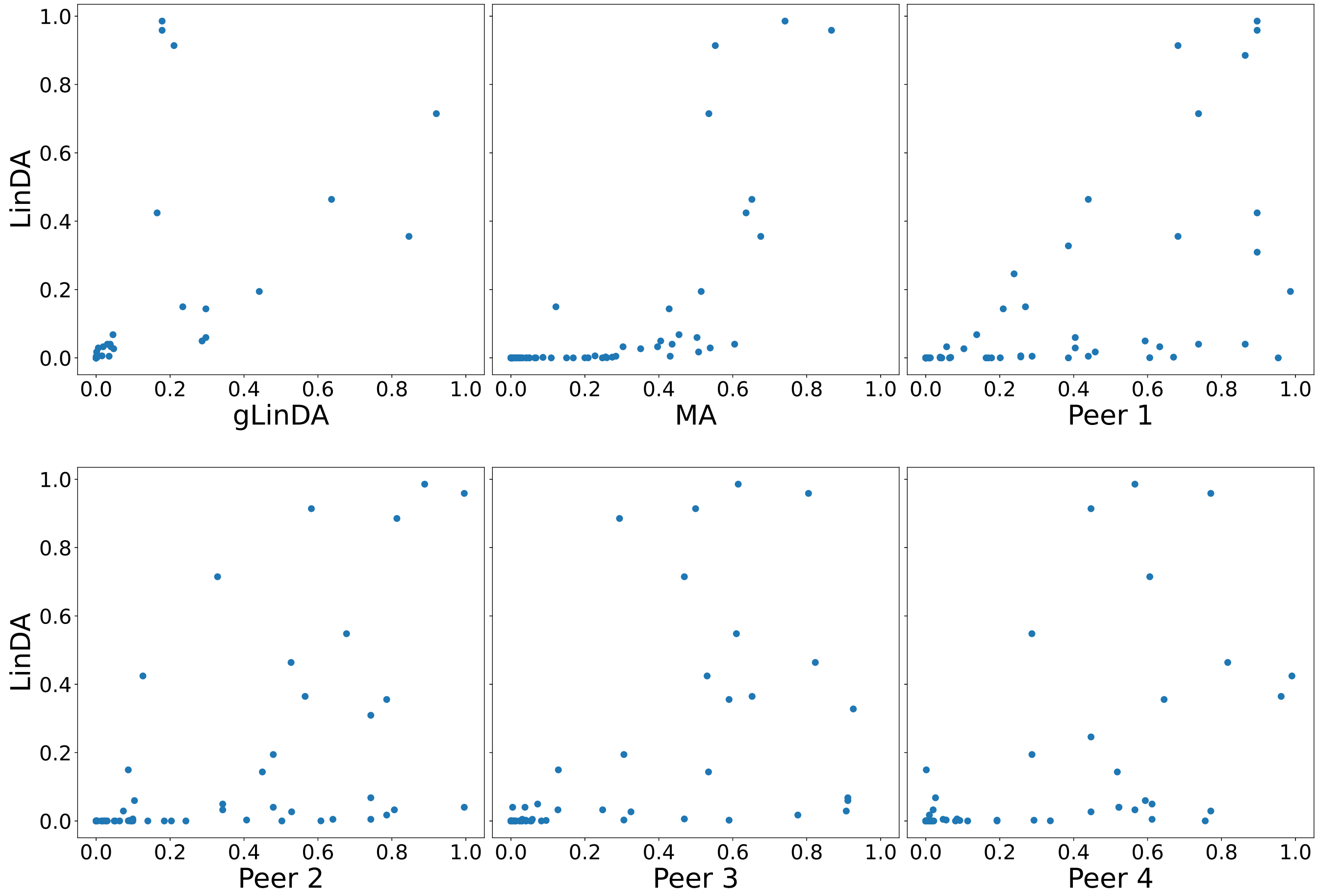

**Fig. S44.** P-values for the AGP-Diabetes dataset for four peers with equal data distribution with the union approach.

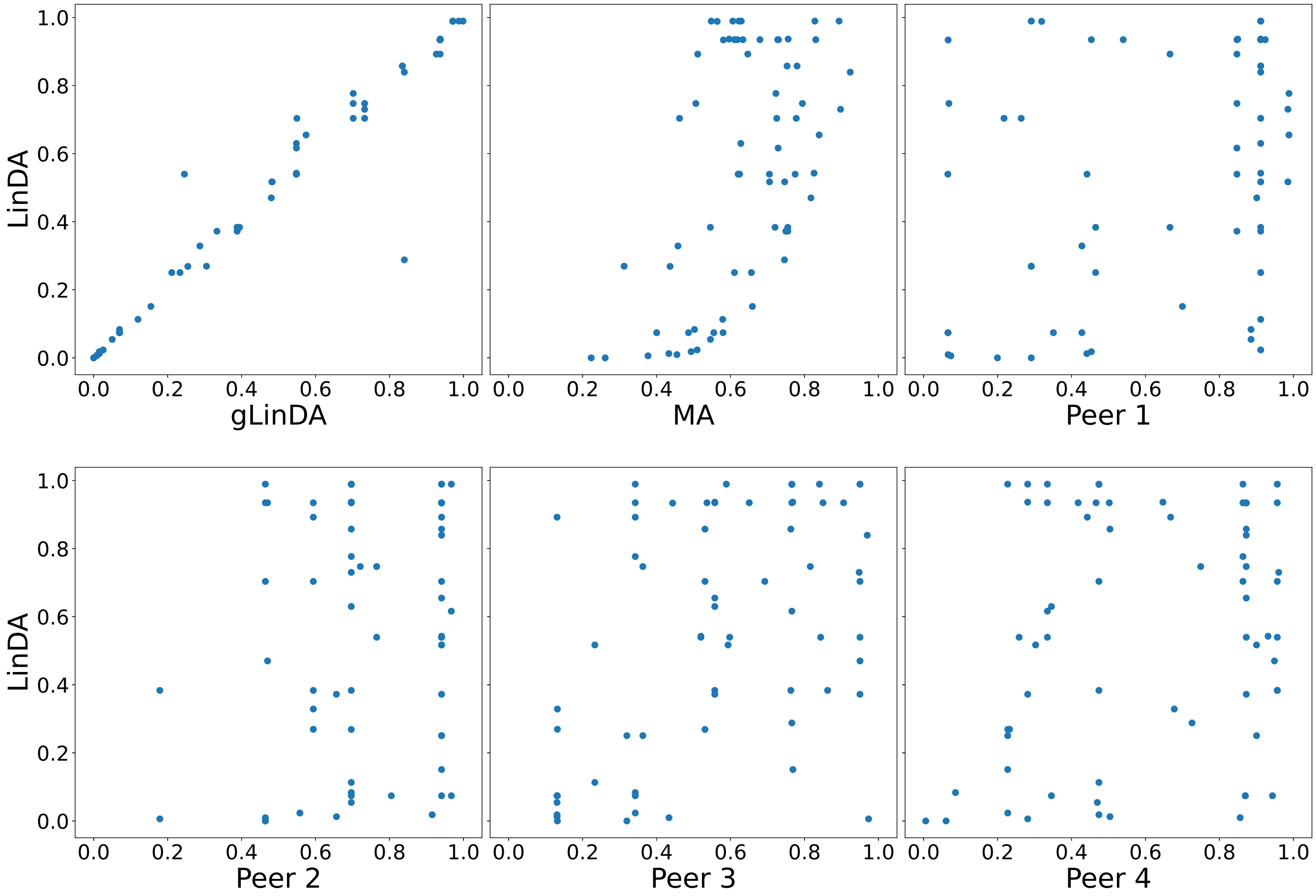

**Fig. S45.** P-values for the AGP-Diabetes dataset for four peers with equal data distribution with the intersection approach.

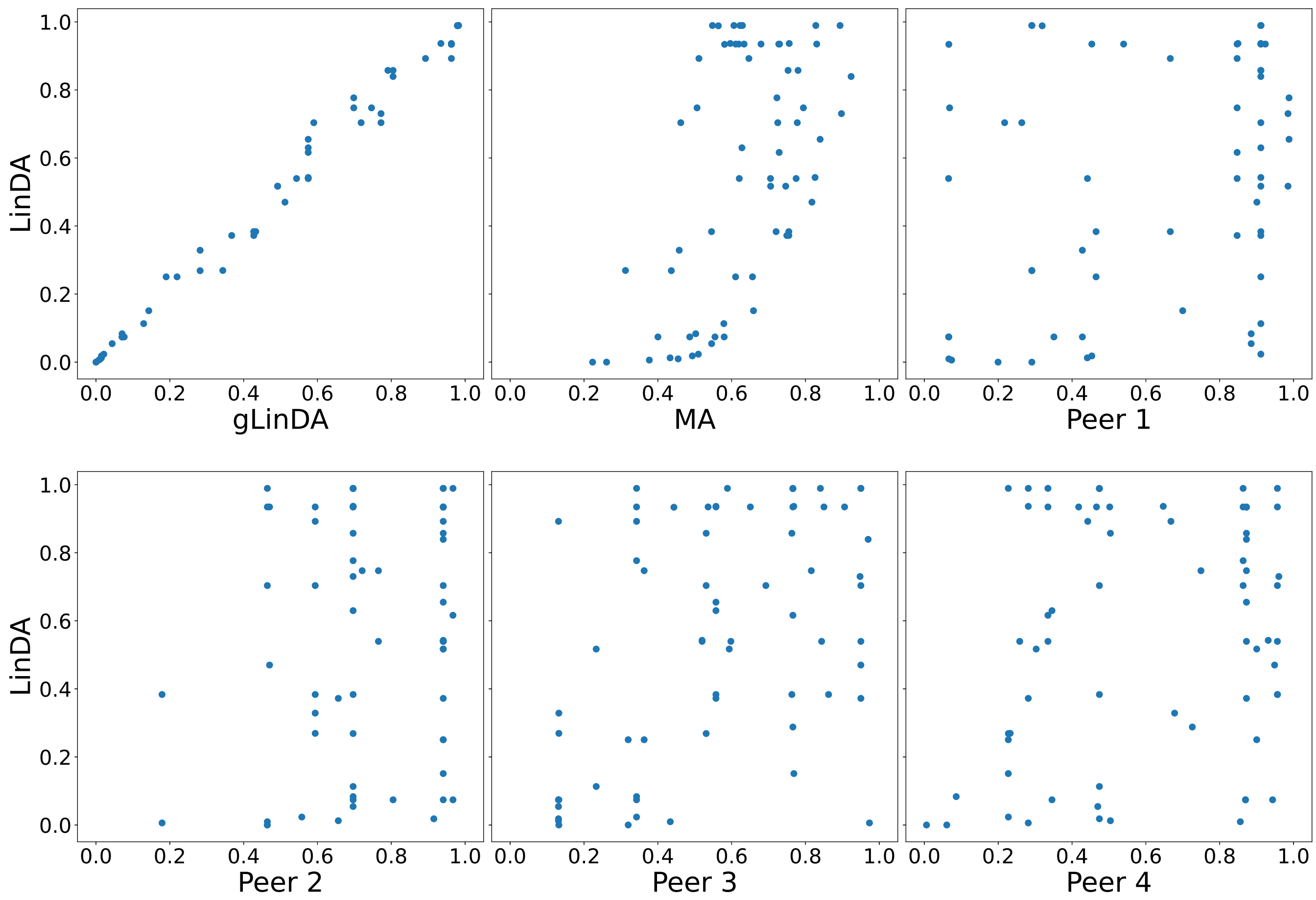

**Fig. S46.** P-values for the AGP-Ibs dataset for four peers with equal data distribution with the union approach.

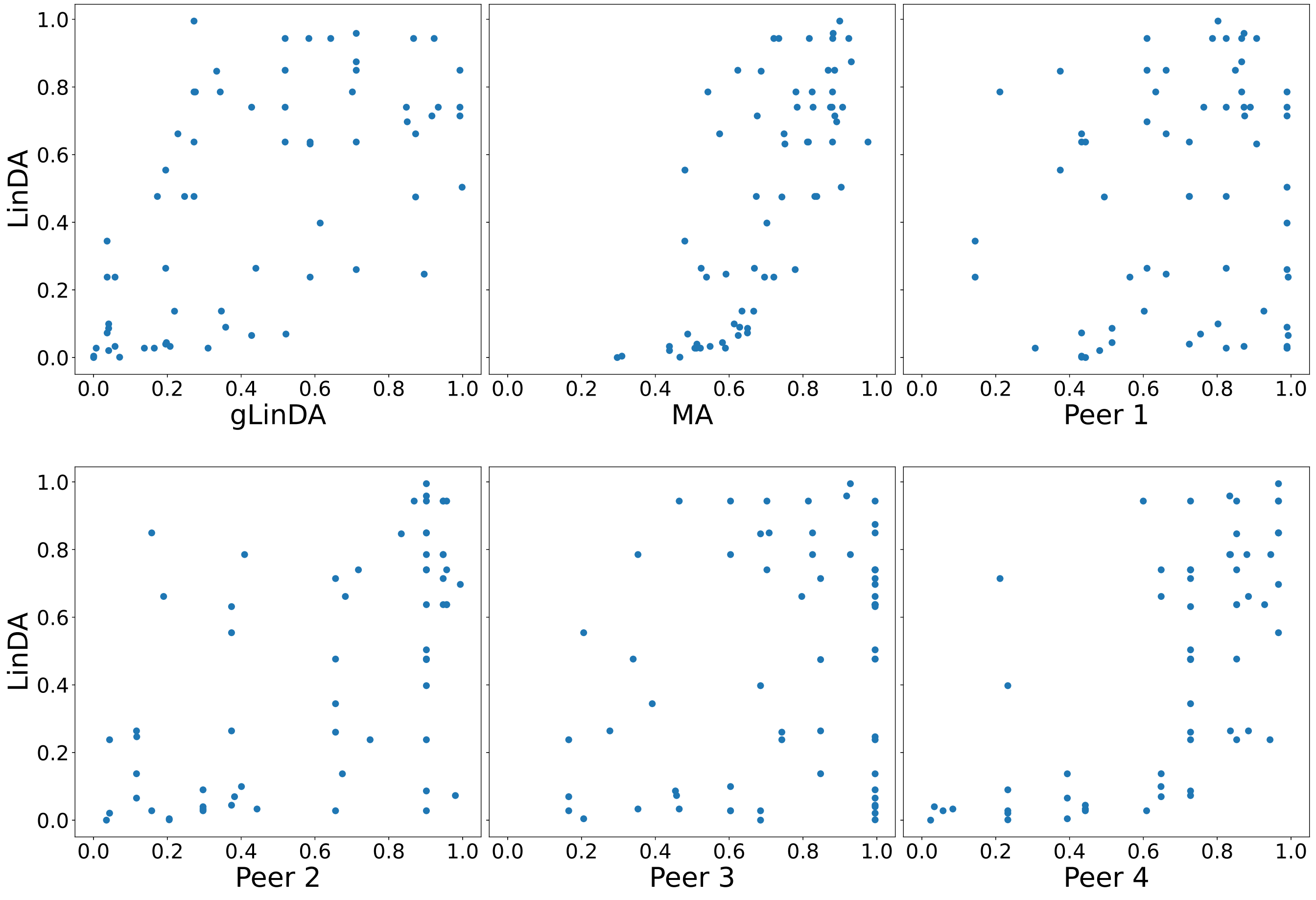

**Fig. S47.** P-values for the AGP-Ibs dataset for four peers with equal data distribution with the intersection approach.

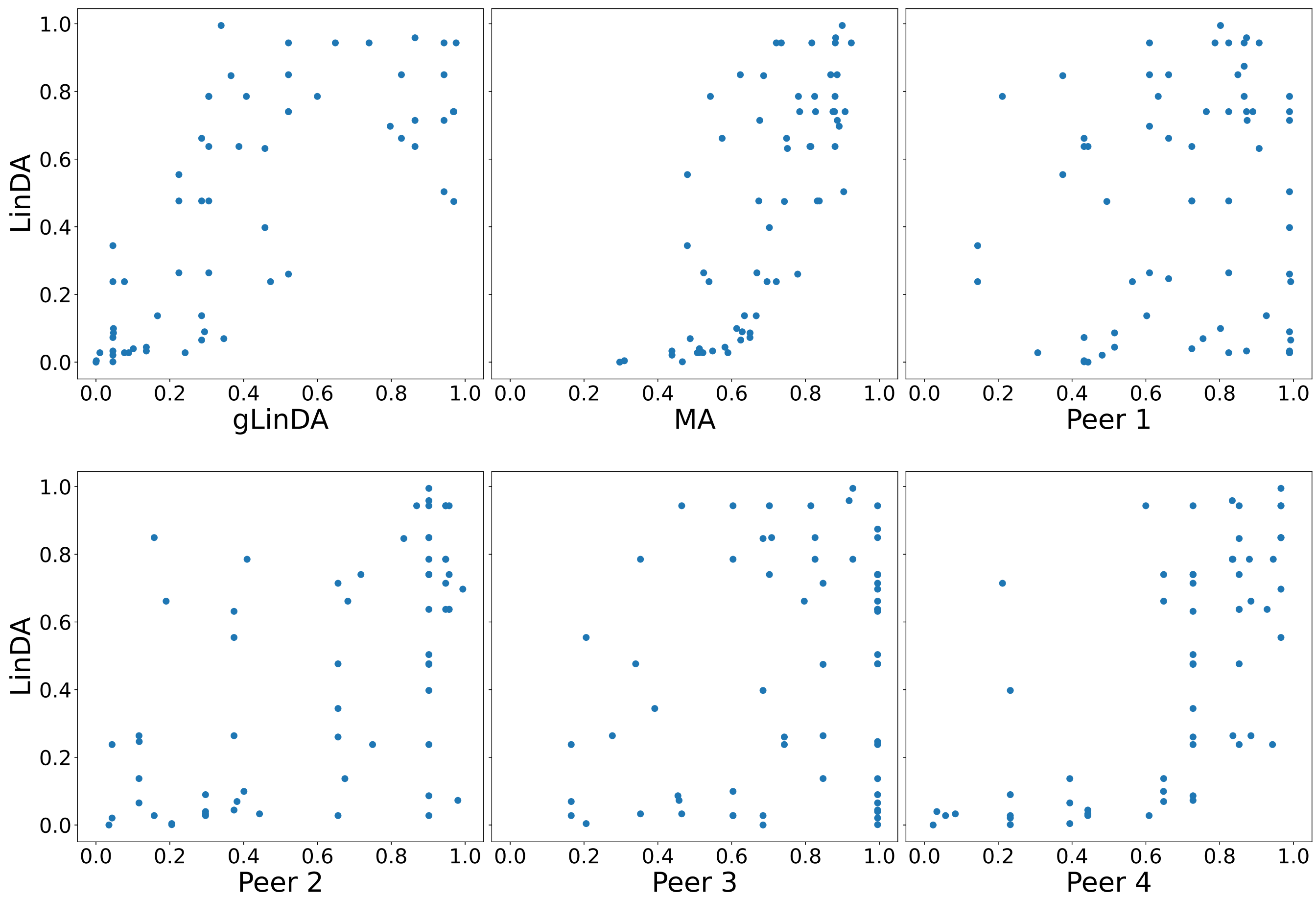

**Fig. S48.** P-values for the Smoke dataset for four peers with equal data distribution with the union approach.

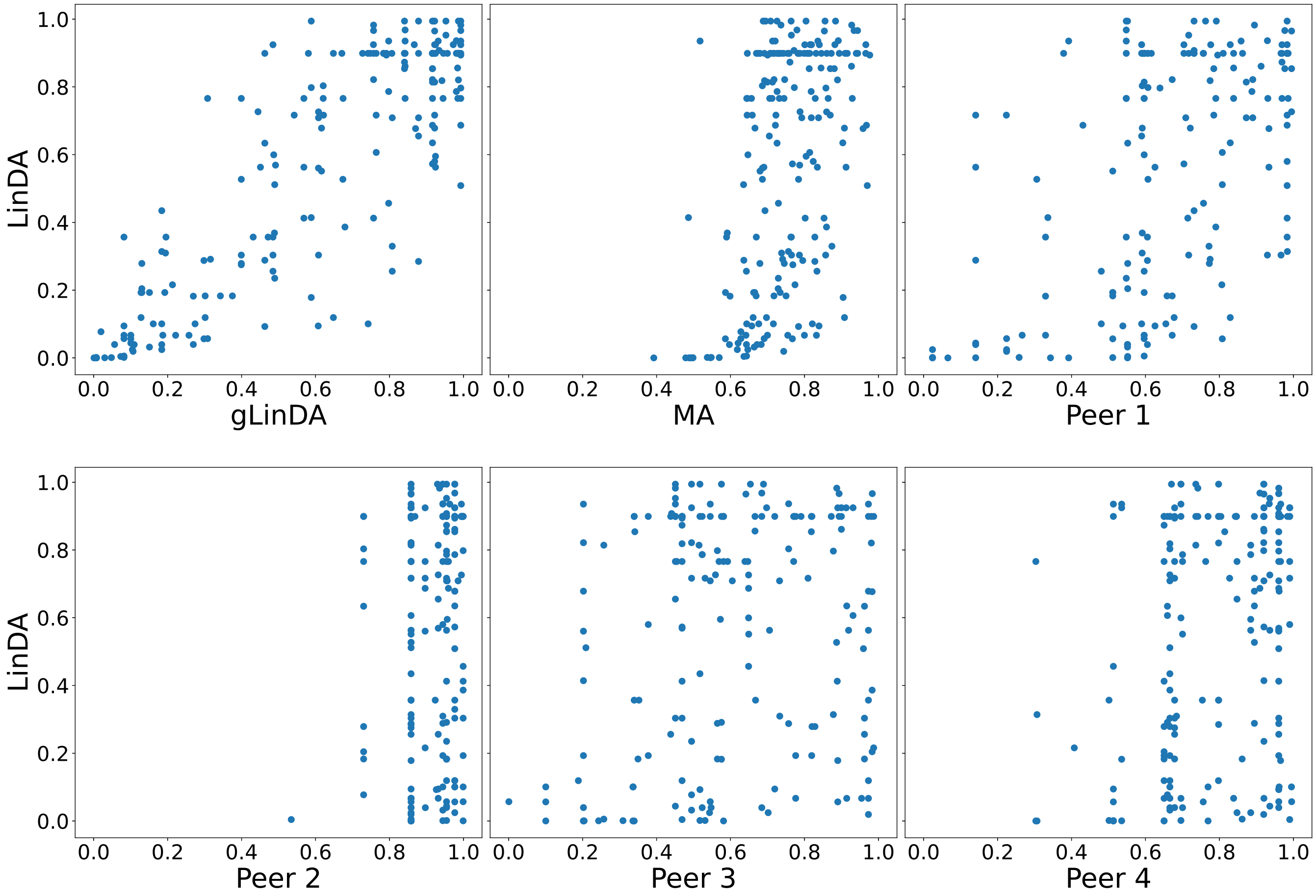

**Fig. S49.** P-values for the Smoke dataset for four peers with equal data distribution with the intersection approach.

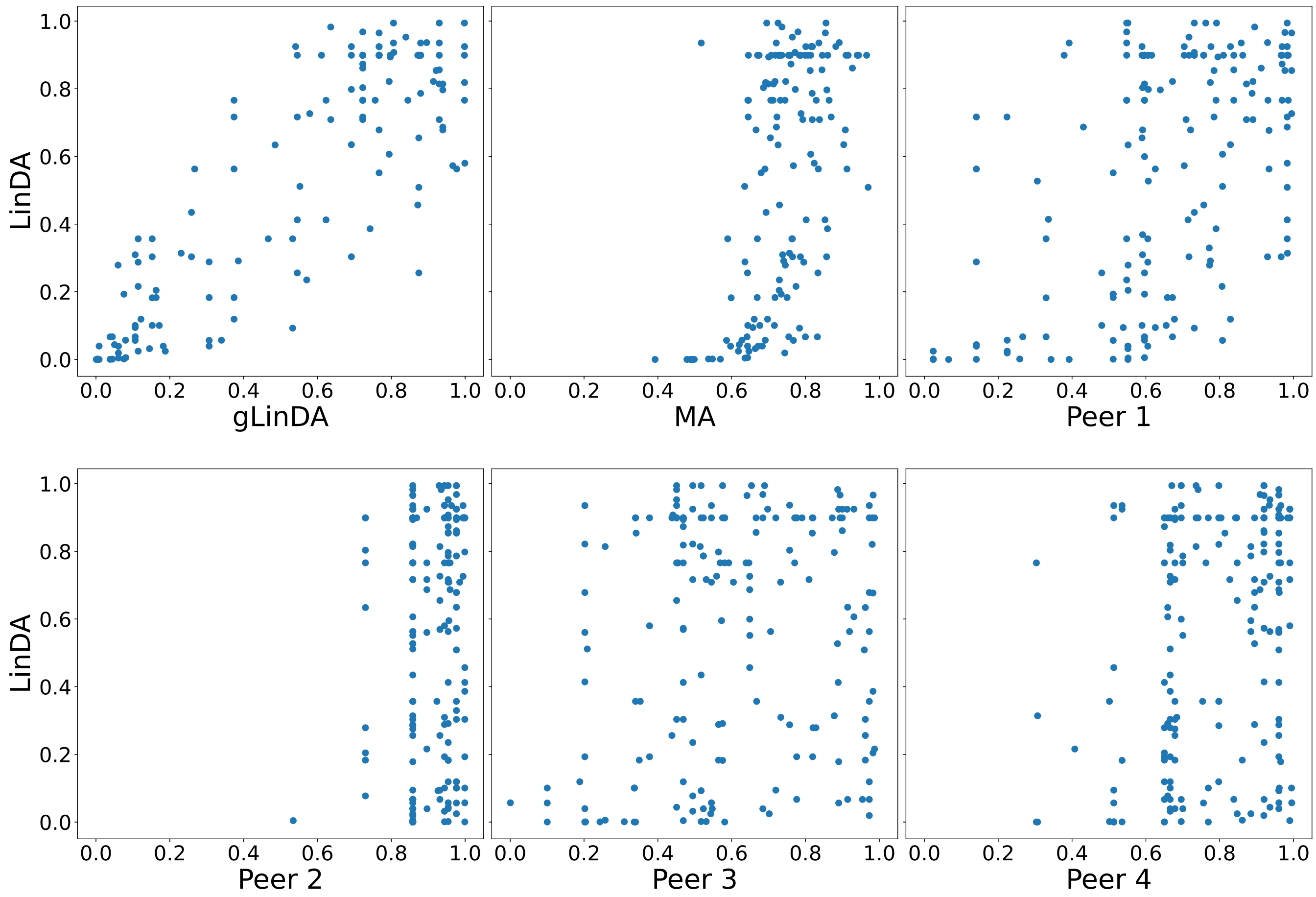

**Fig. S50.** P-values for the S-5000 dataset for eight peers with equal data distribution with the union approach.

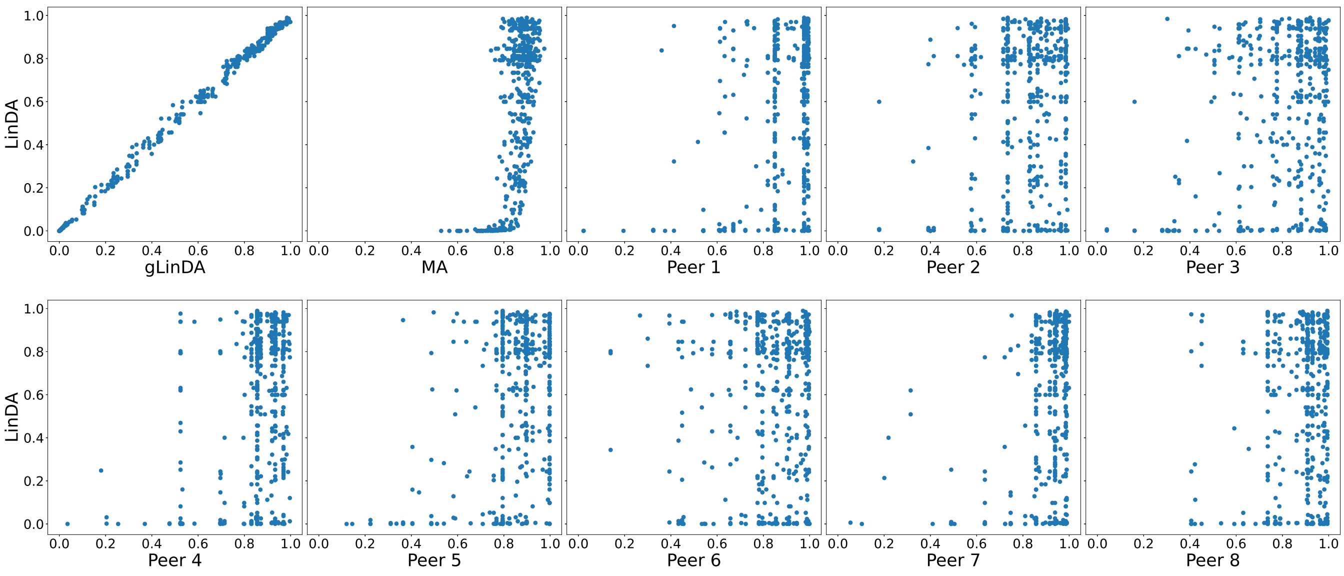


 Leon Fehse, Mohammad Tajabadi, Roman Martin, Hajo Holzmann, Dominik Heider

**Fig. S51.** P-values for the S-5000 dataset for eight peers with equal data distribution with the intersection approach.

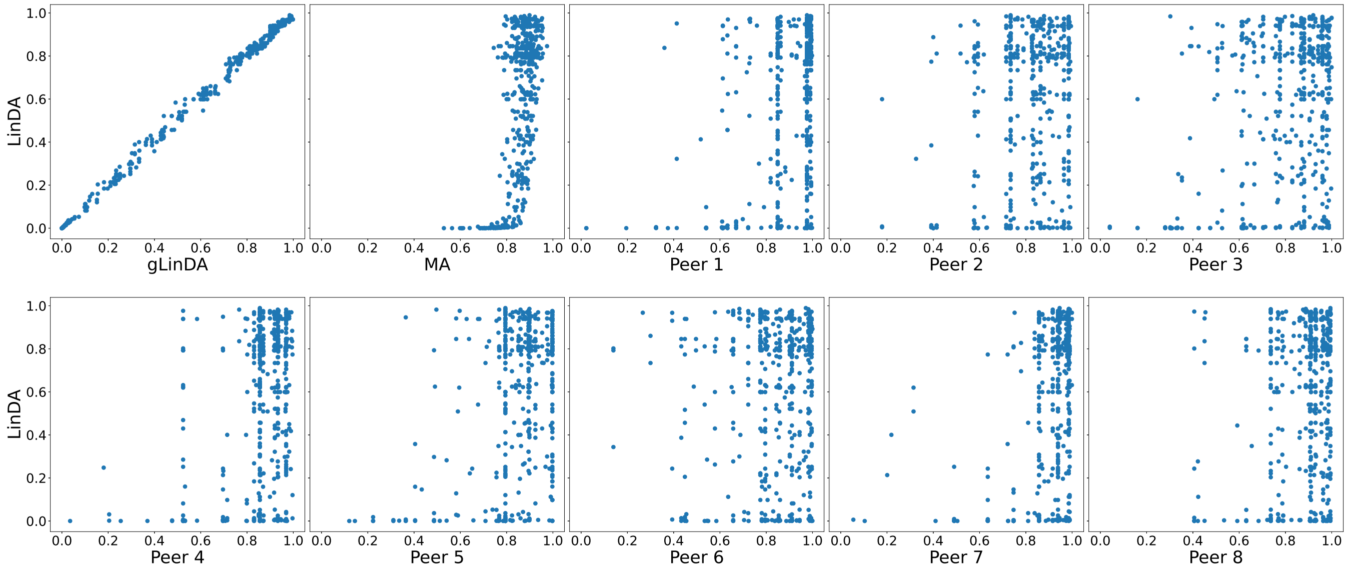

**Fig. S52.** P-values for the OB-Goodrich dataset for eight peers with equal data distribution with the union approach.

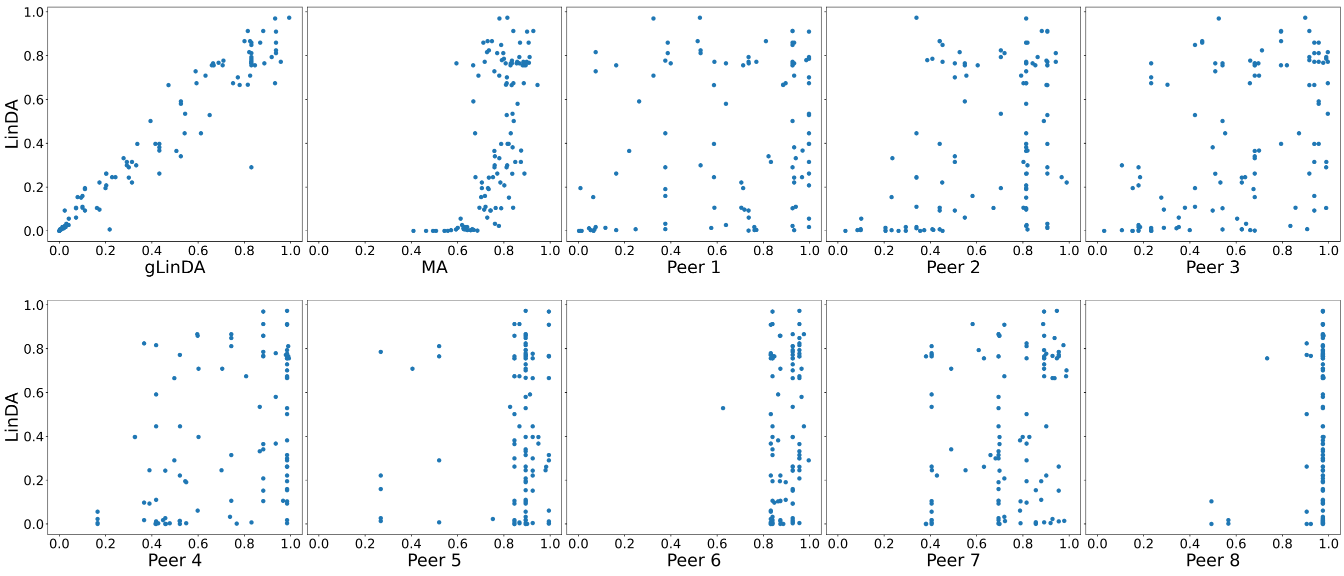

**Fig. S53.** P-values for the OB-Goodrich dataset for eight peers with equal data distribution with the intersection approach.

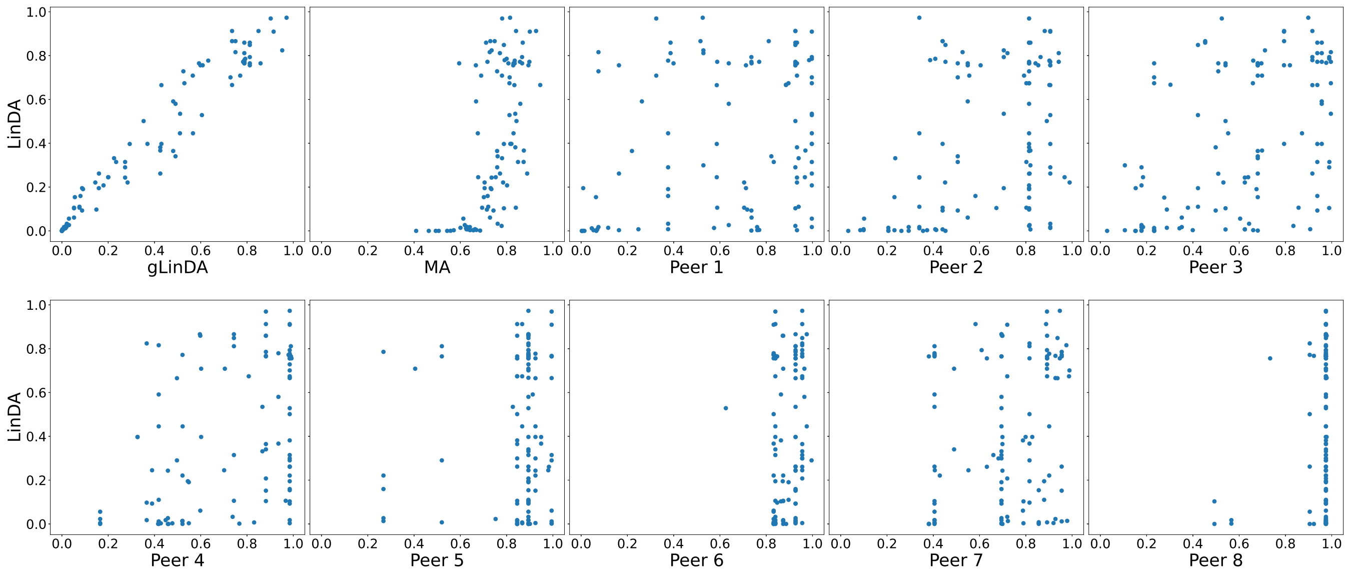

**Fig. S54.** P-values for the GWMC-Hot-Cold dataset for eight peers with equal data distribution with the union approach.

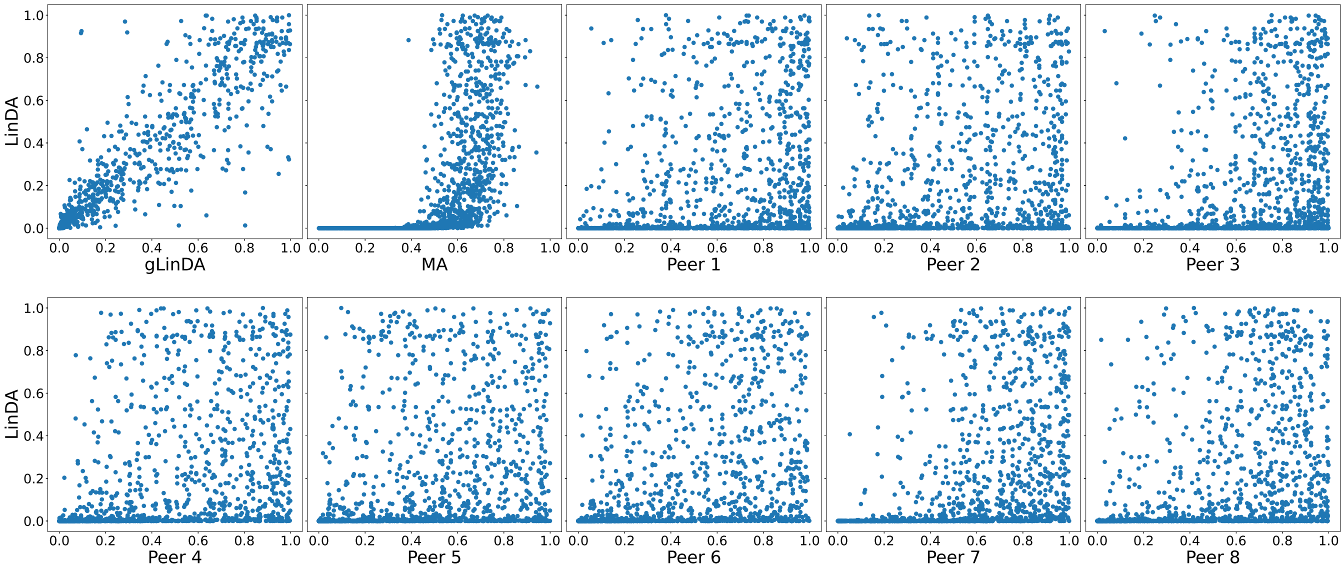

**Fig. S55.** P-values for the GWMC-Hot-Cold dataset for eight peers with equal data distribution with the intersection approach.

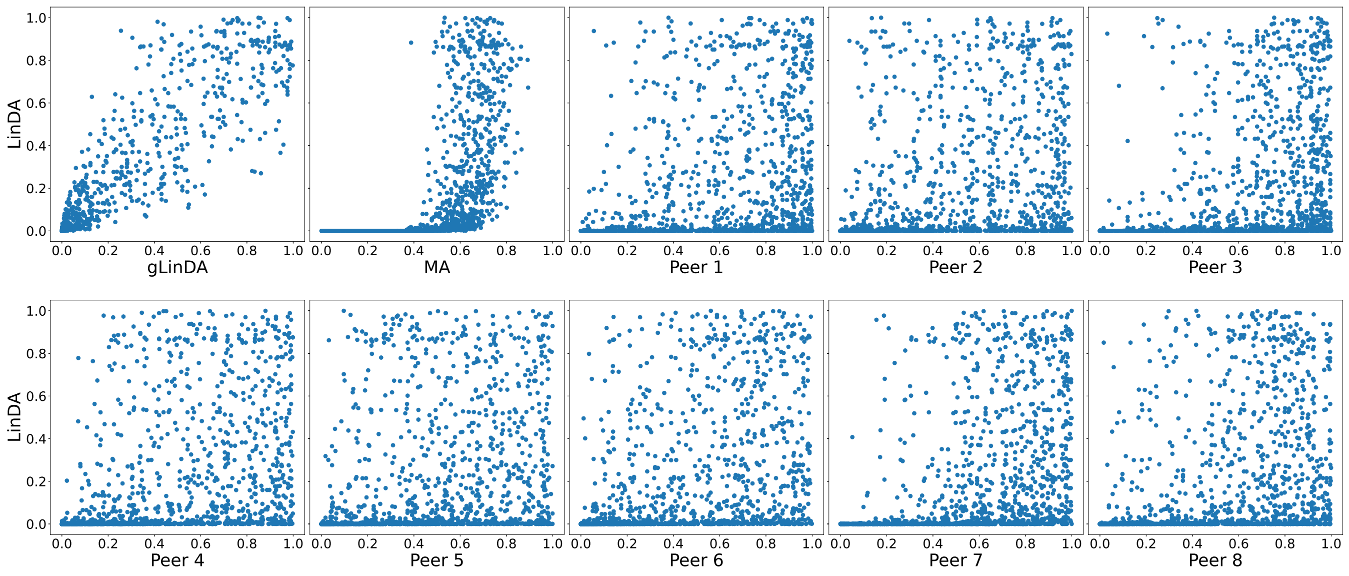

**Fig. S56.** P-values for the Arctic-Freshwater dataset for eight peers with equal data distribution with the union approach.

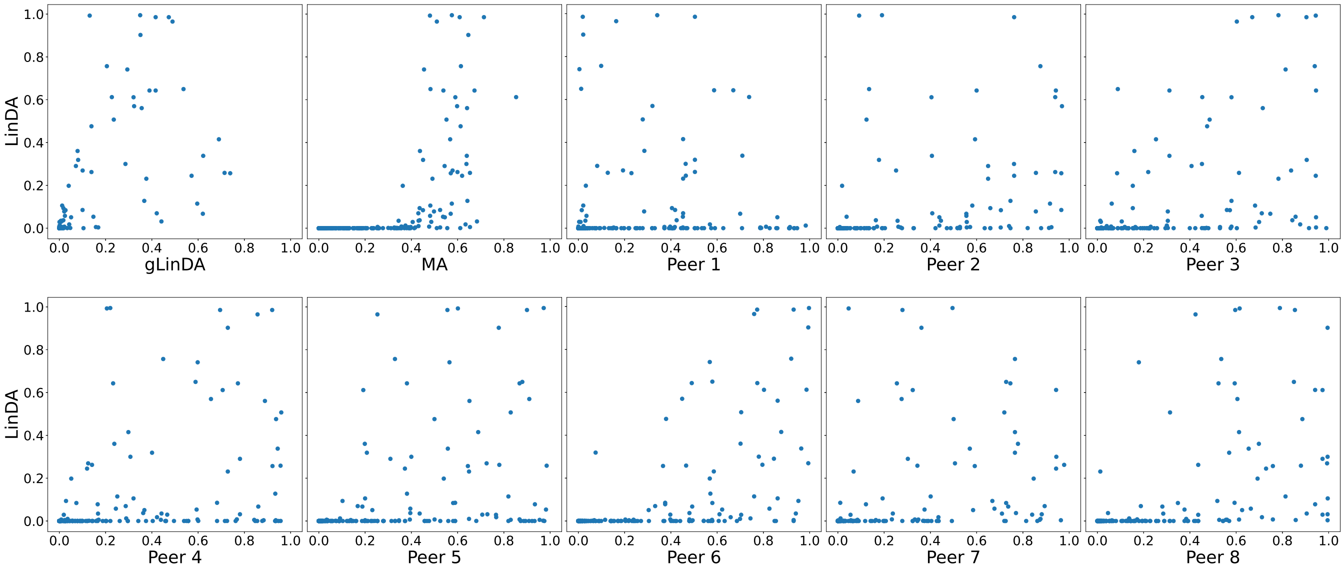

**Fig. S57.** P-values for the Arctic-Freshwater dataset for eight peers with equal data distribution with the intersection approach.

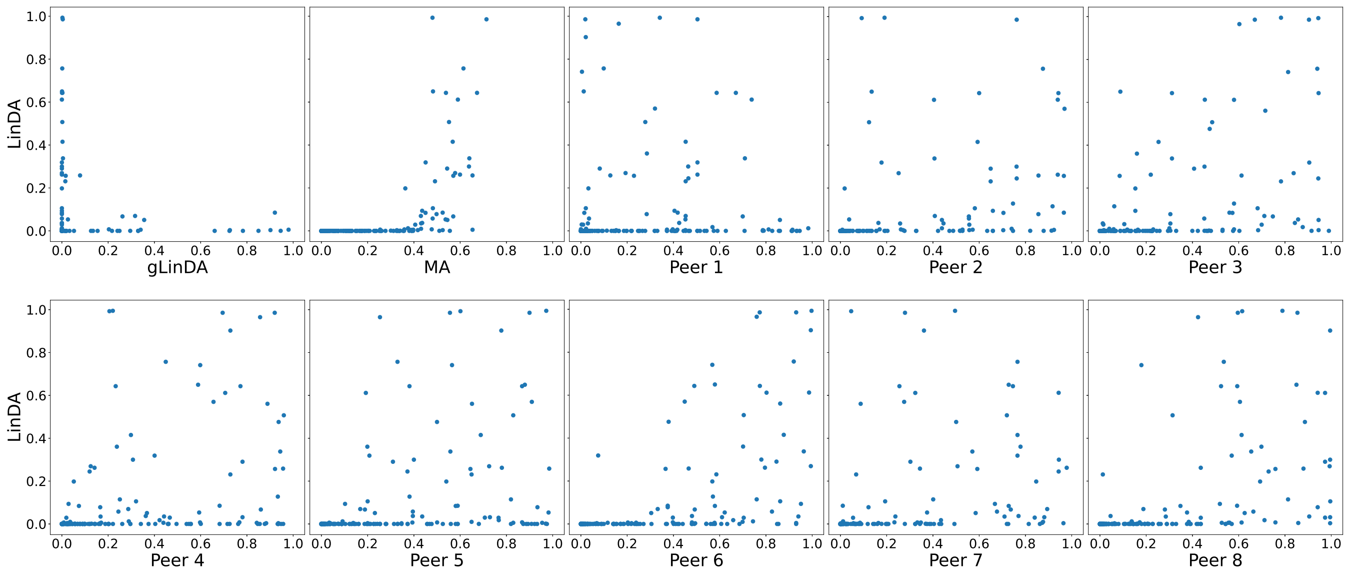

**Fig. S58.** P-values for the CDI-Shubert dataset for eight peers with equal data distribution with the union approach.

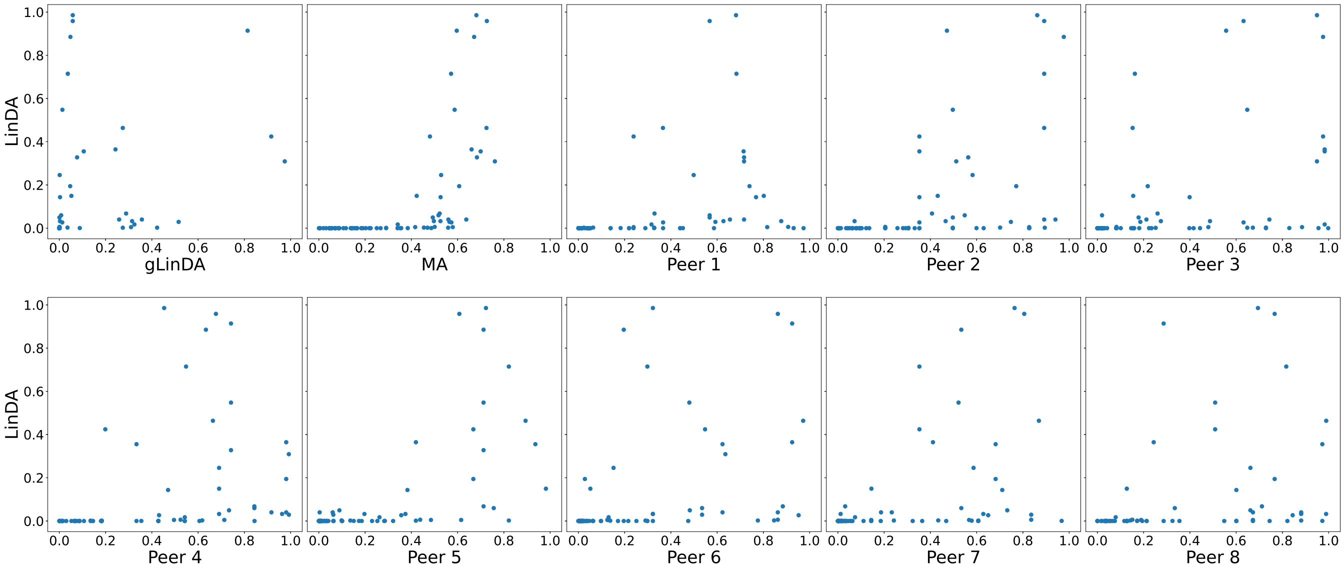

**Fig. S59.** P-values for the CDI-Shubert dataset for eight peers with equal data distribution with the intersection approach.

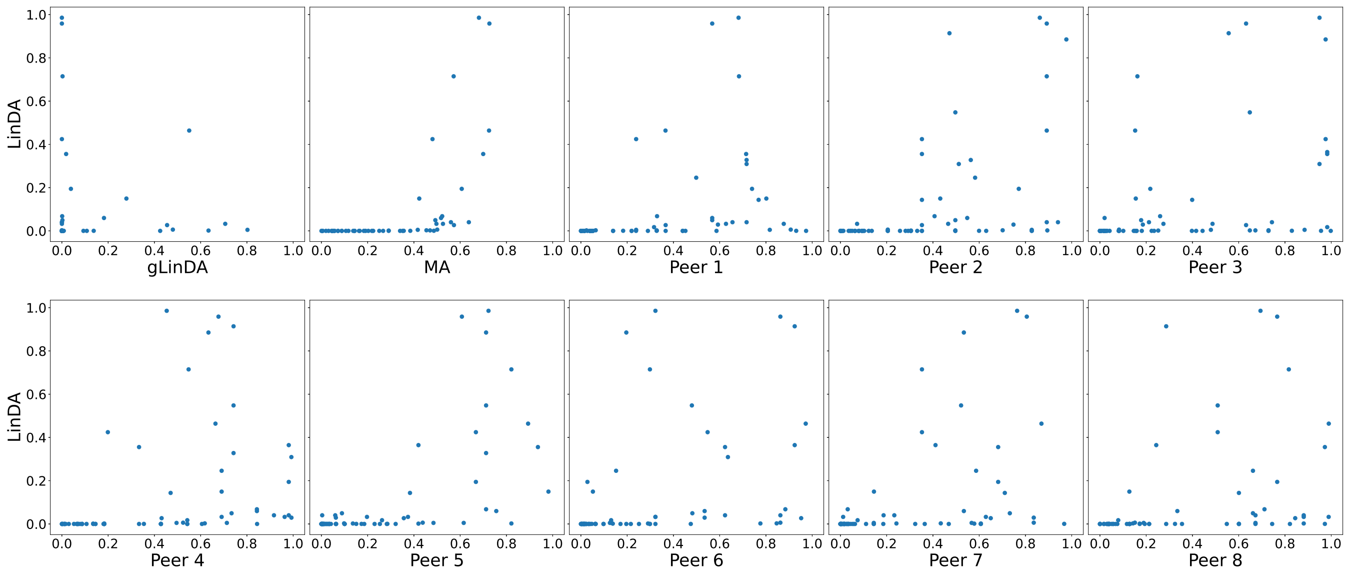

**Fig. S60.** P-values for the AGP-Diabetes dataset for eight peers with equal data distribution with the union approach.

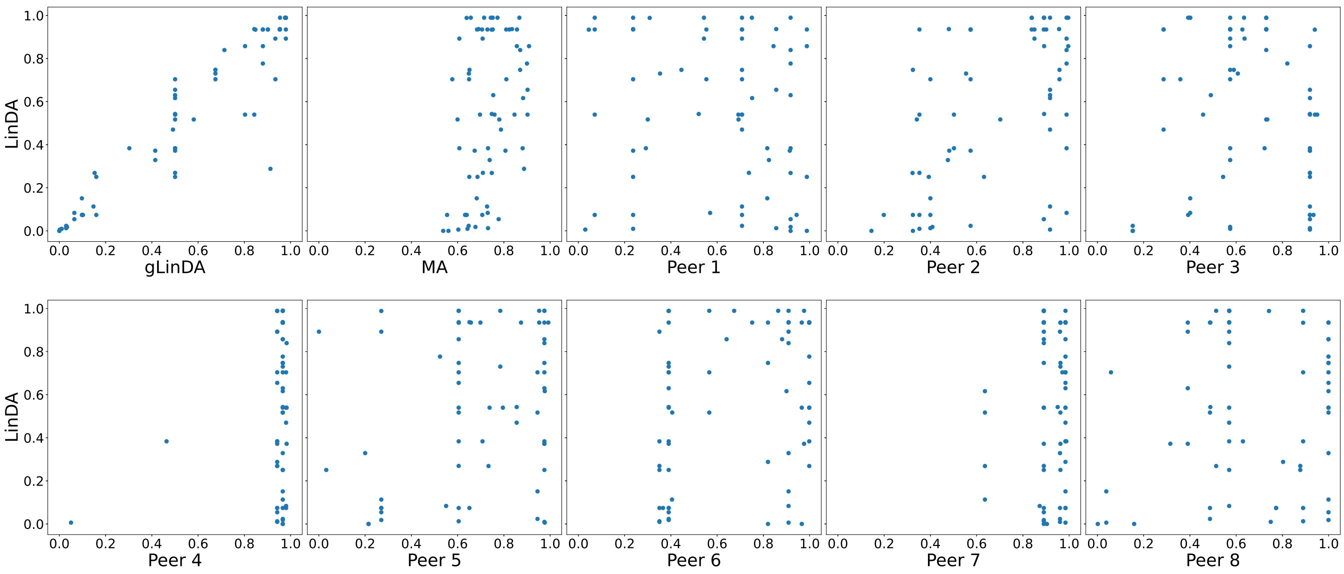

**Fig. S61.** P-values for the AGP-Diabetes dataset for eight peers with equal data distribution with the intersection approach.

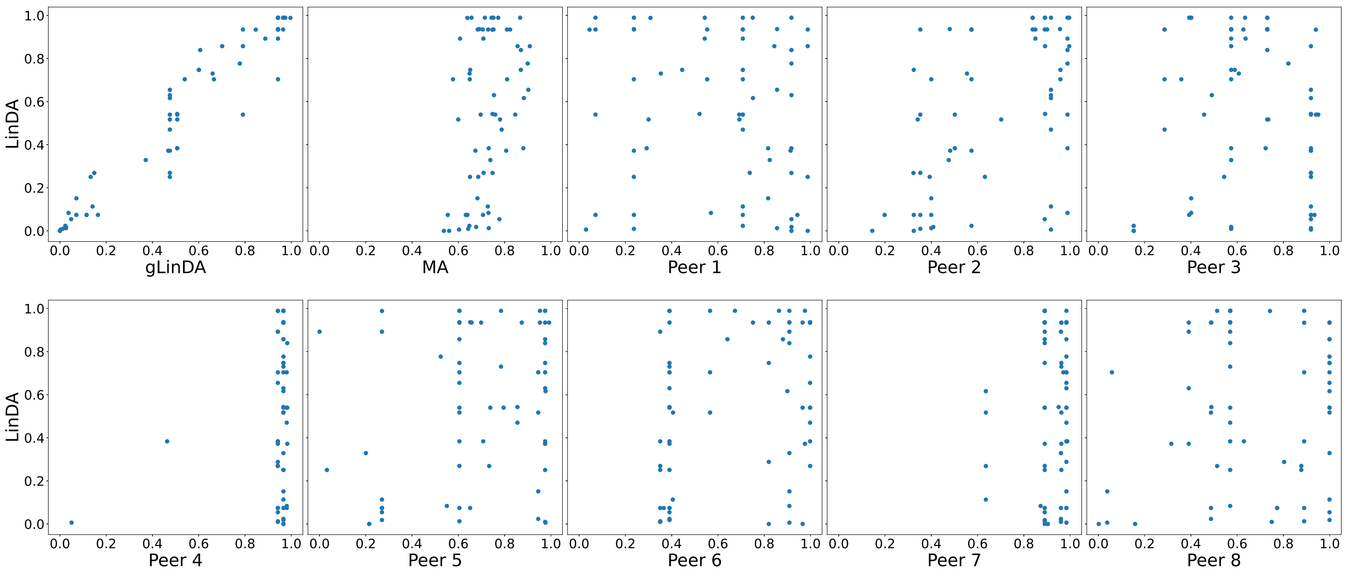

**Fig. S62.** P-values for the AGP-lbs dataset for eight peers with equal data distribution with the union approach.

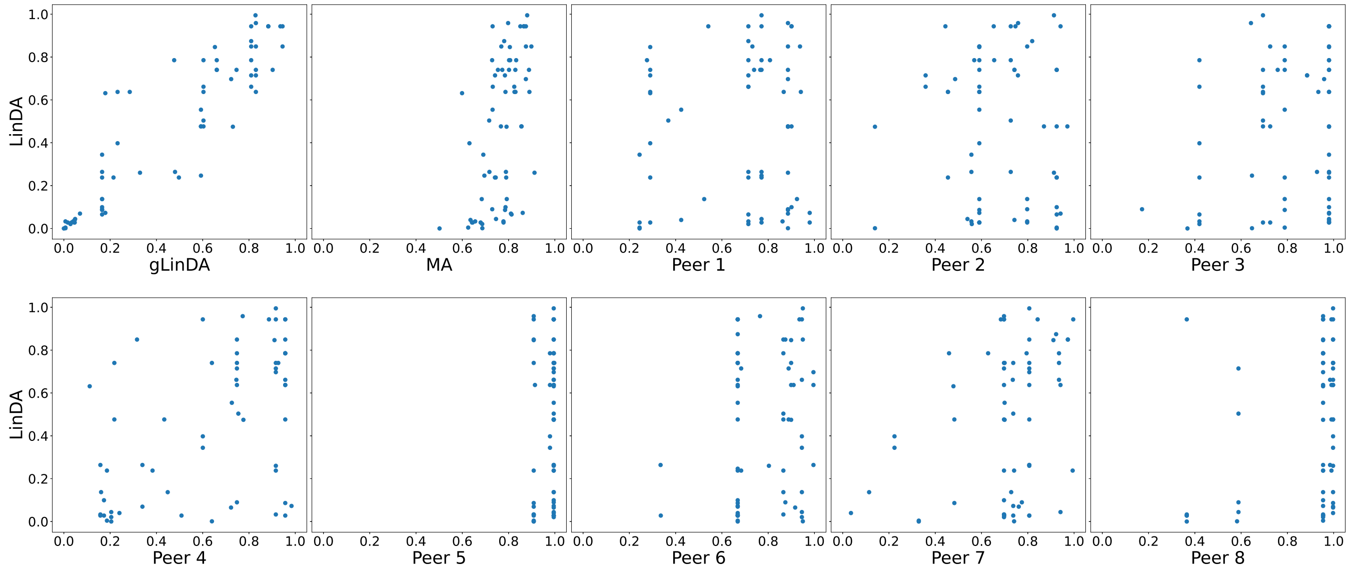


**Fig. S63.** P-values for the AGP-lbs dataset for eight peers with equal data distribution with the intersection approach.

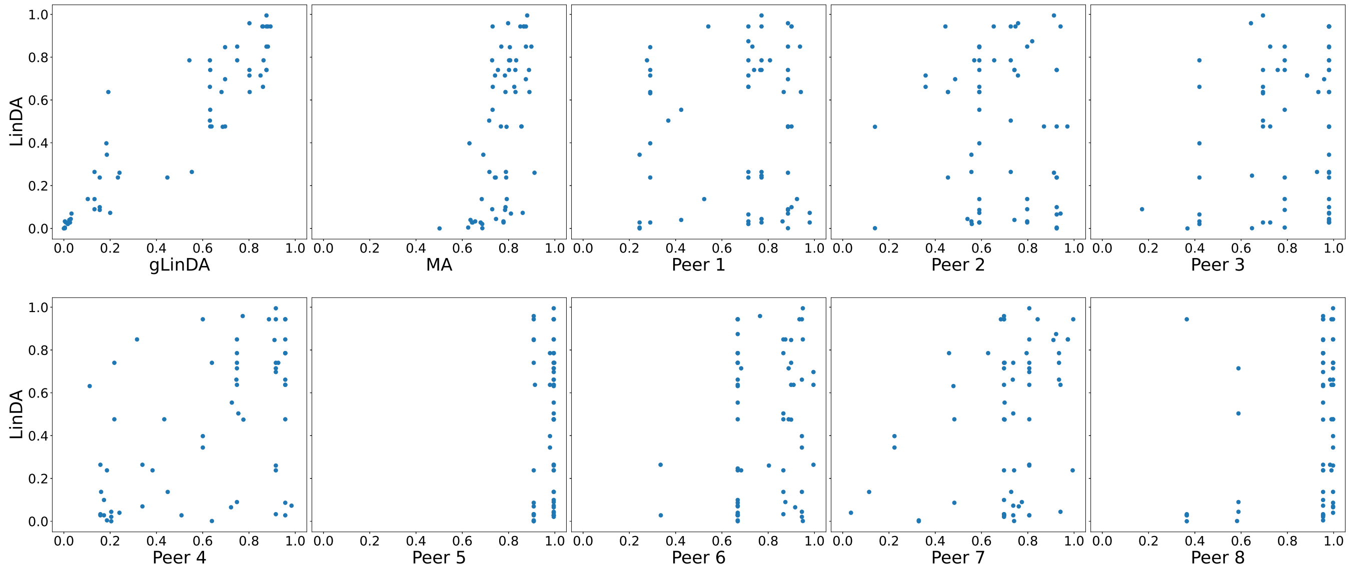

**Fig. S64.** P-values for the Smoke dataset for eight peers with equal data distribution with the union approach.

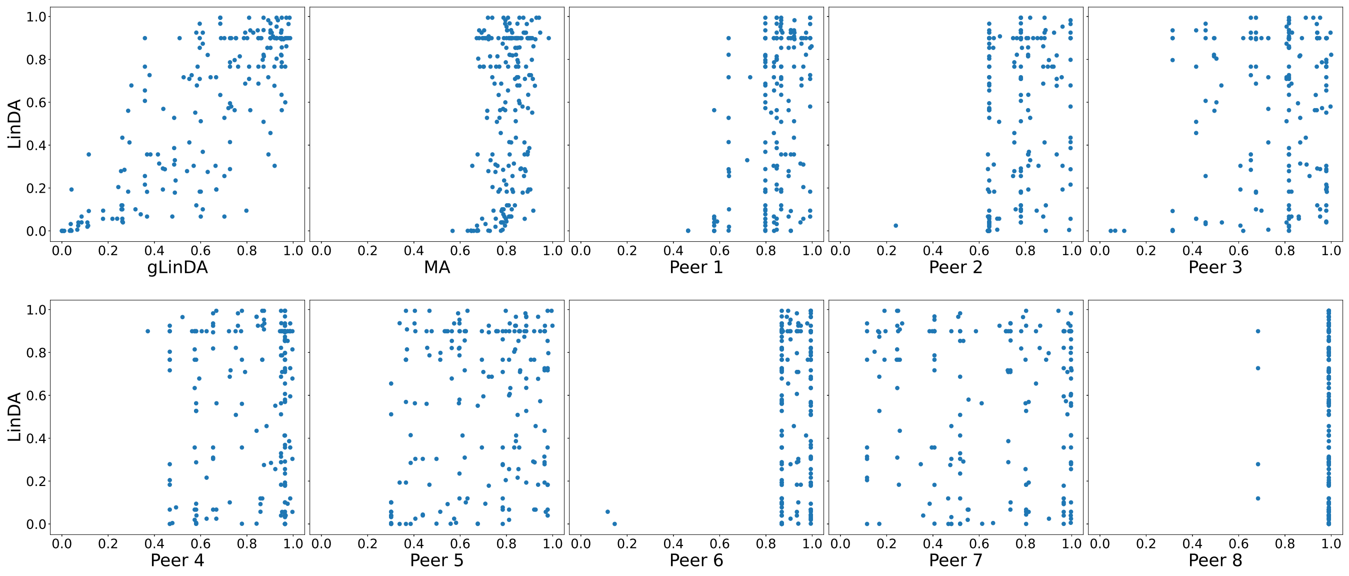

**Fig. S65.** P-values for the Smoke dataset for eight peers with equal data distribution with the intersection approach.

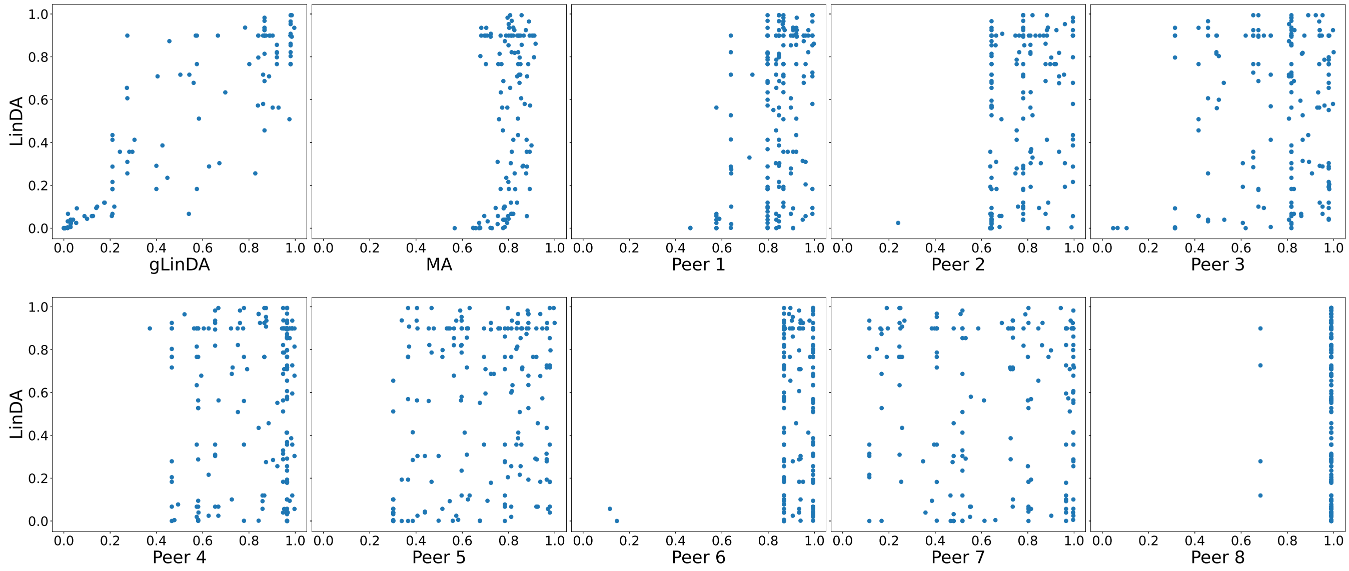


 
